# DNA repair and anti-cancer mechanisms in the long-lived bowhead whale

**DOI:** 10.1101/2023.05.07.539748

**Authors:** Denis Firsanov, Max Zacher, Xiao Tian, Todd L. Sformo, Yang Zhao, Greg Tombline, J. Yuyang Lu, Zhizhong Zheng, Luigi Perelli, Enrico Gurreri, Li Zhang, Jing Guo, Anatoly Korotkov, Valentin Volobaev, Seyed Ali Biashad, Zhihui Zhang, Johanna Heid, Alex Maslov, Shixiang Sun, Zhuoer Wu, Jonathan Gigas, Eric Hillpot, John Martinez, Minseon Lee, Alyssa Williams, Abbey Gilman, Nicholas Hamilton, Ena Haseljic, Avnee Patel, Maggie Straight, Nalani Miller, Julia Ablaeva, Lok Ming Tam, Chloé Couderc, Michael Hoopman, Robert Moritz, Shingo Fujii, Dan J. Hayman, Hongrui Liu, Yuxuan Cai, Anthony K. L. Leung, Mirre J. P. Simons, Zhengdong Zhang, C. Bradley Nelson, Lisa M. Abegglen, Joshua D. Schiffman, Vadim N. Gladyshev, Mauro Modesti, Giannicola Genovese, Jan Vijg, Andrei Seluanov, Vera Gorbunova

## Abstract

At over 200 years, the maximum lifespan of the bowhead whale exceeds that of all other mammals. The bowhead is also the second-largest animal on Earth, reaching over 80,000 kg^1^. Despite its very large number of cells and long lifespan, the bowhead is not highly cancer-prone, an incongruity termed Peto’s Paradox^2^. This phenomenon has been explained by the evolution of additional tumor suppressor genes in other larger animals, supported by research on elephants demonstrating expansion of the p53 gene^3–5^. Here we show that bowhead whale fibroblasts undergo oncogenic transformation after disruption of fewer tumor suppressors than required for human fibroblasts. However, analysis of DNA repair revealed that bowhead cells repair double strand breaks (DSBs) and mismatches with uniquely high efficiency and accuracy compared to other mammals. The protein CIRBP, implicated in protection from genotoxic stress, was present in very high abundance in the bowhead whale relative to other mammals. We show that CIRBP and its downstream protein RPA2, also present at high levels in bowhead cells, increase the efficiency and fidelity of DNA repair in human cells. These results indicate that rather than possessing additional tumor suppressor genes as barriers to oncogenesis, the bowhead whale relies on more accurate and efficient DNA repair to preserve genome integrity. This strategy which does not eliminate damaged cells but repairs them may be critical for the long and cancer-free lifespan of the bowhead whale.

## Introduction

The Alaskan Iñupiat Inuit, who carry on a long tradition of subsistence hunting of the bowhead whale (*Balaena mysticetus*), maintain that these animals “live two human lifetimes”^6^. A series of bowhead whales captured in the late-twentieth and early-twenty-first centuries lent new credence to these claims, as embedded in their bodies were traditional stone harpoon points and bomb lance fragments dating to the Victorian era^7^. Subsequent scientific study and age estimation through quantification of ovarian corpora, baleen dating, and eye lens aspartic acid racemization analysis supported a maximum lifespan exceeding 200 years in the bowhead whale^7–12^. Thus, the range of mammalian lifespans covers roughly 2 orders of magnitude, with the model organism *Mus musculus* living for 2-3 years while the bowhead whale lives 100 times as long.

The increased number of cells and cell divisions in larger organisms does not lead to increased cancer incidence and shorter lifespans^13^. The apparent contradiction between expected and observed cancer rates in relation to species body mass has been noted for decades and is known as Peto’s Paradox^2,14–16^. Cancer resistance and longer lifespans in larger species are theorized to result from compensatory evolutionary adaptations driven by reduced extrinsic mortality^2^. The bowhead whale exceeds 80,000 kg in mass and 200 years in lifespan. Both factors predispose it to accumulating large numbers of DNA mutations throughout life. To remain alive for so long it must possess uniquely potent genetic mechanisms to prevent cancer and other age-related diseases. However, primary research publications on genetic and molecular mechanisms of aging in the bowhead whale are scarce, consisting primarily of genome and transcriptome analysis^17–19^.

The multi-stage model of carcinogenesis posits that the transition from a normal cell to a cancer cell involves multiple distinct genetic “hits,” or mutations^20^. Larger and longer-living species might require greater numbers of “hits” for oncogenic transformation, given their greater cell number and increased lifespan. Indeed, there is experimental evidence to support this hypothesis. Rangarajan et al. found that while mouse fibroblasts require perturbation of 2 pathways for tumorigenic transformation (p53 and Ras), human fibroblasts require 5 hits (p53, pRb, PP2A, telomerase and Ras)^21^. A human should thus have a dramatically lower per-cell incidence of malignant transformation than a mouse, and as a result can maintain a larger number of cells for a longer period of time.

Species that are large-bodied and long-lived may be expected to have even more layers of protection against oncogenic transformation than humans. In support of this hypothesis, recent studies have identified copy number expansion and functional diversification of multiple tumor suppressor genes, such as *TP53* and *LIF*, in elephants and other taxa^3,5,22–24^. These studies identified multiple copies of *TP53* in the elephant genome, several of which were confirmed to be transcribed and translated in elephant fibroblasts and contributed to an enhanced apoptotic response to genotoxic stress^25^.

However, additional copies of p53 genes are unlikely to slow down aging^26,27^. One promising mechanism that could explain both cancer resistance and slower aging in long-lived mammals is more accurate or efficient DNA repair. Genetic mutations have been identified as causal factors in carcinogenesis for over a century^28^. Perhaps one of the most compelling lines of evidence supporting the role of DNA repair in the pathogenesis of aging and cancer comes from studies of mutants with accelerated aging phenotypes. Remarkably, most such mutants have defects in DNA repair enzymes^29–33^. Across species, several studies have also pointed toward improved DNA repair capacity and reduced mutation accumulation as characteristics associated with species longevity^34–38^. Here, we identify specific cellular and molecular traits characterizing bowhead whale cancer resistance and longevity that distinguish it from shorter-lived mammals including humans. We show that bowhead whale cells are not more prone to apoptosis and do not require additional genetic hits for malignant transformation relative to human cells. Instead, the bowhead whale relies on more accurate and efficient DNA double strand break (DSB) repair promoted by CIRBP and RPA2, as well as more efficient mismatch repair. This more “conservative” strategy that does not needlessly eliminate cells but repairs them may be critical for the long and cancer-free lifespan of the bowhead whale.

## Results

### Growth characteristics, cellular senescence, and cell death in the bowhead whale

Most human somatic cells lack telomerase activity and as a result undergo replicative senescence with serial passaging in culture^39^. Replicative and stress-induced senescence are important mechanisms for preventing cancer. Using TRF and TRAP assays to measure telomere length and telomerase activity, we found that bowhead whale skin fibroblasts, like human fibroblasts, lack telomerase activity and experience telomere shortening followed by replicative senescence with serial passaging in culture (Figure 1a, b). In both species, nearly all cells stained positive for senescence-associated β-galactosidase upon terminal growth arrest (Figure 1c, d). As in human fibroblasts, stable overexpression of human telomerase reverse transcriptase (*hTERT)* to maintain telomere length prevented replicative senescence in bowhead cells (Figure 1a). Senescence can also be induced by DNA damage. Like human cells, bowhead whale skin fibroblasts readily entered senescence but did not significantly induce cell death in response to 10 or 20 Gy of γ-irradiation (Figure 1c-e).

**Figure 1.**
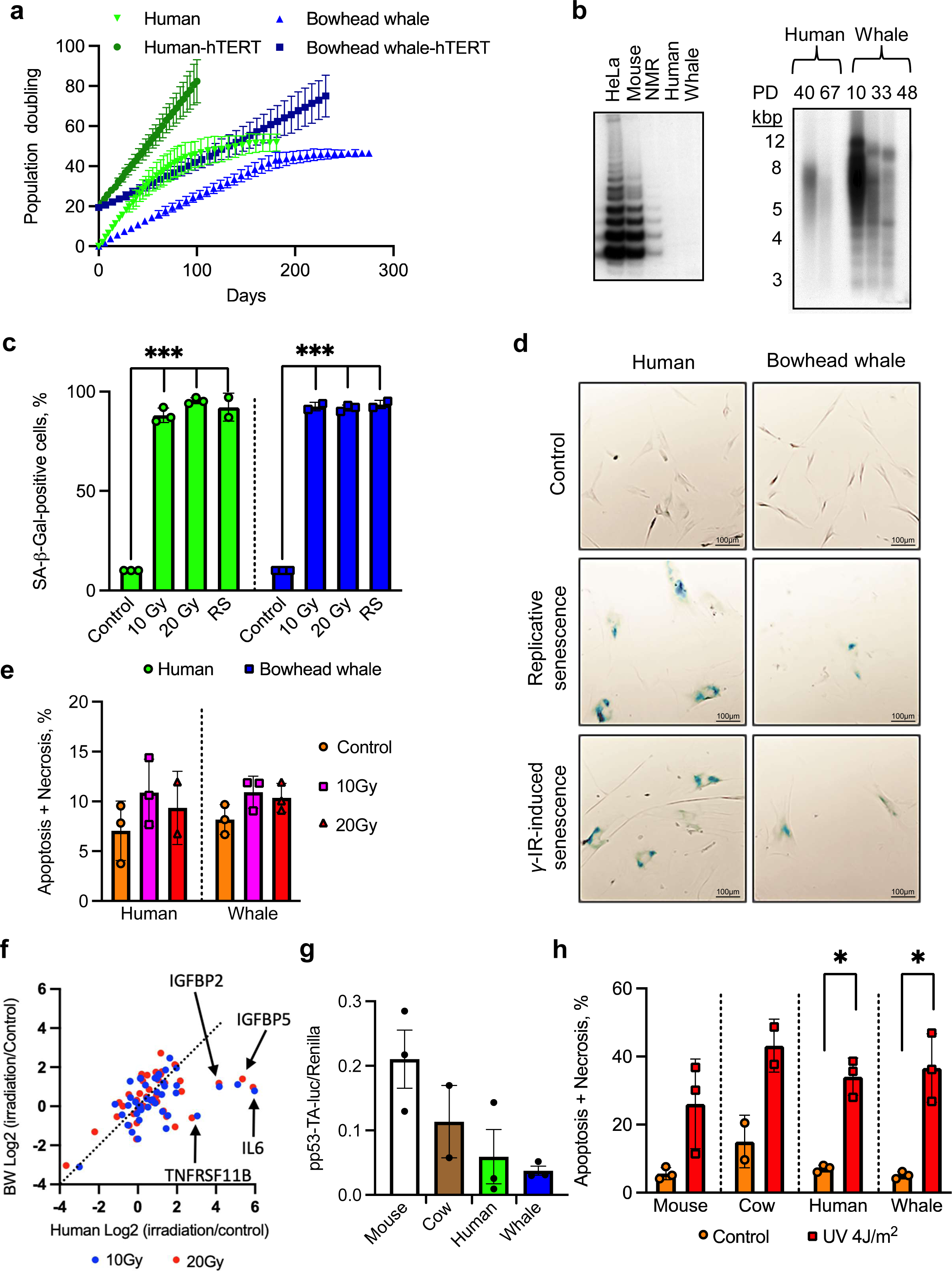
Bowhead whale fibroblasts exhibit senescence with reduced SASP and low basal p53 activity. **a**, Growth curves of primary and hTERT-immortalized skin fibroblasts (n=2 for each cell line). **b**, Telomerase activity and telomere length in skin fibroblasts. **c**, Quantification of β-gal–positive human and bowhead skin fibroblasts in response to *γ*-irradiation (12 days) and replicative senescence (n=3 for each species). **d**, Representative images of SA-β-gal staining of human and bowhead skin fibroblasts in response to *γ*-irradiation and replicative senescence. The bar is 100 µm. **e**, Apoptosis of human and bowhead whale fibroblasts in response to *γ*-irradiation. Three days after *γ*-irradiation, cells were harvested and subjected to an Annexin V apoptosis assay using flow cytometry (n=3 for each species). **f**, Log fold change of SASP mRNA expression in human and bowhead whale skin fibroblasts 12 days after *γ*-irradiation. **g**, Relative luciferase expression in mouse, cow, human and bowhead whale fibroblasts transfected with the p53 reporter vector. Data are shown as ratios of firefly/renilla luciferase (to normalize for transfection efficiency) expression 24 h after transfections (n=3 for mouse, human, BW; n=2 for cow). **h**, Apoptosis of mouse, cow, human and bowhead whale fibroblasts in response to UVC. Two days after UVC, cells were harvested and subjected to an Annexin V apoptosis assay using flow cytometry. Error bars represent mean ± SD. * p<0.05, *** p<0.001. Welch’s t-test was used to quantify the significance. Whale, bowhead whale; NMR, naked mole-rat. RS, replicative senescence.

Interestingly, transcriptome analysis of human and bowhead whale senescent fibroblasts showed reduced induction of senescence-associated secretory phenotype (SASP) factors in bowhead whale fibroblasts (Figure 1f) relative to human cells.

Paracrine effects of SASP on surrounding cells are thought to contribute to age-related diseases and carcinogenesis. These transcriptomic differences may indicate that senescence is able to preserve its anti-cancer function in the bowhead with reduced harmful paracrine signaling.

To test whether increased p53 activity could contribute to cancer resistance in the bowhead whale, we transiently transfected primary mouse, cow, human and bowhead whale skin fibroblasts with a luciferase reporter vector containing a p53-response element. The bowhead whale cells had the lowest basal p53 activity of the species tested (Figure 1g). Additionally, we did not observe any differences in the induction of apoptosis in response to UVC between species (Figure 1h). Together, our results argue against the idea that increased clearance of damaged cells through apoptosis contributes to cancer resistance in the bowhead whale.

### Requirements for oncogenic transformation of bowhead whale cells

We initially identified a minimal combination of oncogene and tumor suppressor hits required for *in vitro* malignant transformation of bowhead whale skin fibroblasts using the soft agar assay, which measures anchorage-independent growth, a hallmark of cancer. While normal cells undergo growth arrest or programmed cell death (anoikis) in soft agar, malignant cells continue to grow without substrate adhesion and form visible colonies^40^. We introduced constructs targeting oncogene and tumor suppressor pathways into primary skin fibroblasts with PiggyBac (PB) transposon vectors, which integrate into the genome and drive stable expression. Since bowhead whale primary fibroblasts, like human fibroblasts, exhibit progressive telomere shortening and lack telomerase activity (Figure 1b), we used cell lines expressing (*hTERT*) to bypass replicative senescence.

In agreement with published findings, malignant transformation of human *hTERT*+ fibroblasts required combined expression of H-Ras^G12V^, SV40 Large T (LT) antigen (which binds and inactivates p53 and the Rb family of tumor suppressors), and SV40 Small T (ST) antigen (which binds and inactivates PP2A) (Figure 2a)^21^. Rather than requiring hits to additional pathways, however, bowhead whale *hTERT*+ fibroblasts were transformed by H-Ras^G12V^ and SV40 LT alone, suggesting that bowhead cells may require fewer genetic mutations to become cancerous compared to human cells (Figure 2a). These findings were supported by mouse xenograft assays, in which the number of hits needed for tumor growth matched findings from soft agar (Figure 2b).

**Figure 2.**
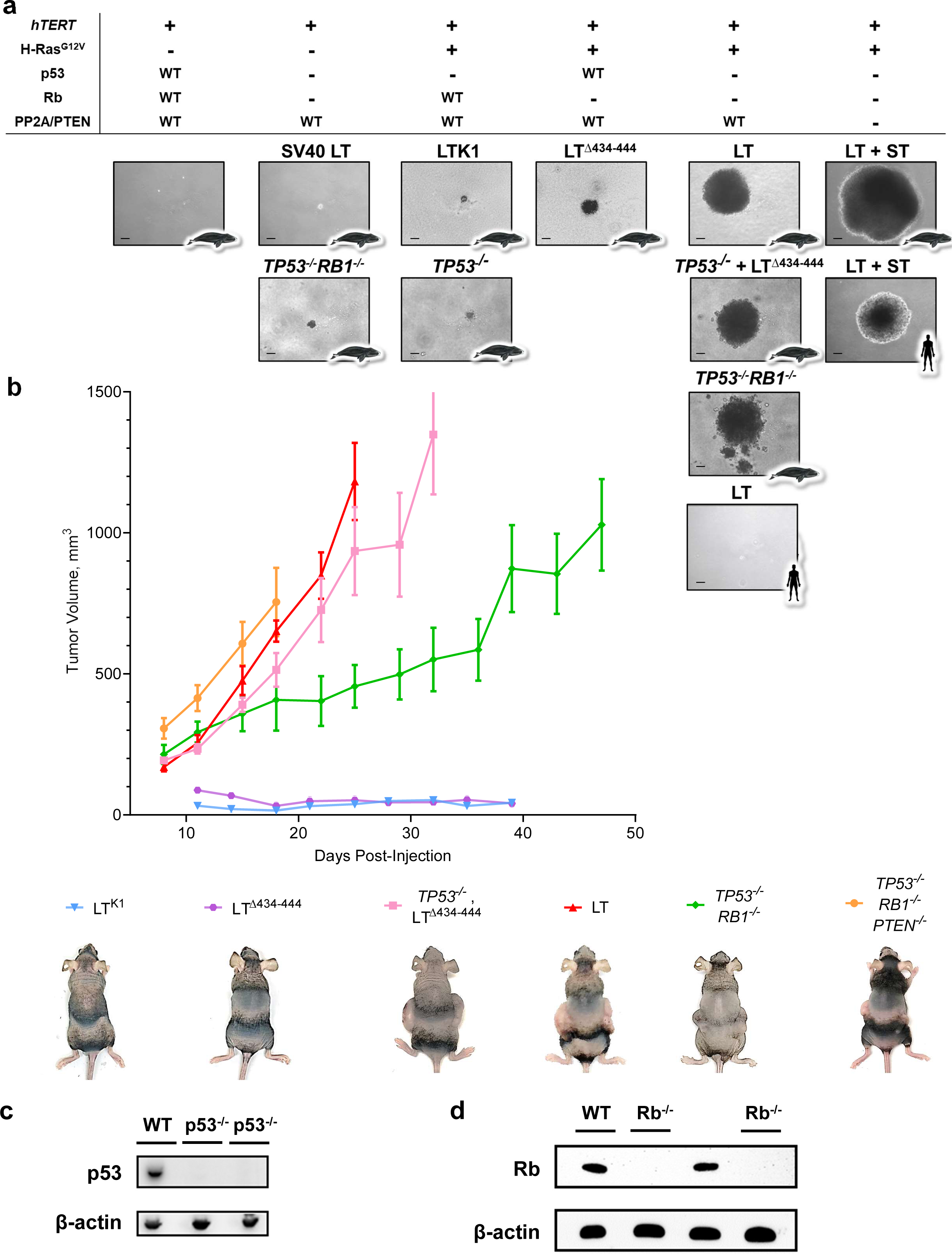
Fewer tumor suppressor hits are required for oncogenic transformation of bowhead fibroblasts than for human fibroblasts. **a,** Images of representative fibroblast colonies for tested cell lines after 4 weeks of growth in soft agar. The top panel indicates whether the cell lines in the column below have the indicated protein overexpressed (+), inactivated (-), or expressed in the active endogenous form (WT). Text above individual images indicate for that cell line whether tumor suppressors are inactivated through genetic knockout or SV40 Large T (or LT mutants) or Small T (ST) antigen. Icons in corners of images indicate species. Scale bar represents 250 µm. **b,** Volumetric growth curves for the indicated bowhead whale fibroblast cell lines in mouse xenograft assays. All cell lines shown stably express H-Ras^G12V^ and *hTERT* in addition to the genotype indicated in the figure legend. Data points represent averages from 3 immunodeficient nude mice injected bilaterally (6 injections) for each cell line, except for *TP53^-/-^RB1^-/-^* double knockouts, for which 2 independent cell lines were tested, for a total of 6 mice/12 injections. Experiments were terminated based on predetermined thresholds for maximum tumor length or duration of experiment as described in Methods. Images in the legend show a representative mouse for the indicated cell line at the final measured time point. Error bars show SEM. **c,** Western blot for p53 protein in clonally isolated fibroblast colonies following CRISPR targeting of *TP53*. Underlined lanes indicate colonies selected for further validation and experiments. **d,** Western blot for Rb protein in clonally isolated fibroblast colonies following CRISPR targeting of *RB1* on an existing p53 knockout background.

We next sought to confirm these findings at the genetic level, through CRISPR editing of individual tumor suppressor genes in bowhead fibroblasts. While the sequenced bowhead genome has not revealed copy number expansion of canonical tumor suppressor genes^17,18^, CRISPR knockout allows for more precise quantification of the number of genetic mutations required for oncogenesis. While the most important target of SV40 LT is thought to be Rb (*RB1* gene), it is also known to inactivate p130 and p107, two additional members of the Rb-family, providing some level of functional redundancy. Using CRISPR, we introduced targeted mutations into the bowhead *RB1* gene, along with *TP53* (the other target of LT), and *PTEN* (an upstream inhibitor of Akt signaling commonly mutated in human cancers and operating in the same pathway as PP2A). Following transfection of *hTERT*+ bowhead fibroblasts with Cas9-guide RNA ribonucleoprotein complexes targeting each of the aforementioned genes, we screened clonally isolated colonies for loss of the targeted protein by Western blot (Figure 2c, d, Extended Data Figure 1a, b). We additionally screened the colonies with luciferase reporter assays to confirm loss of protein function and activity (Extended Data Figure 1c, d). For each selected clone, we sequenced the CRISPR-targeted genes to confirm homozygous knockout at the genetic level and determine the causal mutations (Supplementary Figures 1, 2). Through this strategy, we generated single and compound homozygous knockout bowhead whale fibroblasts for *TP53*, *RB1*, and *PTEN*. In agreement with our initial findings, genetic inactivation of *TP53* and *RB1* in bowhead whale fibroblasts expressing *hTERT* and H-Ras^G12V^ was sufficient for malignant transformation in both soft agar and mouse xenograft assays (Figure 2).

These findings suggest that despite its larger size and longer lifespan, the cells of the bowhead whale unexpectedly require fewer mutational hits for malignant transformation than human cells.

### Mismatch repair, excision repair and mutagenesis in the bowhead whale

As defects in mismatch repair genes are well-characterized drivers of oncogenesis, we assessed the efficiency of mismatch repair (MMR) in bowhead whale cells using a reporter assay that measures cellular correction of a targeted G/T mismatch introduced to a plasmid *in vitro* ^41^. We found that correction of the mismatch was significantly more efficient in whale cells than in mouse, cow, and human cells (Extended Data Figure 2a).

We next assessed the efficiency of nucleotide excision repair (NER) and base excision repair (BER) repair in bowhead whale cells. NER is primarily responsible for removing helix-distorting DNA lesions. To quantify NER activity, we utilized a host cell plasmid reactivation assay^33^ and quantified clearance of cyclobutane pyrimidine dimers (CPDs) by ELISA to measure repair of UVC-induced DNA damage. NER efficiency by plasmid reactivation was similar between bowhead and human cells (Extended Data Figure 2b), but the kinetics of CPD removal tended to be slower in whale cells (Extended Data Figure 2c). BER is responsible for ameliorating many types of spontaneous DNA base damage, such as oxidation and deamination. The efficiency of BER, as measured by the plasmid reactivation assay, trended toward higher BER activity in bowhead whales compared to human cells, but this difference was not statistically significant (Extended Data Figure 2d).

We found that PARP activity in bowhead fibroblasts exposed to H_2_O_2,_ and γ-irradiation was dramatically higher than in human cells (Extended Data Figure 3a, b). Basal PARP activity was also much higher in untreated bowhead whale nuclear extracts (Extended Data Figure 3c). PARP proteins are recruited to sites of DNA damage, where they participate in the DNA damage response and repair. Bowhead whale cells also displayed higher survival rates after H_2_O_2_ treatment in comparison to human cells (Extended Data Figure 3d). When we measured the kinetics of damage repair after H_2_O_2_ treatment by alkaline comet assay, repair was slightly accelerated in bowhead whale relative to human cells, which may relate to its increased PARP activity (Extended Data Figure 3e).

To determine whether whale cells might accumulate fewer mutations after DNA damage, we measured mutation frequency following treatment with the potent mutagen and alkylating agent N-ethyl-N-nitrosourea (ENU) using single-molecule, quantitative detection of low-abundance somatic mutations by high-throughput sequencing (SMM-seq)^42^. ENU treatment resulted in a statistically significant increase in somatic mutation frequency in fibroblasts from all tested species (Extended data Figure 2e). Specifically, we found that mouse cells showed the greatest increase in ENU-induced single nucleotide variants, while bowhead whale cells experienced the lowest mutation induction. The levels of induced mutational load in cow and human cells were intermediate, in line with their relative lifespans. This suggests a correlation between the rate of mutation induction and maximum lifespan among the included species (Extended Data Figure 2e), confirming previous findings by us and others^4344^.

Importantly, the excessive mutational burden observed in ENU-treated cells predominantly comprised an increased fraction of A to T transversions (Extended Data Figure 2f), the type of mutation preferentially induced by ENU^42^.

We additionally compared mutation induction in response to chemical mutagen treatment in the bowhead whale and human using the HPRT mutagenesis assay, which relies on loss of HPRT activity after mutagen treatment^45^. The *HPRT* gene exists as a single copy on the X chromosome in male mammalian cells, a feature we found to be true for the bowhead (see Methods). We treated primary fibroblast lines from male bowhead whale and human with ENU and then plated cells in selective media containing 6-thioguanine, which kills cells with functional HPRT. Despite a slightly higher sensitivity to ENU in bowhead whale cells as indicated by colony-forming efficiency in non-selective media, the rate of HPRT mutant colony formation was markedly lower in bowhead whale than human fibroblasts, an effect which remained significant after adjusting for plating efficiency (Extended Data Figure 2g, h). This result supports the above SMM-seq data that bowhead whale cells may possess more accurate DNA repair than humans. There was no difference in the rate of apoptosis in response to ENU treatment in human and bowhead whale fibroblasts, suggesting that apoptosis is not responsible for the observed differences in ENU sensitivity (Extended Data Figure 2i). To further validate these findings, we also measured HPRT mutant colony formation after γ-irradiation. As with ENU, we observed markedly lower HPRT mutant colony formation in bowhead whale cells (Extended Data Figure 2j, k).

### Double-strand break repair and chromosomal stability in the bowhead whale

DNA DSBs are toxic if not repaired and may lead to mutations through inaccurate repair. DSBs are repaired through two major pathways: non-homologous end joining (NHEJ) and homologous recombination (HR). To assess relative NHEJ and HR efficiencies, we integrated fluorescent GFP-based reporter cassettes ^46^ (Extended Data Figure 5a) into fibroblasts from mouse, cow, human and bowhead whale. Following DSB induction with I-SceI, we observed markedly elevated NHEJ efficiency in bowhead whales relative to other species (Figure 3a, Extended Data Figure 4a). We also found that HR efficiency is significantly higher in whale cells than in human cells (Figure 3b).

**Figure 3.**
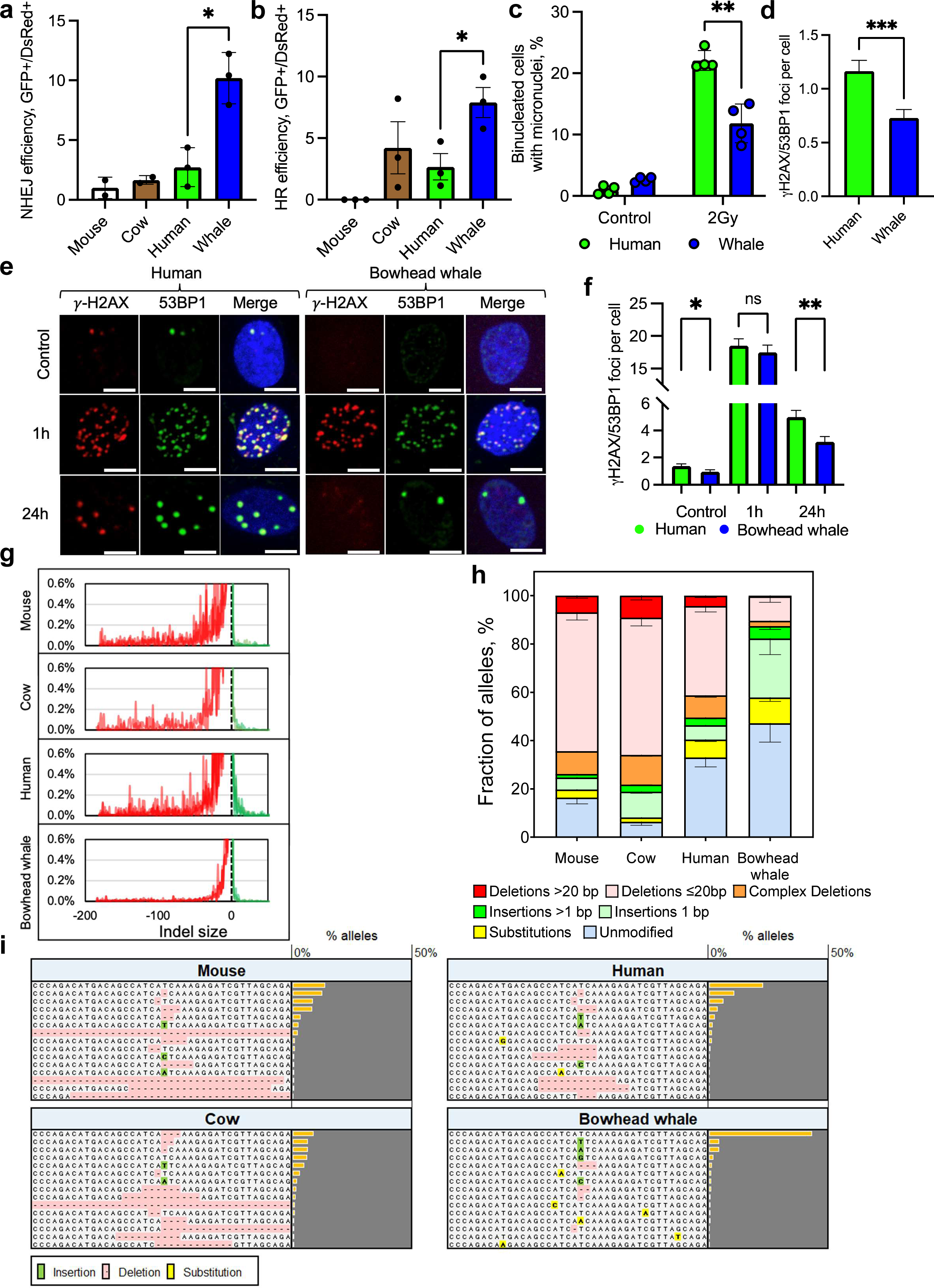
The bowhead whale repairs DSBs with higher accuracy and efficiency than other mammals. **a,b**, NHEJ and HR efficiency were measured using fluorescent reporter constructs. Successful NHEJ and HR event leads to reactivation of the GFP gene. NHEJ and HR reporter constructs were integrated into primary, low passage skin fibroblasts. NHEJ and HR repair efficiency were assayed by transfecting cells with I-SceI expression vector and a DsRed plasmid as a transfection control. The repair efficiency was calculated as the ratio of GFP+/DsRed+ cells. Experiments were repeated at least 3 times for each cell line. Error bars represent SD. * p<0.05 (Welch’s t-test). Whale, bowhead whale. **c**, Percent of binucleated cells containing micronuclei in human and bowhead whale fibroblasts after 2Gy *γ*-irradiation (n=4). Error bars represent SD. ** p<0.005 (Welch’s t-test). **d,** Endogenous γ-H2AX/53BP1 foci in human and whale cells. Results are combined from two independent experiments. 200 nuclei were analyzed. Error bars represent SEM. *** p<0.001 (two-tailed t-test). **e,** Representative confocal images of human and bowhead whale cells stained for γ-H2AX and 53BP1 at no treatment (control) and 1h-24h after bleomycin treatment at concertation 5μg/mL. Scale bar indicates 10 µm. **f,** Quantification of γH2AX/53BP1 foci with and without DSB induction. Exponentially growing cells were treated for 1h with bleomycin (BLM) at concertation 5μg/mL, washed twice with PBS and fresh media was added. At indicated time-points cells were fixed and processed for immunocytochemistry. Foci were counted by eye in green and red channels. 150-170 nuclei were analyzed. Error bars represent SEM. *p<0.05, ** p<0.01 (two-tailed t-test). **g**, Histograms of CRISPR indel size distribution by species. Data for biological replicates are superimposed and partially transparent with lines connecting data points for each sample. Unmodified alleles and alleles with substitutions only are excluded from this analysis. **h**, Distribution of sequenced PTEN allele variants by species after CRISPR DSB induction at a conserved region of the endogenous PTEN gene. Data are averages from multiple primary dermal fibroblast lines isolated from different individual animals for bowhead whale (n=3), human (n=3), cow (n=2), and mouse (n=3). Error bars represent SEM. **i**, Allele plots showing 15 most frequent allele types after CRISPR for one representative cell line per species. Sequences are displayed within a window centered on the cleavage site and extending 20 bp in each direction. Data bars and values indicate proportion of total alleles represented by each row. For the purposes of this display and quantification, all individual alleles with identical sequences in the 40-bp window have been pooled, so rows represent composites of alleles that may differ outside the display window.

To examine whether this more efficient DSB repair could promote chromosomal stability, we measured formation of micronuclei induced by γ-irradiation and by I-SceI cleavage. One potential outcome resulting from an unrepaired DSB in mitotic cells is the loss of an acentric chromosome fragment, which can be measured as the formation of a micronucleus. We found that bowhead whale fibroblasts accumulated fewer micronuclei than human fibroblasts after 2 Gy γ-irradiation (Figure 3c, Extended Data Figure 4b).

We also observed that DSB induction with I-SceI increased the rate of micronucleus formation, likely reflecting acentric fragment loss, and that this rate was reduced in the bowhead whale compared to human (Extended Data Figure 4c). Thus, the more efficient rejoining of DSB ends observed in bowhead whale cells appears to guard against chromosomal instability.

We also measured resolution of γH2AX and 53BP1 foci, which mark cellular DSBs. We found that endogenous levels of these foci are significantly lower in whale cells, suggesting reduced baseline burden of DSBs (Figure 3d). We observed that the kinetics of DSB repair after γ-irradiation are not faster in the bowhead whale than in human cells (Extended Data Figure 4d, e)^33^. We further tested the ability of bowhead whale cells to resolve DSBs after treatment with the DSB-inducing drug bleomycin. We observed similar induction of foci one hour after bleomycin treatment in human and whale cells; however, fewer foci remained in cells from the bowhead whales than in those from humans after 24 hours (Figure 3e, f), indicating a reduced burden of residual unrepaired damage in the bowhead whale.

### Fidelity of DSB repair in the bowhead whale

As mutations resulting from inaccurate DSB repair can promote cancer development, we next sought to assess the fidelity of DSB repair in the bowhead whale. Sequencing and analysis of repair junctions from integrated NHEJ reporter (Extended Data Figure 5a, b) and extra-chromosomal NHEJ reporter (Extended Data Figure 5a, c) assays suggested higher fidelity of NHEJ in bowhead whale cells: compared to human, the bowhead whale is less prone to producing deletions during the repair of incompatible DNA termini and far more frequently joins ends without deleting any bases beyond the small overhang region.

We also measured the fidelity of repair at an endogenous genomic locus. To systematically compare mutational outcomes of CRISPR break repair in the bowhead whale to those of humans and shorter-living mammals, we performed CRISPR transfections in primary fibroblast lines, 2-3 individual animals per species, from bowhead whale, human, cow, and mouse, and used deep amplicon sequencing of the targeted locus to generate detailed profiles of repair outcomes. We took advantage of the fact that exon 1 of the *PTEN* tumor suppressor gene is highly conserved across mammals, with 100% sequence identity across included species (Supplementary Figure 3). We were therefore able to examine species-specific DSB repair outcomes at an endogenous genomic locus while minimizing intra-species variation in the break-proximal sequence context.

Analysis of sequencing data revealed species-specific repair outcomes, which were consistent across cell lines derived from multiple individual animals of each species (Figure 3g-i). In human, cow, and mouse, the most common mutational outcomes were deletions. In contrast, the bowhead was the only species for which a single-base insertion was the most common mutational event. The frequency of unmodified alleles, which are known to occur after error-free repair of CRISPR DSBs^47,48^, was the highest in bowhead whale (Figure 3h). Sequencing of untreated control samples confirmed that the detected insertions and deletions were CRISPR-induced (Supplementary Table 1). As analysis of CRISPR RNP transfection efficiency by flow cytometry and cleavage efficiency by digital droplet PCR showed similar CRISPR efficiencies across species (Extended Data Figure 5d, Supplementary Figure 4a, b), differences observed in the unmodified allele fraction most likely result from differences in repair fidelity. While small indels predominated in all species, we observed a marked inverse correlation between the frequency of large deletions and species lifespan, with the bowhead producing fewer large deletions than human, cow, and mouse (Figure 3g-i). Intriguingly, this reduction in large deletions was not accompanied by reduced microhomology usage (Extended Data Figure 5f, g). When we assigned frequency-based percentile ranks from most negative to most positive indel size (largest deletions to largest insertions), we observed a strong correlation between species lifespan and 5th percentile indel size, corresponding to large deletions (Pearson’s *r*=0.85, p=0.0009) (Extended Data Figure 5e, Supplementary Table 2). The results of these experiments suggest a greater fidelity of DSB repair in the bowhead whale relative to humans and other mammals.

To determine whether these differences in repair outcomes of targeted DSBs might predict the types of genomic changes accumulated through spontaneous cellular DNA damage, we performed whole genome sequencing (WGS)^49^ of bowhead whale, human and mouse fibroblast-derived tumor xenografts and assessed somatic mutations through comparison to parental non-transformed primary fibroblast cultures sequenced in tandem (Extended Data Figure 1e). The frequency of spontaneous de novo somatic single nucleotide variants (SNVs) was significantly lower in bowhead whale tumor xenografts than in human and mouse (Extended Data Figure 1g). Intriguingly, we observed no differences in the relative proportions of each type of SNV across species, suggesting shared underlying mutational drivers during tumor evolution despite differences in overall mutation rate (Extended Data Figure 1f). We further assessed WGS data for small indels (Extended Data Figure 1h, i) and large structural variants (SVs) (Extended Data Figure 1j-l) across species; strikingly, whale tumors were characterized by a significant reduction in both small and large deletions, as well as small insertions, large duplications and inversions (Extended Data Figures 1g-l, Supplementary Table 3). SV size distributions were remarkably different in bowhead whale tumors in comparison to human and mouse tumors: whale tumors showed a significant reduction in the proportion of large SVs (>500Kb, p < 0.0001, Extended Data Figure 1m, n). Altogether, these data demonstrate consistent reductions in both the frequency and size of inserted and deleted bases in bowhead whale cells relative to those of shorter-lived mammals in response to both nuclease-induced and endogenous DNA breaks. These differences are likely to reduce the accumulation of deleterious genomic instability over time.

### CIRBP contributes to high DSB repair efficiency and chromosomal stability in the bowhead whale

To identify mechanisms contributing to the efficiency and accuracy of DSB repair in the bowhead whale, we compared expression of DNA repair proteins in the bowhead whale to other mammalian species by Western blot, quantitative mass spectrometry, and transcriptome sequencing (Figure 4a, Extended Data Figures 6-7). Unexpectedly, we found that levels of three canonical NHEJ proteins-Ku70, Ku80, and DNA-PKcs- are substantially higher in human cells than any other species tested (Figure 4a, Extended Data Figure 6f), while their abundance in the bowhead whale appears to be at more typical mammalian levels. We speculate that the unusually high levels of Ku/DNA-PKcs in humans may be a human-specific adaptation to promote DSB repair and genome stability.

**Figure 4.**
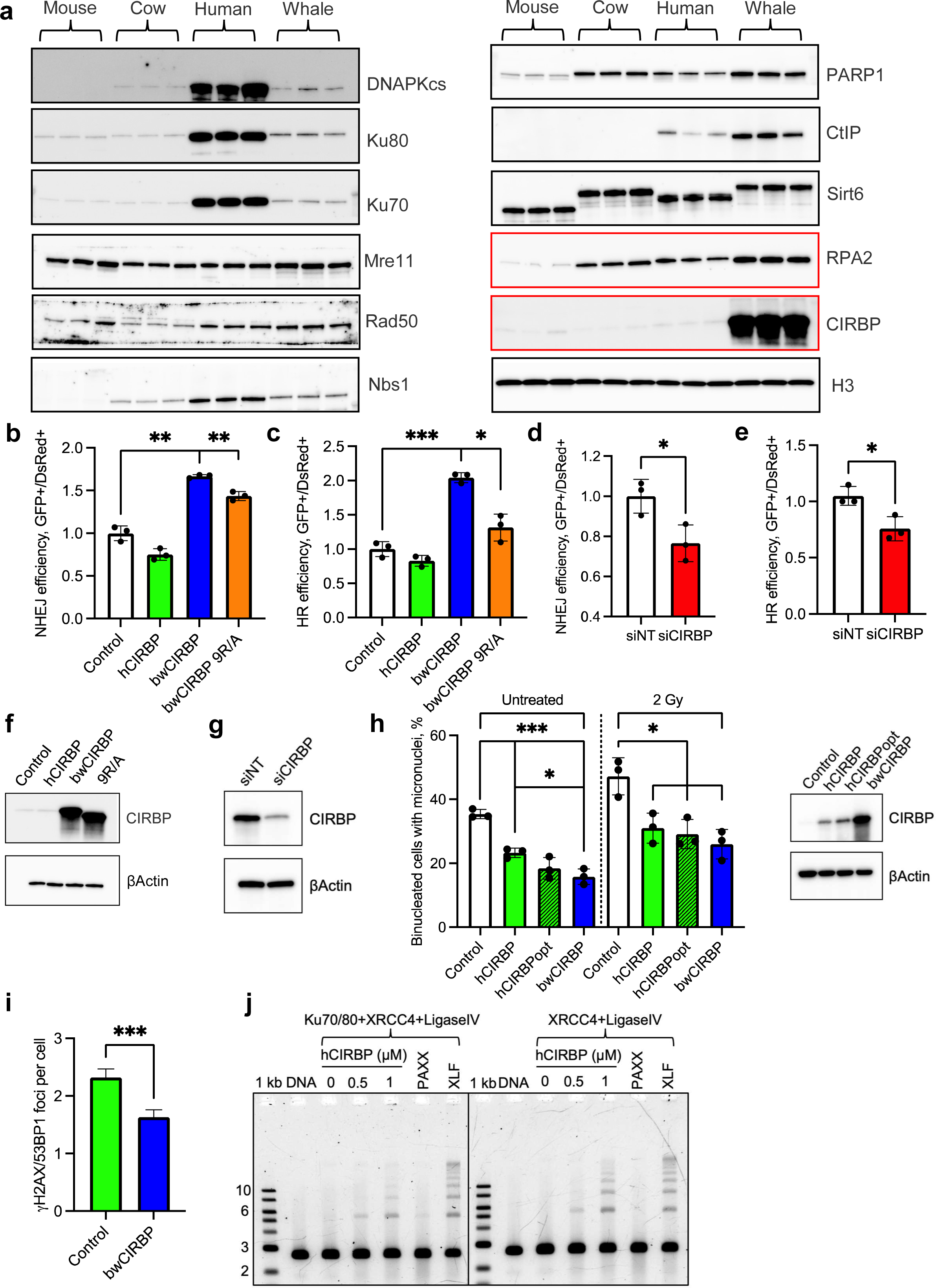
CIRBP is highly expressed in bowhead whale fibroblasts and promotes DNA DSB repair and genome stability. **a**, Western blots of DNA repair proteins in primary fibroblasts from different species. **b-c**, bwCIRBP promotes NHEJ and HR in human cells as measured by flow cytometric GFP-reporter assays (see Methods). In these assays DSBs are induced within inactive NHEJ or HR reporter cassettes by expressing I-SceI endonuclease. Successful NHEJ or HR events lead to reactivation of the fluorescent GFP reporters that are scored by flow cytometry. All experiments in these figures were repeated at least 3 times. **d,e**, Knockdown of CIRBP in bowhead whale fibroblasts decreases NHEJ and HR efficiency. siNT = non-targeting siRNA. **f**, Western blot of human fibroblasts overexpressing human CIRBP, whale CIRBP or 9R/A mutated whale CIRBP; **g,** Western blot of bowhead whale fibroblasts with knockdown of CIRBP. **h**, Overexpression of CIRBP decreases the percentage of binucleated cells containing micronuclei in human cells 3d after 2Gy *γ*-irradiation (n=3) (left panel); Western blot of human fibroblasts overexpressing human CIRBP, human CIRBP with optimized codons and whale CIRBP (right panel). Error bars represent mean ± SD. * p<0.05, ** p<0.01, *** p<0.001 (Welch’s t-test). siNT-negative control siRNAs that do not target any gene product. **i,** Number of endogenous γH2AX/53BP1 foci in human cells with bwCIRBP overexpression. Error bars represent SEM. *** p<0.001 (two-tailed t-test). **j,** CIRBP stimulates NHEJ-mediated ligation *in vitro in*. Linearized pUC19 plasmid with cohesive ends was mixed with human recombinant proteins XRCC4/ Ligase IV complex, and 0 to 1 μM CIRBP. Where indicated, reaction mixtures contained Ku70/80 heterodimer, PAXX dimer or XLF dimer. The reaction mixtures were incubated for 1 hr at 30°C, proteins were denatured with SDS at 65°C and loaded onto agarose gel. Each sample were loaded onto 0.7% agarose gel, followed by gel electrophoresis. The gel was stained with ethidium bromide.

However, we consistently observed a strikingly higher abundance of cold-inducible RNA-binding protein (CIRBP) in cells and tissues of the bowhead whale than in other mammalian species (Figure 4a, Extended Data Figure 6a, b, d, f). Levels of PARP1, a functional partner of CIRBP during DNA repair^50^, were also increased relative to human and showed even greater enrichment at an *in vitro* DSB substrate (Extended Data Figure 6g). Interestingly, we also found that the CtIP protein, which is required for efficient HR^51^, is more abundant in whale cells compared to humans. It is possible that CtIP upregulation in the whale contributes to better HR compared to humans.

CIRBP is an RNA- and PAR-binding protein whose expression is induced by a variety of cellular stressors including cold shock, hypoxia, and UV irradiation^50,52–54^. CIRBP has been shown to bind the 3’ UTR of mRNAs that encode proteins involved in cellular stress and DNA damage responses and promote their stability and translation^55–58^.

There is also evidence for a more direct role of CIRBP in DNA repair: PARP-1-dependent localization of CIRBP to sites of DNA damage promotes DSB repair and antagonizes micronucleus formation^50^.

To test whether CIRBP contributes to efficient NHEJ and HR in bowhead whale cells, we overexpressed human (hCIRBP) and bowhead whale (bwCIRBP) in human reporter cells. Overexpression of bwCIRBP, but not hCIRBP, enhanced NHEJ and HR efficiencies in human cells (Figure 4b, c, f). Conversely, CIRBP depletion in bowhead whale cells by siRNA significantly reduced NHEJ and HR efficiency (Figure 4d, e, g).

Consistent with published observations, overexpression of bwCIRBP with nine arginines in the repeated RGG motif mutated to alanine (9R/A), which impairs CIRBP’s ability to bind to PAR-polymers^50^, failed to stimulate HR and reduced stimulation of NHEJ (Figure 4b, c, f).

To test the effects of CIRBP overexpression on chromosomal stability, we quantified the formation of micronuclei in human cells after γ-irradiation. In cells overexpressing wild-type hCIRBP, a codon-optimized hCIRBP, and bwCIRBP, we observed a significant decrease in both basal and γ-irradiation-induced micronuclei in all CIRBP-overexpressing cells relative to control vector (Figure 4h). Micronucleus formation appeared to decrease as CIRBP protein expression increased. To probe the relationship between high NHEJ efficiency, high CIRBP expression, and genome stability, we also overexpressed human and bowhead whale CIRBP in human NHEJ reporter cells and observed a reduction in I-SceI-induced micronuclei in cells overexpressing CIRBP (Extended Data Figure 8f).

We next examined the effect of CIRBP overexpression on formation of gross chromosomal aberrations in human cells exposed to γ-irradiation. We observed a markedly decreased frequency of chromosomal aberrations after irradiation in cells overexpressing hCIRBP and bwCIRBP (Extended Data Figure 8g). This further suggests a protective effect of high CIRBP expression on genomic stability. Basal γH2AX/53BP1 foci were reduced by bwCIRBP overexpression, consistent with improved genome stability (Figure 4i).

The human and bowhead CIRBP proteins differ by only 5 C-terminal amino acids, which do not overlap with any residues of known functional significance (Extended Data Figure 9a). Substitution of these 5 codons in hCIRBP with bowhead codons increased protein expression, while substitution of bwCIRBP with the 5 hCIRBP codons decreased it (Extended Data Figure 9d). Interestingly, although CIRBP abundance increased following introduction of the 5 bowhead substitutions, it did not achieve the expression levels of bwCIRBP, suggesting that synonymous changes to the mRNA coding sequence contribute to higher translation efficiency of bwCIRBP. Consistent with this notion, bwCIRBP has a higher codon adaptation index (CAI)^59^ than hCIRBP (Extended Data Figure 9e). We also conducted a phylogenetic analysis of the CIRBP variant present in the bowhead whale. Serine 126 appears to be ancient, already present in bovids and bats. The unique cluster of 4 amino acids starting at position 147 is more recent, only appearing in *Balaenopteridae*, the baleen whales (Extended Data Figure 9b). All baleen whales are very large and long-lived. Interestingly, analysis of primary fibroblasts from other marine mammals and hippopotamus (who share a common ancestor with whales), showed that CIRBP is similarly abundant in the humpback whale but not in sea lions or hippos, while dolphins showed a very mild increase in CIRBP compared to other mammals (Extended Data Figure 7e).

We did not observe significant upregulation of canonical CIRBP targets in the bowhead whale following DNA damage. However, CIRBP knockdown showed a trend towards reducing RPA2 levels (Extended Data Figure 8b). Conversely, bwCIRBP overexpression in human cells showed a trend towards upregulation of RPA2 levels (Extended Data Figure 8c).

When we compared the PAR-binding affinity of bwCIRBP to that of hCIRBP through fluorescence polarization (FP) measurements of labeled PAR, we observed similar K_D_ values for both proteins, indicating similar affinity for PAR. K_D_ values were lower for longer PAR polymers (PAR_28_ or PAR_16_) than shorter PAR chains (PAR_8_), indicating higher-affinity binding of CIRBP to the longer polymers (Extended Data Figure 8j). As CIRBP is present in over 10-fold excess in the whale, there is likely to be greater overall PAR-binding capacity in whale cells. Intriguingly, despite similar affinity for PAR, bwCIRBP produced a greater increase in FP of labeled PAR than did hCIRBP, indicating a stronger effect on PAR hydrodynamics. This raises the possibility that amino acid differences between the two proteins might lead to differences in binding conformation or stoichiometry of the CIRBP-PAR complex.

We next investigated the direct involvement of CIRBP in the DSB repair process. We observed that in bowhead whale cells, the majority of CIRBP is in the nuclear soluble fraction, but some is always associated with chromatin. This association with chromatin appeared to be in large part RNA-dependent (Extended Data Figure 8a).

Chen et al ^50^ demonstrated CIRBP recruitment to laser induced DSBs. To confirm this result using a different method, we treated whale cells with the DSB-inducing agent neocarzinostatin (NCS) and tested CIRBP enrichment in the chromatin fraction. Within minutes after the addition of NCS, CIRBP became transiently enriched in the chromatin fraction (Extended Data Figure 8d). Damage-induced CIRBP enrichment was sensitive to RNase A treatment, suggesting that local RNA binding contributes to the association of CIRBP with chromatin upon DNA damage (Extended Data Figure 8e).

Upon *in vitro* incubation with various nucleic acid substrates, recombinant human CIRBP produced a concentration-dependent electrophoretic mobility decrease for both RNA and DNA (Extended Data Figure 8k, l), lending support to prior findings of DNA binding by CIRBP^60,61^. With sufficient CIRBP, nearly all sheared RNA and DNA fragments were retained in the well, suggesting an ability of CIRBP to physically tether or aggregate nucleic acid fragments.

We next investigated whether CIRBP directly facilitates *in vitro* end joining of linearized plasmid incubated *in vitro* with human XRCC4-Ligase IV complex. We observed that the ligation of cohesive DNA ends was markedly enhanced by CIRBP in a concentration-dependent manner (Figure 4j). In contrast, the addition of the accessory protein PAXX^62^ failed to stimulate ligation. Interestingly, without addition of Ku70/Ku80, ligation stimulation by CIRBP was more pronounced and almost comparable to stimulation by XLF, which directly interacts with the XRCC4-Ligase IV complex and is considered a core component of the NHEJ machinery^63^. This result suggests that CIRBP directly stimulates NHEJ and can promote NHEJ in the absence of Ku70/Ku80. In the whale, abundant CIRBP may compensate for lower levels of Ku70/Ku80 relative to human.

We additionally tested the effect of bwCIRBP overexpression on repair fidelity in human cells and observed a reduction in indel rates (Figure 5c, Extended Data Figure 8h). We also knocked down CIRBP in whale cells harboring an integrated NHEJ reporter and assessed the mutation spectrum after I-SceI-induced DSBs by sequencing. We observed an increase in deletions, upon CIRBP knockdown (Extended Data Figure 8i), suggesting that CIRBP also contributes to repair fidelity.

**Figure 5.**
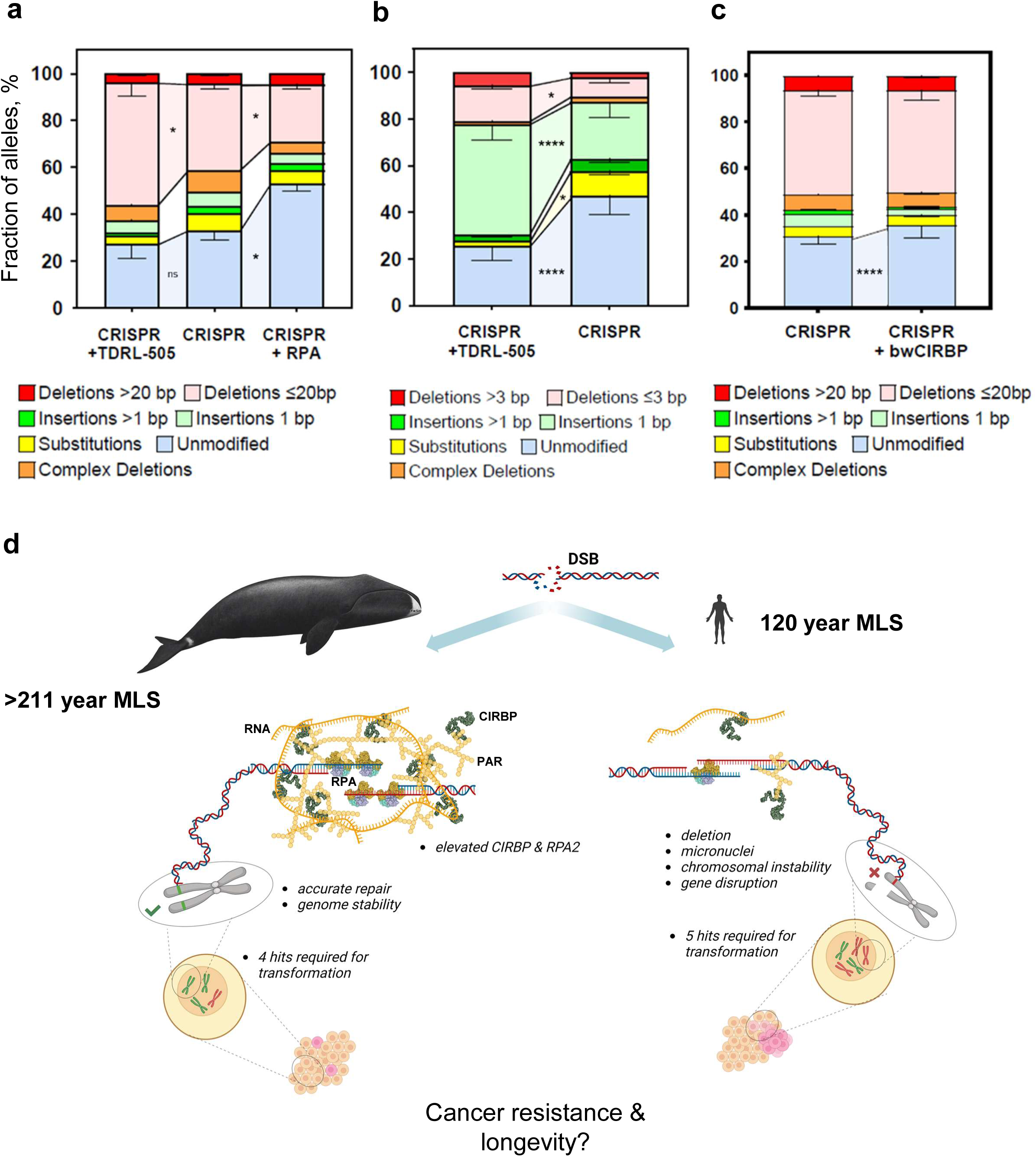
RPA and CIRBP contribute to increased DNA repair fidelity. **a,** Distribution of sequenced PTEN allele variants in human primary fibroblasts treated with TDRL-505 or rhRPA after CRISPR DSB induction at a conserved region of the endogenous PTEN gene. Data are averages from experiments performed in triplicate. Error bars represent SEM. **b,** Distribution of sequenced PTEN allele variants by species in bowhead whale primary fibroblasts treated with TDRL-505 after CRISPR DSB induction at a conserved region of the endogenous PTEN gene. Data are averages from experiments performed in triplicate. Error bars represent SEM. **c,** Distribution of sequenced PTEN allele variants by species in human fibroblasts with lentiviral overexpression of luciferase or bwCIRBP after CRISPR DSB induction at a conserved region of the endogenous PTEN gene. Data are averages from experiments performed in triplicate. Error bars represent SEM. * p<0.05, **** p<0.0001. All charts analyzed by two-way ANOVA with Fisher’s LSD. p-values should be considered nominal indices of significance. **d,** Graphical summary. The bowhead whale has evolved efficient and accurate DSB repair mediated by high levels of CIRBP and RPA2. This enhanced DNA repair may help the bowhead whale resist cancer despite its cells requiring fewer mutational hits for malignant transformation than human cells. Improved DNA repair rather than enhanced elimination of damaged cells through apoptosis or senescence may promote longevity in the bowhead whale.

We additionally observed that human fibroblasts with integrated NHEJ reporters displayed an increase in NHEJ efficiency when cultured at 33°C rather than 37°C. This increase in NHEJ efficiency was accompanied by an increase in CIRBP protein abundance in the cells cultured at 33°C (Extended Data Figure 8m).

### Effect of bowhead whale CIRBP on anchorage-independent cell growth

We investigated whether CIRBP overexpression affects the anchorage-independent growth of transformed cells. bwCIRBP was overexpressed in human fibroblasts containing SV40 LT, SV40 ST, H-Ras^G12V^, and hTERT. These cells showed delayed formation of colonies in soft agar compared to control cells (Extended Data Figure 11a, b). Importantly, there was no difference in the proliferation rate and cell viability between CIRBP-expressing and control cells in 2D culture, as assessed by MTT and trypan blue exclusion assays (Extended Data Figure 11c, d). Furthermore, there was no significant change in the expression of SV40 LT, H-Ras^G12V^, and cyclin-dependent kinase inhibitors (p16INK4a and p21) (Extended Data Figure 11e). Strikingly, we also observed a lower frequency of chromosomal aberrations in human transformed cells overexpressing bwCIRBP (Extended Data Figure 11f), suggesting that a possible explanation for the delay in colony formation is a reduction in chromosomal instability.^65^

### Role of RPA2 in bowhead whale DNA repair fidelity

LC-MS proteomics data, Western blots, and transcriptome analysis suggested increased abundance of the single-stranded DNA binding protein RPA2 in bowhead whale cells and tissues relative to other mammals (Figure 4a, Extended Data Figure 6c, e-g, Extended Data Figure 7a-d). RPA is a conserved heterotrimeric ssDNA-binding protein complex required for eukaryotic DNA replication, which plays a critical role in DNA repair and DNA damage signaling^64,65^. RPA deficiency increases the frequency of DSBs in human cells under both basal^66^ and stressed^67^ conditions. Conversely, RPA overexpression increases resistance to genotoxic insults^67–70^. RPA promotes NHEJ *in vitro*^71^ and protects ssDNA overhangs at DSBs^72,73^.

Treatment of cells transfected with CRISPR to induce DSBs with the small-molecule RPA DNA-binding inhibitor TDRL-505^74^ significantly increased indel rates in both bowhead and human fibroblasts (Figure 5a, b; Supplementary Table 5, 6). In bowhead, RPA inhibition also increased the frequency of larger deletions, although this difference did not reach significance (Figure 5b; Supplementary Table 6). Conversely, co-transfection of trimeric recombinant human RPA protein during CRISPR significantly decreased the frequency of mutated alleles in human cells without affecting CRISPR RNP transfection efficiency (Figure 5a, Supplementary Figure 4c).

In summary, these results suggest that increased abundance of CIRBP and RPA2 positively affects the fidelity of DSB repair, promoting genomic stability in the bowhead whale.

## Discussion

By studying a mammal capable of maintaining its health and avoiding death from cancer for over two centuries, we are offered a unique glimpse behind the curtain of a global evolutionary experiment that tested more mechanisms affecting cancer and aging than humans could hope to approach. Through experiments using primary fibroblasts from the bowhead whale, we experimentally determined genetic requirements for oncogenic transformation in the world’s longest living mammal and provide evidence that additional tumor suppressors are not the only solutions to Peto’s Paradox. Instead, we find that the bowhead whale solution lies upstream of tumor suppressor loss and is defined by a capacity for highly accurate and efficient DNA DSB repair, as well as improved mismatch repair. We also present evidence that two proteins highly expressed in the bowhead relative to other mammals, RPA2 and CIRBP, contribute to more efficient and accurate DSB repair.

CIRBP is highly abundant in the bowhead whale compared to most other mammals. We speculate that baleen whales evolved this high abundance of CIRBP as an adaptation to life in cold water and CIRBP was subsequently co-opted to facilitate genome maintenance. We show that CIRBP directly stimulates NHEJ, and in a purified system can promote NHEJ in the absence of Ku70/80. A potential mechanism by which CIRBP could promote end joining *in vivo* is through recruitment to damaged DNA in a manner enhanced by PAR and RNA, and by promoting the ligation of DNA ends either through direct interaction with DNA and/or through interaction with DNA repair proteins within a synaptic complex, tethering broken ends. Another potential mechanism by which CIRBP could promote ligation of DNA ends might involve liquid-liquid phase separation (LLPS). Recent evidence suggests that RNA-binding proteins can promote phase separation around DSB sites. For example, the FUS RNA-binding protein was found to maintain genomic stability and promote formation of droplet-like compartments in response to DNA damage^75^. Furthermore, it has been shown that long non-coding RNAs synthesized at DSBs are necessary to drive molecular crowding of 53BP1 into foci that exhibit LLPS condensate properties^76^. CIRBP has also been shown to undergo LLPS *in vitro*^77^. CIBRP may attract DNA into condensates through its affinity for nucleic acids, increasing rates of intermolecular interaction and ligation kinetics.

Cytosine deamination is an important endogenous source of age-related point mutations. Our mismatch repair assay measured repair of a G-T mismatch, one possible intermediate during cytosine deamination. The markedly improved efficiency of repair of this type of mismatch which we observed in the bowhead whale may hint at improved repair of this important source of age-related mutation.

While the source of the single-base insertion preference in the bowhead whale remains an open question, we speculate that this may be related to differential employment of bypass polymerases during repair synthesis. While mammals have over a dozen different DNA polymerases, many being specialized error-prone enzymes involved in lesion bypass, polymerase mu (Pol μ) was specifically found to be lost in the cetacean lineage^78^. Pol μ is one of the primary error-prone polymerases involved in NHEJ, along with polymerase lambda (Pol λ) and in some cases terminal deoxynucleotidyl transferase (TdT)^79^. Pol μ, in comparison with other repair polymerases such as Pol λ, has a unique propensity to “skip” ahead of 3’ unpaired bases, generating deletions^79,80^. The absence of Pol μ may lead to increased reliance on other repair polymerases, many of which are able to add a single terminal templated or untemplated nucleotide to a 3’ end and could explain the 1-bp insertion bias observed in the bowhead whale.

While we did not identify a signature of positive selection on amino acids in bowhead whale RPA2, we did identify a difference of possible functional significance in an N-terminal phosphorylation domain. Previous research has shown that hyperphosphorylation of multiple S/T residues within this domain drives important alterations in RPA2’s function. Bowhead whale RPA2 has more N-terminal S/T residues (13) than human RPA2 (9). It has been shown that RPA2 N-terminal hyperphosphorylation inhibits DNA end resection^81^, prevents localization to replication centers but not to damage sites^82^, and is specifically required for the maintenance of genome stability during replication stress^83^.

Prior work has identified positive selection in cancer-related genes such as *CXRC2, ADAMTS8, and ANXA1* in cetaceans, as well as cetacean-specific evolutionary changes to multiple FGF genes^84,85^. Expansion of the eukaryotic initiation factor 2, polyadenylate binding protein, and 60S ribosomal L10 gene families have also been identified in the genome of the bowhead whale, potentially implying substantial alterations to translational regulation^86^. An additional anti-cancer mechanism was recently reported for the bowhead whale in the form of a *CDKN2C* checkpoint gene duplication^87^. It has further been suggested that the low body temperature and low metabolic rate of the bowhead whale may also contribute in part to its extended lifespan^88^. Indeed, it is likely that numerous individual factors combine to modify cancer risk and produce the longest mammalian lifespan.

Improvements in DNA repair pathways have also been previously implicated in the evolution of mammalian longevity. Expression of DNA repair genes was found to be positively correlated with longevity in a transcriptomic study of 26 mammalian species.^89^ Interestingly, a prior study found higher levels of PAR synthesis and higher PARP1 recruitment to a DNA probe in vitro in the long-lived naked mole rat relative to the mouse^90^, which mirror cellular phenotypes we observed in the bowhead whale relative to human. While this study also noted higher NER activity in the naked mole rat relative to mouse, a subsequent study using additional rodent species found that the efficiency of DSB repair correlates more strongly with longevity across rodent species^91^.

One potential drawback of a very accurate DNA repair system could be a reduction in standing genetic variation and thus a slower rate of evolution of new traits. However, species living in safe and stable environments have less evolutionary pressure to rapidly evolve new adaptations. A genome analysis of long-lived rockfishes living in deep ocean revealed positive selection in DNA repair pathways^92^. Interestingly, a recent analysis of germline mutations in baleen whales based on analysis of pedigrees concluded that germline mutation frequencies are similar to those in primates, in contrast to prior studies finding reduced germline mutation rates in whales^93^. Thus, it appears that germline and somatic mutation rates are not inherently linked and could respond to selection independently.

Why would improved DNA repair have evolved in the bowhead whale, as opposed to the increased tumor suppressor copy number and elevated apoptotic response found in the elephant and often proposed as a solution to Peto’s Paradox? One possible explanation is that tumor suppressors, apoptosis, and senescence all appear to pose costs to the organism and force tradeoffs between cancer and cell depletion leading to age-related degeneration. Simply shifting the balance from apoptosis/senescence to survival and repair could be detrimental if not also coupled with increased fidelity, as evidenced by the frequent upregulation of DNA repair pathways in cancer cells^94^.

However, evolutionary improvements that couple high efficiency with high fidelity, as found in the bowhead whale, would promote long-term tissue function and maintenance at both the cellular and genomic levels. Maintenance of genome stability would reduce cancer risk and, as suggested by a rapidly growing body of evidence implicating age-related somatic mosaicism as a ubiquitous feature and functional driver of aging^95–99^, likely protect against numerous other aspects of age-related decline. Thus, the lower accuracy and efficiency of DNA repair observed in mammals with shorter lifespans may simply reflect the absence of sufficient selective pressure^88^. Indeed, there is little selective advantage of DNA repair capacity to last far beyond the age of first reproduction.

There are currently no approved therapeutics which aim to bolster DNA repair for the prevention of cancer or age-related decline, and it has been suggested that DNA repair would be difficult or impossible to improve^100^. However, the bowhead whale provides evidence that this notion is incorrect. Expression of bwCIRBP in human cells improves the efficiency and accuracy of DSB repair. Therapeutics based on the evolutionary strategy of the bowhead whale, including trying to increase activity or abundance of proteins like CIRBP or RPA2, could one day enable the treatment of genome instability as a modifiable disease risk factor (for discussion of therapeutic hypothermia see Supplementary Note). Improving DNA repair could be especially important for patients with increased genetic predisposition for cancer, or more generally, for aging populations at increased risk for cancer development.

## Acknowledgements

We would like to thank the researchers at the North Slope Borough Department of Wildlife Management, the Alaska Eskimo Whaling Commission, and the Iñupiaq community of Barrow for generously sharing bowhead whale samples, time, resources, skill, and knowledge, and without whom the above work would not have been possible. We would like to give a special thanks to John Craighead “Craig” George, whose pioneering field work established the remarkable longevity of the bowhead whale, and whose kind collaboration and insights helped initiate this project, but who sadly was unable to see its completion. We would like to thank Carlo Maley for suggesting collaboration between Schiffman and Gorbunova groups during a memorable meeting in Arcachon. We would like to thank Aaron Rogers, Mallory Wilmot, and Ryan Kennington for technical support. We would like to thank Tara Harrison, Leigh Duke, the Exotic Species Research Alliance, and Georgia Aquarium for facilitating the collection of the bottlenose dolphin sample. We thank the Marine Mammal Care Center Los Angeles for collecting samples from California sea lions. We would like to thank San Diego Zoo Wildlife Alliance for providing cells from hippopotamus, common dolphin, and humpback whale. Experiments on in vitro ligation were supported by the French National Research Agency. Experiments on PAR-binding were supported by National Institutes of Health R01GM104135 to A.K.L.L. We thank V.K. Thomas in the URMC Center for Advanced Light Microscopy and Nanoscopy (RRID:SCR_023177) for images acquired on the Abberior 3D STED instrument (S10 OD023440). This work was supported by grants from US National Institutes on Aging to VNG, ZZ, JV, AS and VG, and by an award from The Milky Way Research Foundation to VG.

## Author contributions

VG, AS, DF, MZ designed research; DF conducted molecular cloning, lentivirus production, immunofluorescence experiments, PFGE, senescence experiments, cell survival experiments, DNA repair assays, micronuclei assays, PARP experiments, CIRBP experiments, and assessed DNA repair fidelity using NHEJ reporter with help from SAB, AP, EH, AW, NM, ML; MZ conducted tumor suppressor CRISPR experiments, RPA experiments, HPRT assays, comet assays, micronuclei analysis, EMSA, and assessed DNA repair fidelity using CRISPR with help from SAB, MS, NH; XT analyzed tumorigenicity; DF and XT conducted cell growth curves and telomere experiments; VV analyzed chromosomal aberrations; AK conducted MMR assay; YZ, CC, Zhihui Zhang and AG assisted with mouse tumor studies; MM and SF conducted NHEJ ligation assay; JCG, TLS, MZ, DF collected bowhead specimens; LP, EG, LZ, and GG performed tumor xenograft sequencing and analysis with help from MZ; JH, AM, SS, and JV performed SMM-Seq of ENU-treated cells with help from JA; ZW assisted with LC-MS and micronuclei; JG assisted with micronuclei and the alkaline comet assay; JM assisted with the HPRT assay; JG performed STED imaging; MH, RM, and GT did LC-MS of liver tissue; LMT did nuclear extractions from liver; MZ, GT performed cell proteomics; JYL and Zhizhong Zheng analyzed RNAseq with help from DF; HL, YC, AKLL performed PAR-binding assays; VG, Zhengdong Zhang, JV contributed data analysis and conceptualization; VG and AS obtained funding and supervised the study; MZ, DF, AS and VG wrote the manuscript with input from all authors.

## Methods

### Reagents

Detailed information on reagents, such as antibodies and sequences of primers, probes, CRISPR guides, and siRNAs, is provided in Supplementary Methods.

### Animal experiments

All animal experiments were approved and performed under pre-approved protocols and in accordance with guidelines set by the University of Rochester Committee on Animal Resources (UCAR).

### Whale sample collection

Bowhead whale tissues were obtained from adult bowhead whales (*Balaena mysticetus*) captured during 2014 and 2018 Iñupiaq subsistence harvests in Barrow (Utqiaġvik), AK, in collaboration with the North Slope Borough Department of Wildlife Management (NSB DWM) and Alaska Eskimo Whaling Commission after signing a Memorandum of Understanding (September 2014 and March 2021). Tissues were sampled immediately after bowhead whales were brought ashore, after permission to sample was given by the whaling captain, and explants kept in culture medium on ice or at 4°C through initial processing and shipping until arrival at the University of Rochester (UR) for primary fibroblast isolation from skin and lung. Transfer of bowhead whale samples from NSB DWM to UR was under NOAA/NMFS permit 21386.

### Primary cell cultures

Primary skin fibroblasts were isolated from skin (dermal) tissues as previously described.^101^ Briefly, skin tissues were shaved and cleaned with 70% ethanol. Tissues were minced with a scalpel and incubated in DMEM/F-12 medium (ThermoFisher) with Liberase™ (Sigma) at 37°C on a stirrer for 15-90 min. Tissues were then washed and plated in DMEM/F-12 medium containing 12% fetal bovine serum (GIBCO) and Antibiotic-Antimycotic (GIBCO). All subsequent maintenance culture for fibroblasts from bowhead and other species was in EMEM (ATCC) supplemented with 12% fetal bovine serum (GIBCO), 100 units/mL penicillin, and 100 mg/mL streptomycin (GIBCO). All primary cells were cultured at 37°C with 5% CO_2_ and 3% O_2_ except bowhead whale cells, which were cultured at 33°C with 5% CO_2_ and 3% O_2_ based on published field measurements of bowhead body temperature, which measured a core temperature of 33.8 °C and a range of lower temperatures in muscle and peripheral tissue.^102,103^ Prior to beginning experiments with bowhead whale fibroblasts, optimal growth and viability conditions were empirically determined through testing of alternative temperatures, serum concentrations, and cell culture additives, with optimal culture medium found to be the same for bowhead and other species. Following isolation, low population-doubling (PD) primary cultures were preserved in liquid nitrogen, and PD was continually tracked and recorded during subsequent use for experiments.

Established, primary fibroblasts from mammals were obtained from San Diego Zoo Wildlife Alliance (hippopotamus, common dolphin, and humpback whale) or generated at Huntsman Cancer Institute from bottlenose dolphin tissues collected by Georgia Aquarium through Tara Harrison (Exotic Species Cancer Research Alliance) and California sea lion tissues collected by the Marine Mammal Care Center Los Angeles under Institutional Animal Care and Use Committee oversight and National Marine Fisheries Service permit number 21636.

### Soft agar assay

Fibroblast culture medium as described above was prepared at 2X concentration using 2X EMEM (Lonza). To prepare the bottom layer of agar plates, 2X medium was mixed with a sterile autoclaved solution of 1.2% Noble Agar (Difco) at a 1:1 volumetric ratio, and 3 mL of 1X medium/0.6% agar was pipetted into each 6-cm cell culture dish and allowed to solidify at room temperature in a tissue culture hood. To plate cells into the upper layer of soft agar, cells were harvested and washed, and immediately prior to plating were resuspended in 2X medium at 20,000 cells/1.5 mL and diluted twofold in 0.8% Noble Agar pre-equilibrated to 37°C. The cells in 0.4% agar/1X medium were pipetted gently to ensure a homogeneous single cell suspension, and 3 mL (20,000 cells) per 6 cm dish were layered on top of the solidified lower layer. After solidifying in tissue culture hoods for 20-30 min, additional medium was added to ensure the agar layers were submerged, and dishes were moved into cell culture incubators. Fresh medium was added onto the agar every 3 days. 4 weeks after plating, viable colonies were stained overnight with nitro blue tetrazolium chloride (Thermo Fisher) as previously described.^40^ All cell lines were plated in triplicate.

Images of colonies in soft agar were captured using the ChemiDoc MP Imaging System (Bio-Rad). Colony quantification was performed using ImageJ software (NIH). Initially, images were converted to 8-bit format. Subsequently, the threshold function was adjusted to eliminate any red pixels highlighting non-colony objects. Following threshold adjustment, images were converted to binary. Colony counting was executed using the ‘Analyze Particles’ function with the following parameters: Size (pixel^2) = 1 to infinity; Circularity = 0.5 to 1.

### Mouse xenograft assay

NIH-III nude mice (Crl:NIH-Lystbg-J Foxn1nuBtkxid) were purchased from Charles River Laboratories Inc. (Wilmington, MA, USA). Seven-week-old female mice were used to establish xenografts. For each injection, 2 × 10^6^ cells were harvested and resuspended in 100 μl of ice-cold 20% matrigel (BD Bioscience, Franklin Lakes, NJ) in PBS (Gibco). Mice were anesthetized with isoflurane gas, and 100 μl solution per injection was injected subcutaneously into the right and left flanks of each mouse with a 22 gauge needle. 3 mice were injected bilaterally, for a total of 6 injections, per cell line tested. Tumor length and width were measured and recorded every 3-4 days. Mice were euthanized after reaching a predetermined humane tumor burden endpoint of a maximum tumor dimension of 20mm in diameter, determined by the longest dimension of the mouse’s largest tumor. For mice that did not reach tumor burden endpoints, experiments were terminated, and mice euthanized after a maximum of 60 days.

Euthanized mice were photographed, and tumors were excised, photographed, and weighed to determine the mass of each tumor. Sections of each tumor were frozen at −80°C and preserved in formalin.

### MTT Assay

Cell metabolic activity was determined using Thiazolyl Blue Tetrazolium Bromide (MTT) (Sigma). Cells were seeded in 24-well plates at a density of 20,000 cells per well one day before the assay. An MTT solution in PBS was added to the growth medium to achieve a final concentration of 0.5 mg/mL, and cells were then incubated for 4 hours in a CO2 incubator. Following incubation, the growth medium was discarded, and 0.5 mL of DMSO was added to each well to solubilize the purple formazan crystals completely. The plate was further incubated until the crystals were fully dissolved. Spectrophotometric absorbance of the samples was measured at a wavelength of 570 nm using a Tecan Spark 20M plate reader.

### Telomere lengths

Telomere length was analyzed by Southern blot using the TRF method. Genomic DNA was extracted from cultured fibroblasts at different population doublings, digested with a mixture of AluI, HaeIII, RsaI, and HinfI restriction enzymes that do not cut within telomeric repeat sequences, separated using pulsed-field gel electrophoresis, and hybridized with a radiolabeled oligonucleotide containing telomeric sequence (TTAGGG)_4_. Pulsed field gels were run using a CHEF-DR II apparatus (Bio-Rad) for 22h at a constant 45 V, using ramped pulse times from 1 to 10 s.

### Telomeric repeat amplification protocol

Telomeric repeat amplification protocol assay was performed using the TRAPeze kit (Chemicon, Temecula, CA, USA) according to manufacturer instructions. Briefly, in the first step of the TRAP assay, radiolabeled substrate oligonucleotide is added to 0.5 μg of protein extract. If telomerase is present and active, telomeric repeats (GGTTAG) are added to the 3′ end of the oligonucleotide. In the second step, extended products are amplified by PCR. Telomerase extends the oligonucleotide by multiples of 6 bp, generating a ladder of products of increasing length. A human cancer cell line overexpressing telomerase as well as rodent cells were used as a positive control.

### CRISPR ribonucleoprotein transfection

CRISPR RNP complexes were formed in vitro by incubating Alt-R™ S.p.Cas9 Nuclease V3 (Integrated DNA technologies) with tracRNA annealed to target-specific crRNA (Integrated DNA Technologies) according to manufacturer instructions. For generation of tumor suppressor knockouts, 3 RNP complexes with crRNAs targeting different sites in a single target gene were combined and Alt-R Cas9 Electroporation Enhancer (Integrated DNA Technologies) was added to transfection mixes prior to electroporation. For comparative analysis of repair fidelity, 3 μg of pmaxGFP plasmid (Lonza) was added to transfection mixes to monitor transfection efficiency. Cells were trypsinized and washed with PBS, and 1 x 10^6^ cells were resuspended in 100 μL of NHDF Nucleofector Solution (Lonza). The cell suspension was then combined with the CRISPR transfection solution and gently mixed prior to electroporation on an Amaxa Nucleofector 2b (Lonza) using program U-23. For RPA inhibition, human and bowhead fibroblasts were treated with 50 uM TDRL-505 (Sigma) (diluted 1000x into culture medium from 50mM stock solution in DMSO) for 3 hours prior to CRISPR transfection and kept in medium with 50 uM TDRL-505 for 18 hours after transfection. For RPA co-transfection, 1 μg of recombinant human RPA heterotrimer (Enzymax) was added to CRISPR transfection solution immediately prior to electroporation.

### Isolation of clonal cell colonies and screening for tumor suppressor knockout

Following CRISPR transfection, cells were plated at low density in 15 cm dishes to allow for the formation of isolated colonies. Once clonal colonies of sufficient size had formed, positions of well-isolated colonies were visually marked on the bottom of the cell culture dish while under a microscope using a marker. Dishes were aspirated and washed with PBS. Forceps were used to dip PYREX® 8×8 mm glass cloning cylinders in adhesive Dow Corning® high-vacuum silicone grease (Millipore Sigma) and one glass cylinder was secured to the dish over each marked colony. 150 µL of trypsin was added to each cylinder and returned to the incubator. When cells had rounded up from the plate, the trypsin in each cylinder was pipetted to detach cells and each colony was added to a separate well in a 6-cm culture dish containing culture medium. After colonies were expanded and split into 2 wells per colony, one well was harvested for Western blot screening for absence of target proteins, while the remaining well was kept for further experiments.

### Luciferase reporter assays for knockout verification

For p53 activity measurement, 1 x 10^6^ cells of control (WT) and clonally isolated p53 KO cell lines were electroporated with 3 µg p53 firefly luciferase reporter plasmid pp53-TA-Luc (Clontech/Takara) and 0.3 μg renilla luciferase control plasmid pRL-CMV (Promega) on an Amaxa Nucleofector 2b (Lonza). 24h later, cells were treated with 200 μM etoposide (Sigma) to induce p53 activity. 24h following etoposide treatment, cells were harvested, and luciferase activity of cell lysates was measured using the Dual-Luciferase Reporter Assay System (Promega) in a GloMax 20/20 Luminometer (Promega) according to manufacturer instructions.

For Rb activity measurement, 2 different reporters were tested in parallel: pE2F-TA-Luc (Clontech/Takara) to measure E2F transcriptional activity (repressed by Rb), and pRb-TA-Luc (Clontech/Takara) (promoter element directly suppressed by Rb). 1 x 10^6^ cells of control (WT) and clonally isolated Rb KO cell lines were electroporated with 3 µg of either pE2F-TA-luc or pRb-TA-luc and 0.3 ug renilla luciferase plasmid on an Amaxa Nucleofector 2b (Lonza). Following transfection, cells were grown in complete medium for 24h followed by serum-free medium for 24h. Cells were then harvested, and luciferase activity measured as described above.

### Error-corrected sequencing by SMM-seq of ENU-mutated cells

Skin fibroblasts from mouse, cow, human and whale were isolated and cultured as described before. Confluent cells were treated with 20mg/ml ENU overnight. Then cells were split 1:4 and grown until confluence for harvest.

Genomic DNA (gDNA) was isolated from frozen cell pellets using the Quick DNA/RNA Microprep Plus Kit (Zymo D7005). Three hundred ng were used for library preparation as described before^104^: in brief, DNA was enzymatically fragmented, treated for end repair before adapter ligation and exonuclease treatment. A size selection step was performed using a 1.5% cassette on a PippinHT machine prior pulse rolling circle amplification (RCA) and indexing PCR. Library quality was determined with a Tape Station (Agilent) and quantified with Qubit (Thermo Fisher). All libraries were sequenced by Novogene Corp. Inc (CA) on an Illumina platform.

Sequencing analysis and mutation calling was performed as described^104^. After trimming and alignment to the corresponding reference genome, additional filtering was performed to distinguish germline mutations from somatic mutations. Graphs were generated and statistical testing was performed using GraphPad Prism.

### Next-generation sequencing of CRISPR repair products

72h after transfection, cells were harvested, and genomic DNA was isolated with the Wizard Genomic DNA Purification Kit (Promega). DNA concentration was measured on a Nanodrop spectrophotometer and 100 ng of DNA per sample was PCR-amplified with KAPA2G Robust HotStart ReadyMix (Roche) based on findings of low PCR bias for KAPA polymerase^105,106^. Primers targeted a conserved region surrounding *PTEN* exon 1 (Extended Data Figure 4a). PCR was performed according to manufacturer instructions, with an annealing temperature of 66°C for 30 cycles. To purify samples for next-generation sequencing, PCR products were electrophoresed on a 0.8% agarose gel and post-stained with SYBR Gold Nucleic Acid Gel Stain (Thermo Fisher). Gels were visualized on a blue light tray (BioRad) to minimize damage to DNA. A gel slice for each lane was excised using a scalpel, and each slice was cut to include the region ranging from just above the prominent *PTEN* PCR band down to and including the “primer dimer” region to ensure inclusion of any deletion alleles. DNA was extracted from gel slices using the QiaQuick Gel Extraction Kit (Qiagen), and triplicate PCR reaction eluates per sample were pooled for sequencing. Sample concentrations were measured by Nanodrop and adjusted as necessary prior to submission for 2×250 bp paired-end Illumina MiSeq sequencing with target depth of >40,000 reads/sample (Genewiz).

### Analysis of CRISPR NGS data

FASTQ files from each sequenced sample were analyzed with both CRISPResso2,^107^ which uses an alignment-based algorithm, and CRISPRPic,^108^ which uses a kmer-based algorithm. CRISPResso2 was run using the following parameters: window size = 30, maximum paired-end overlap = 500, bp excluded from left and right ends = 15, minimum alignment score = 50, minimum identity score = 50, plot window size = 20. For CRISPRPic analysis, SeqPrep^109^ was used to merge overlapping read pairs and trim adapter sequences. CRISPRPic was run on merged FASTQ sequences for each sample with the following parameters: index size = 8, window size = 30.

### HPRT mutation assay

For the HPRT mutation assay, cells used were low-passage primary dermal fibroblasts from multiple species that were known to originate from male animals, to ensure single copy-number of the X-linked *HPRT* gene. Each species was tested with 3 different cell lines from 3 individual animals. The bowhead *HPRT* coding sequence was BLASTed against bowhead genome scaffolds^17^ and neighboring gene sequences were analyzed to confirm mammal-typical localization of *HPRT* on the bowhead X-chromosome. Cells were cultured in standard fibroblast growth medium, but with FBS being replaced with dialyzed FBS (Omega Scientific, Inc.) and supplemented with Fibroblast Growth Kit Serum-Free (Lonza) to improve growth and viability in dialyzed FBS. Dialyzed FBS was found in optimization experiments to be necessary for efficient 6-thioguanine selection. Prior to mutagenesis, cells were cultured for 7 days in medium containing HAT Supplement (Gibco) followed by 4 days in HT Supplement (Gibco) to eliminate any pre-existing HPRT mutants. To induce mutation, cells were incubated for 3 hours in serum-free MEM containing 150 µg/mL N-ethyl-N-nitrosourea (ENU) (Sigma), or exposed to 2 Gy γ-irradiation. Cells were then maintained in ENU-free medium for 9 days to allow mutations to establish and existing HPRT to degrade. 1 x 10^6^ cells from each cell line were harvested and plated in dialyzed FBS medium containing 5 µg/mL 6-thioguanine (Chem-Impex), in parallel with 1 x 10^6^ untreated control cells for each cell line. Cells were plated at a density of 1 x 10^5^ cells per 15-cm dish to allow for efficient selection and colony separation, and to prevent potential “metabolic cooperation“^45^. In tandem, for each cell line 200 cells from untreated and control conditions were plated in triplicate 10-cm dishes in non-selective medium to calculate plating efficiency. After 3 weeks of growth, surviving colonies were fixed and stained with a crystal violet/glutaraldehyde solution as previously described^110^. Colonies were counted, and HPRT mutation rate was calculated as plating-efficiency adjusted number of HPRT-negative colonies containing >50 cells. Appropriate concentrations of ENU and 6-TG, as well as optimal plating densities and growth conditions, were determined prior to the experiment described above through optimization and dose titration experiments.

### Digital droplet PCR measurement of CRISPR cleavage rate

A ddPCR assay similar to a previously published method^111^ was used for time-course quantification of CRISPR DSB induction across species. qPCR primers at conserved sites flanking the guide RNA target site in the *PTEN* gene were designed such that cleavage would prevent PCR amplification. As an internal copy number reference control, a second set of previously validated qPCR primers targeting an ultraconserved element present in all mammals as a single copy per genome (UCE.359) was designed based on published sequences^112^. To allow for multiplexing and copy number normalization of *PTEN* within each ddPCR reaction, 5‘fluorescent hydrolysis probes (FAM for *PTEN* and HEX for UCE.359) targeting conserved sequences were designed, with 3‘ Iowa Black® and internal ZEN™ quenchers (Integrated DNA Technologies). All primers and probes were checked for specificity by BLAST against each species’ genome ^112^. Fibroblasts were transfected with *PTEN* CRISPR RNP as described in “Next-generation sequencing of CRISPR repair products” and returned to cell culture incubators. At the indicated times post-transfection, cells were harvested, flash frozen, and genomic DNA was isolated with the Wizard Genomic DNA Purification Kit (Promega). During isolation, newly lysed cells were treated with Proteinase K and RNase A for 30 minutes each at 37°C to minimize the possibility of residual CRISPR RNP activity. DNA concentration was measured on a Nanodrop spectrophotometer, and genomic DNA was pre-digested with BamHI-HF (NEB) and XhoI (NEB), which do not cut within target amplicons, to maximize PCR efficiency and distribution across droplets. 15 ng of genomic DNA per sample was added to duplicate PCR reactions using the ddPCR™ Supermix for Probes (No dUTP) master mix (Bio-Rad). Droplets were prepared and measured according to manufacturer instructions. Briefly, each 20 µL reaction was mixed with 70 µL Droplet Generation Oil for Probes (Bio-Rad) and droplets were formed in a QX100 Droplet Generator (Bio-Rad). 40 µL of droplets per reaction were transferred to 96-well PCR plates and sealed with a PX1 PCR Plate Sealer (Bio-Rad). The sealed plates were then subjected to PCR using a pre-optimized cycling protocol. Following PCR, the plates were loaded into a QX100™ Droplet Reader (Bio-Rad) and each droplet measured on both FAM and HEX channels. *PTEN* copy number normalized to UCE.359 reference copy number within each well was determined with QuantaSoft™ software (Bio-Rad). For each species, positive/negative gates in mock-transfected control samples were adjusted as necessary to compensate for differences in multiplex PCR efficiency/specificity and “rain” droplets between species and bring normalized *PTEN* copy number closer to 1. The control gates were then applied across all samples/time points within the same species and used for *PTEN* copy number calculation.

### Flow cytometric measurement of CRISPR RNP transfection efficiency

CRISPR RNP transfections were performed as described above, but with ATTO-550 fluorescently labeled tracRNA (Integrated DNA Technologies). At 0h and 24h post-transfection, cells were harvested, pelleted, and analyzed by flow cytometry on a CytoFlex S Flow Cytometer (Beckman Coulter). Gain and ATTO-550 positive gates were set based on mock-transfected control cells included in each experiment.

### Senescence-associated β-galactosidase (SA-β-gal) staining

SA-β-gal staining was performed as previously described ^113,114^. Cells were washed twice with PBS and fixed in a solution containing 2% formaldehyde and 0.2% glutaraldehyde in PBS for 5 min at room temperature. After fixation, cells were immediately washed twice with PBS and stained in a solution containing 1 mg/mL 5-bromo-4-chloro-3-indolyl P3-D-galactoside (X-Gal), 40 mM citric acid/sodium phosphate buffer, pH 6.0, 5 mM potassium ferrocyanide, 5 mM potassium ferricyanide, 150 mM NaCl, and 2 mM MgCl_2_. Plates were incubated at 37°C for 16 h without CO_2_. Colorimetric images were taken from different areas of each plate and quantified.

### Cell survival assay

Percentage of live cells was quantified using the Annexin-V FLUOS Staining Kit (Roche) and Annexin V Apoptosis Kit [FITC] (Novus Biologicals) following the manufacturer’s instructions. After staining, cells were analyzed on a CytoFlex S flow cytometer (Beckman Coulter). Where indicated cell viability was assessed using a trypan blue exclusion assay. All cells (both floated and attached to the culture dish) were collected into the same tube, centrifuged, and resuspended in PBS. The cells were then mixed in a 1:1 ratio with 0.4% trypan blue solution, and approximately 3 minutes later, the percentage of dead cells was assessed using the Countess 3FL instrument (ThermoFisher) according to the manufacturer’s instructions.

### p53 activity

To test p53 activity in cultured primary fibroblasts, 150,000 cells were seeded in 6-well plates 1 day before transfection with 1 μg pp53-TA-Luc vector (Clontech) and 0.015 μg pRL-CMV-Renilla (Promega) to normalize for transfection efficiency. Transfections were performed using PEI MAX Transfection Grade Linear Polyethylenimine Hydrochloride (MW 40,000) (Polysciences) according to manufacturer instructions. 24h after transfections cells were lysed using 50μl passive lysis buffer (Promega) per 10^5^ cells and flash frozen/thawed two times in liquid nitrogen and a 37°C water bath. Luciferase assays were performed using the Dual-Luciferase Reporter Assay System (Promega) and program DLR-2-INJ on a Glomax 20/20 Luminometer (Promega) with 20μl cell extract as the input.

### Generation of NHEJ and HR reporter cell lines

NHEJ and HR reporter constructs ^46^ were digested with NheI restriction enzyme and purified with the QIAEX II gel extraction kit (QIAGEN). The same plasmid DNA preparation was used for generating all reporter cell lines of the studied species. Cells PD < 15 were recovered from liquid nitrogen and passaged once before the integration of the constructs. 0.25 µg of linearized NHEJ and HR constructs were electroporated into one million cells for each cell line. Two days after transfection, media was refreshed, and G418 was applied to select stable integrant clones. Triplicates of each reporter in each cell line were prepared to obtain an adequate number of stable clones. Clones from triplicate plates were pooled to get at least 50 clones per reporter per cell line.

### DSB repair assays and flow cytometry analysis

DSB repair assays were performed as previously described^115^. Briefly, growing cells were co-transfected with 3 µg of plasmid encoding I-SceI endonuclease and 0.03 µg of plasmid encoding DsRed. The same batch of I-SceI and DsRed mixture was used throughout all species to avoid batch-to-batch variation. To test the effect of CIRBP on DSB repair, 3 µg of CIRBP plasmids were co-transfected with I-SceI and DsRed plasmids. Three days after transfection, the numbers of GFP+ and DsRed+ cells were determined by flow cytometry on a CytoFlex S Flow Cytometer (Beckman Coulter). For gating strategy see Supplementary Figure 5 in Supplementary Information. For each sample, a minimum of 50,000 cells was analyzed. DSB repair efficiency was calculated by dividing the number of GFP+ cells by the number of DsRed+ cells.

For NHEJ knockdown experiments, bowhead whale cells containing the NHEJ reporter were transfected with 120 pmol of anti-bwCIRBP or control siRNAs (Dharmacon) three days before I-SceI/DsRed transfections using an Amaxa Nucleofector (U-023 program). For HR knockdown experiments, bowhead whale cells containing the HR reporter were transfected twice every three days with a final concentration of 10 nM anti-bwCIRBP or negative control siRNAs (Silencer Select, Thermo Fisher) using Lipofectamine RNAiMAX transfection reagent (Thermo Fisher) following the manufacturer’s instructions. Cells were further transfected with I-SceI/DsRed plasmids using a 4D-Nucleofector (P2 solution, DS150 program). The efficiency of knockdown was determined by Western blot.

For the extrachromosomal assay and fidelity analysis, NHEJ reporter plasmid was digested with I-Sce1 for 6h and purified using a QIAEX II Gel Extraction Kit (QIAGEN). Exponentially growing cells were transfected using an Amaxa nucleofector with the U-023 program. In a typical reaction, 10^6^ cells were transfected with 0.25 µg of predigested NHEJ reporter substrate along with 0.025 µg of DsRed to serve as a transfection control. 72h after transfection, cells were harvested and analyzed by flow cytometry on a BD LSR II instrument. At least 20,000 cells were collected for each sample. Immediately after FACS, genomic DNA was isolated from cells using the QIAGEN Blood & Tissue kit. DSB repair sites in the NHEJ construct were amplified by PCR using Phusion polymerase (NEB), cloned using the TOPO Blunt cloning kit (NEB), and sent for Sanger sequencing. At least 100 sequenced clones were aligned and analyzed using the ApE software.

### Western blotting

All antibodies were checked for conservation of the target epitope in the protein sequence of each included species, and only those targeting regions conserved across these species were used. For a limited number of proteins where the available antibodies with specific epitope information disclosed did not target conserved regions, we selected antibodies based on demonstrated reactivity across a broad range of mammal species and always confirmed these results with multiple antibodies. Information on antibodies is provided in Supplementary Methods.

Exponentially growing cells were harvested with trypsin and counted, and 10^6^ cells were resuspended in 100 µL of PBS containing protease inhibitors. 100µL of 2x Laemmli buffer (Bio-Rad) was added, and samples were boiled at 95°C for 10 minutes. Samples were separated with 4-20% gradient SDS-PAGE, transferred to a PVDF membrane, and blocked in 5% milk-TBS-T for 2 hours at room temperature. Membranes were incubated overnight at +4°C with primary antibodies in 5% milk-TBS-T. After 3 washes for 10 minutes with TBS-T, membranes were incubated for 1 hour at room temperature with secondary antibodies conjugated with HRP or a fluorophore. After 3 washes with TBS-T signal was developed for HRP secondaries with Clarity Western ECL Substrate (Bio-Rad). CIRBP and RPA2 expression were each measured with 3 different antibodies targeting conserved epitopes (Extended Data Figure 7b, c).

For detecting chromatin-bound proteins, cells were lysed in 1 mL of CSK buffer (10 mM Pipes pH 6.8, 100 mM NaCl, 300 mM sucrose, 3 mM MgCl2, 1 mM EGTA, 0.2% Triton X-100) or CSK+R buffer (10 mM Pipes pH 6.8, 100 mM NaCl, 300 mM sucrose, 3 mM MgCl2, 1 mM EGTA, 0.2% Triton X-100, and 0.3 mg/mL RNAse A) at +4°C for 30 min with gentle rotation. Samples were centrifuged for 10 min at 10,000 × g at 4°C, and the supernatant was discarded. Pellets were washed twice with 1 mL of CSK/CSK+R buffer, resuspended in PBS, and an equal volume of 2x Laemmli buffer (Bio-Rad) was added. Samples were boiled at 95°C for 10 minutes and subjected to Western blotting as described above.

For analyzing CIRBP expression in mice and bowhead whale tissues, tissues were pulverized using the cell crusher. For each 5 mg of tissue, 300 µL of 4x Laemmli buffer (Bio-Rad) was added, samples were extensively vortexed, and boiled at 95°C with 1000 rpm for 10 minutes.

### Expression and purification of human and bowhead whale CIRBP protein

N-terminal histidine-tagged (6xHis) CIRBP was cloned into a pET11a expression vector. The plasmid was transformed into Rosetta gami B (DE3) pLysS competent E. coli for protein expression. Bacteria were grown at 37oC to an optical density (OD600) of 2.0 and protein expression was induced by adding 0.4mM isopropyl β-d-1-thiogalactopyranoside (IPTG) for 20 h at 23oC. Bacteria were collected by centrifugation and pellets were flash frozen on liquid nitrogen and stored at −80oC. In Bacteria were resuspended in lysis buffer consisting of 50mM Tris pH 7.5, 2.0M NaCl, 50mM imidazole, 10mg lysozyme, 0.1% Triton X-100, 1mM DTT and protease inhibitors. The bacterial pellets were sonicated, rotated for 1 h at 4oC, and sonicated again. The bacterial lysate was clarified by centrifugation at 22,000g for 20 min at 4oC and the supernatant passed through a 0.45µm filter. The clarified lysate was purified using Ni-NTA agarose beads (Qiagen) washed with 20 column volumes of water and 20 column volumes of buffer containing 50mM Tris pH 7.5, 2.0M NaCl, 1mM DTT, and 50mM imidazole (Wash buffer 1). The lysate was placed onto the washed beads and transferred to a 50mL conical tube and rotated 3 hr at 4oC. The suspended beads were pelleted by centrifugation and washed with 40 column volumes wash buffer 1 and 10 column volumes with buffer containing 50mM Tris pH 7.5, 150mM NaCl, 1mM DTT, and 50mM imidazole. CIRBP was eluted by adding 5 column volumes of buffer containing 50mM Tris pH 7.5, 150mM NaCl, 1mM DTT, and 500mM imidazole and rotated the conical tube for 15 minutes at 4oC. The supernatant was collected by centrifugation and filtered before adding 5% glycerol. The protein was aliquoted, and flash frozen on liquid nitrogen and stored at −80oC.

### NHEJ ligation in vitro assay

The assay was performed essentially as described ^116,117^. Reaction mixtures (10 μl) contained 20 mM Tris-HCl (pH 7.5), 8 mM MgCl2, 0.1 mM ATP, 2 mM DTT, 0.1 M KCl, 2% Glycerol, 4% PEG 8000, 1 nM linearized pUC19 (with cohesive ends via XbaI; 17.3 ng), 10 nM XRCC4/Ligase IV complex, and 0.5 or 1 μM human CIRBP. When indicated, reaction mixtures also contained 10 nM Ku70/80 heterodimer, 1 μM XLF dimer, or 1 μM PAXX dimer. The reaction mixtures were incubated for 1 hr at 30°C, followed by the addition of 2 μl of Gel Loading Dye, Purple (6X) (NEB), and incubation for 5 min at 65°C. Subsequently, 4 μl of each sample was loaded onto a 0.7% agarose gel and subjected to gel electrophoresis (50 V, 50 min). The gel was stained with ethidium bromide, and DNA bands were visualized using a ChemiDoc MP (Bio-Rad).

### PARP activity

PARP activity was measured in cell nuclear extracts with the PARP Universal Colorimetric Assay Kit (Trevigen) according to the manufacturer’s instructions. Nuclear extracts were prepared using EpiQuik Nuclear Extraction Kit (EpigenTek) following manufacturer protocol. 2.5µg of total nuclear extract was added to measure PARP activity.

For measurement of PARylation efficiency, cells were treated with 400µM H_2_O_2_ for 15 and 30 min or subjected to 20 Gy γ-radiation. At the end of incubation, cells were placed on ice, washed once with PBS, and lysed directly on a plate with 2x Laemmli buffer. Samples were boiled for 10min at 95°C and processed by Western Blot.

### Preparation of fluorescent ligands, binding assays and fluorescence polarization measurements

Poly(ADP-ribose) (PAR) oligomers of different lengths (PAR_8_, PAR_16_, and PAR_28_) were synthesized, purified, fractionated, and labeled with Alexa Fluor 488 (AF488) dye at the 1’’ end, following as described previously^118,119^.

To investigate the binding of human and bowhead whale CIRBPs to the fluorescently labeled PAR and RNA oligomers, titration experiments were conducted. CIRBP proteins were 4:3 serially diluted and titrated into solutions containing a fixed concentration (3 nM) of the fluorescently labeled PAR. The binding reactions were performed in triplicate in a buffer comprising 50 mM Tris-HCl pH 7.5, 100 mM KCl, 2 mM MgCl2, 10 mM ß-mercaptoethanol, and 0.1 mg/mL BSA. The reactions were incubated in dark at room temperature for 30 minutes in a Corning 384-well Low Flange Black Flat Bottom Polystyrene NBS Microplate (3575).

After incubation, fluorescence polarization (FP) measurements were performed on a CLARIOstar Plus Microplate Reader from BMG LABTECH equipped with polarizers and Longpass Dichroic Mirror 504 nm. The excitation wavelength was set at 482 nm with 16 nm bandwidth, and emission was monitored at 530 nm with 40 nm bandwidth. The FP values were measured three times, the means of which were analyzed to determine binding affinities. The binding curves were fitted using a nonlinear regression model to determine dissociation constants (K_D_). The FP increase was quantified to indicate the hydrodynamic differences upon proteins binding to ligands. Data analysis and curve fitting were performed using GraphPad Prism.

### Pulsed-field gel electrophoresis and analysis of DSBs

After irradiation and repair incubation, confluent human and bowhead whale skin fibroblasts were harvested, ∼400,000 cells were resuspended in PBS, mixed with an equal volume of 1.4% low gelling temperature agarose and embedded into agarose plugs. Plugs were kept for 1h at +4°C and incubated in lysis solution 1 (0.5M EDTA, 2% sodium sarcosine, 0.5 mg/ml Proteinase K) for 24h at +4°C. Subsequently plugs were placed into lysis solution 2 (1.85M NaCl, 0.15M KCl, 5mM MgCl_2_, 2mM EDTA, 4mM Tris pH 7.5, 0.5% TritonX100) for 40h at +4°C. Plugs were then washed two times for 1h in TE buffer and stored in TE buffer at +4°C. PFGE was carried out with a CHEF DRII system (Bio-Rad) in 0.8% agarose gels. The gels were run at 14°C with linearly increasing pulse times from 50 to 5,000 s for 66 h at a field strength of 1.5 V/cm. Gels were stained with 0.5 mg/ml ethidium bromide for 4h, washed with TBE buffer and imaged. Quantitative analysis was performed with Image Lab software (Bio-Rad). The fraction of DNA entering the gel was quantified. Samples irradiated with various doses and not incubated for repair served as a calibration to determine the percentage of remaining DSBs in the repair samples from the fraction of DNA entering the gel.

### Immunofluorescence

Exponentially growing cells from humans and bowhead whales were cultured on Lab-Tek II Chamber Slides (ThermoFisher Scientific), followed by treatment with bleomycin (BLM) at a final concentration of 5 µg/mL for 1 hour. DNA damage foci were stained with γH2AX and 53BP1 antibodies and quantified at 1 hour and 24 hours. Considering the potential non-specificity of γH2AX and 53BP1 antibodies across species, we used co-localized foci as a more reliable indication of DNA damage.

After BLM treatment, cells were washed twice in PBS, fixed with 2% formaldehyde for 20 minutes at room temperature (RT), washed three times in PBS, and incubated in chilled 70% ethanol for 5 minutes. After three additional washes in PBS, fixed cells were permeabilized with 0.2% Triton X-100 for 15 minutes at RT, washed twice for 15 minutes in PBS, and blocked in 8% BSA diluted in PBS supplemented with 0.1% Tween20 (PBST) for 2 hours at RT. Cells were then incubated with mouse monoclonal anti-γH2AX (Millipore, 05-636, 1:1000) and rabbit polyclonal anti-53BP1 antibodies (Abcam, ab172580, 1:1000) diluted in 1% BSA-PBST at +4°C overnight. After incubation with primary antibodies, cells were washed in PBST three times for 10 minutes and incubated with goat anti-rabbit (Alexa Fluor 488) (Abcam, 1:1500) and goat anti-mouse antibodies (Alexa Fluor 568) (Thermo Fisher Scientific, 1:1000) for 1 hour at room temperature. After four washes for 15 minutes in PBST, slides were mounted in VECTASHIELD Antifade Mounting Medium with DAPI.

For chromatin CIRBP association, cells were pre-incubated with CSK/CSK+R buffer for 3 minutes at RT, washed once in PBS, and subjected to the procedure described above using rabbit monoclonal anti-CIRBP antibodies (Abcam, 1:1000).

Images were captured using the Nikon Confocal system. Confocal images were collected with a step size of 0.5 µm covering the depth of the nuclei. Foci were counted manually under 60x magnification.

### Construction of lentiviral overexpression vectors and lentivirus production

The coding sequences of hCIRBP and bwCIRBP were amplified by PCR using Phusion polymerase (NEB), digested with EcoRI and NotI, and cloned between the EcoRI and NotI sites of the Lego-iC2 plasmid. The sequence was verified by Sanger sequencing. Lentiviral particles were produced in Lenti-X 293T cells (Takara). Approximately 10×10^6^ cells were transfected with a mixture of pVSV-G (1.7 µg), psPAX2 (3.4 µg), and Lego-iC2-bwCIRBP (6.8 µg) using PEI MAX (Polysciences). The day after transfection, the DMEM culture medium (ThermoFisher) was replaced with fresh medium, and lentiviral particles were harvested from the supernatant for the next 3 days.

### Quantification of micronuclei

To analyze binucleated cells containing micronuclei (MN), 10,000-20,000 cells were plated per chamber slide before irradiation or I-SceI transfection. Immediately after treatment, cytochalasin B was added to the cell culture media at a final concentration of 0.5-1 µg/ml, and cells were incubated for an additional 72-120 hours. At the end of the incubation period, cells were washed with PBS, incubated in 75 mM KCl for 10 minutes at RT, fixed with ice-cold methanol for 1.5-3 minutes, air-dried, and stored. Immediately before analysis, cells were stained with 100 µg/ml acridine orange for 2 minutes, washed with PBS, mounted in PBS, and analyzed by fluorescence microscopy. Alternatively, cells were mounted in VECTASHIELD Antifade Mounting Medium with DAPI. At least 100 binucleated cells were analyzed per sample.

### Chromosomal aberration analysis

Metaphase spreads were prepared according to a standard protocol. Briefly, 0.06 µg/ml colchicine (Sigma) was added to the growth medium for 4 hours, and cells were harvested with a 0.25% solution of trypsin/EDTA, treated for 10 minutes with a hypotonic solution (0.075 M KCl/1% sodium citrate) at 37°C, and fixed with three changes of pre-cooled (−20°C) methanol/acetic acid mixture (3:1) at −20°C. Cells were dropped onto pre-cleaned microscope glass slides and air-dried. Metaphase spreads were stained with Giemsa Stain (Sigma) solution in PBS. For each variant, 100 metaphases were analyzed.

### Mismatch repair assay

pGEM5Z(+)-EGFP was a gift from LuZhe Sun (Addgene plasmid #65206; http://n2t.net/addgene:65206; RRID:Addgene_65206). p189 was a gift from LuZhe Sun (Addgene plasmid #65207; http://n2t.net/addgene:65207; RRID:Addgene_65207).

Preparation of the heteroduplex EGFP plasmid was following a published method ^41^ Briefly, pGEM5Z(+)-EGFP plasmid was nicked with Nb.Bpu10I (Thermo Scientific). After phenol/chloroform extraction and ethanol precipitation, the nicked plasmid was digested with Exonuclease III (Thermo Scientific) for 10 minutes at 30°C. p189 was linearized with restriction enzyme BstXI (NEB) and mixed with the purified circular ssDNA at a ratio of 1.0:1.5 to generate a heteroduplex EGFP plasmid containing a G/T mismatch and a nick. The heteroduplex EGFP plasmid with high purity was recovered using a DNA cleanup kit.

Exponentially growing cells were transfected using a 4D-nucleofector (Lonza) with the P1 solution using the DS120 program. In a typical reaction, 2×10^5^ cells were transfected with 50 ng of heteroduplex EGFP plasmid along with 50 ng of DsRed2 to serve as a transfection control. After transfection (48 hours), cells were harvested and analyzed by flow cytometry on a CytoFlex S flow cytometer (Beckman Coulter).

### Host cell reactivation assay

A host cell reactivation assay was employed to assess the repair of UV-induced DNA damage via nucleotide excision repair, following previously described methods^120^.

To evaluate the repair of oxidative DNA damage (base excision repair), a mixture of 20 µg of firefly luciferase (FFL) plasmid and 20-200 µM methylene blue (MB) was prepared, with water added to reach a final volume of 0.4 ml. The DNA-MB mixture was dropped onto a petri dish and placed on ice, with another petri dish containing water positioned on top. Subsequently, the DNA-MB mixture was exposed to visible light for 15 minutes using a 100W lamp positioned at an 11 cm distance. Damaged DNA was then purified, and the host cell reactivation assay was performed as described for UV-induced DNA damage^33^.

### Cyclobutane pyrimidine dimer (CPD) ELISA

Human and bowhead whale skin fibroblasts were cultured until they reached confluency before UVC radiation. Cells were irradiated in PBS at doses of 0, 5, 10, 20, and 30 J/m2 and immediately harvested to construct an induction curve. To assess DNA repair, cells were irradiated at 30 J/m2 and then incubated for 6, 24, and 48 hours before harvesting. Genomic DNA was isolated using the QIAamp Blood Kit (Qiagen). DNA samples were diluted in PBS to a final concentration of 2 µg/ml, denatured at 100°C for 10 minutes, and then incubated in an ice bath for 15 minutes. Next, 100 ng of denatured DNA solution was applied to ELISA plate wells precoated with protamine sulfate (Cosmo Bio) and dried overnight at 37°C. Plates were washed five times with PBS supplemented with 0.05% Tween-20 (PBS-T) and then blocked in 2% FBS in PBS-T for 30 minutes at 37°C. After five washes with PBS-T, plates were incubated with mouse monoclonal anti-CPD antibodies (Clone TDM-2, 1:1000) in PBS for 30 minutes at 37°C. Subsequently, plates were sequentially incubated with goat-anti-mouse biotin IgG (Invitrogen, 1:1000) and streptavidin-HRP (Invitrogen, 1:5000) in PBS for 30 minutes at 37°C each, with five washes with PBS-T before and after each incubation. Plates were then washed with citrate buffer and incubated with a substrate solution (citrate buffer/o-phenylenediamine/hydrogen peroxide) for 30 minutes at 37°C. Finally, the reaction was stopped with 2M H2SO4, and the absorbance was measured at 492 nm using a plate reader.

### CIRBP variant sequence analysis

Identification of rare codons (<10% usage for the corresponding amino acid in human CDS sequences) was performed on CIRBP coding sequences using the Benchling Codon Optimization Tool^121^. Codon adaptation index (CAI) was calculated with human codon frequencies using the E-CAI web server^59^.

### RNA isolation and RNA-seq analysis

RNA from exponentially growing or senescent mouse, cow, human and bowhead whale primary skin fibroblasts was isolated using the RNeasy Plus Mini Kit (Qiagen) according to manufacturer instructions.

Raw reads were demultiplexed using configurebcl2fastq.pl (v1.8.4). Adapter sequences and low-quality base calls (threshold: Phred quality score <20) in the RNA-seq reads were first trimmed using Fastp (0.23.4)^122^. For all species, the clean reads were aligned using Salmon (v1.5.1)^123^ to longest coding sequence (CDS) of each gene extracted from corresponding genome assembly based on human-referenced TOGA annotations. The values of read count and effective gene lengths for each gene were collected and integrated into gene-sample table according to their orthologous relationship. Salmon transcript counts were used to perform differential expression analysis. Only human genes with orthologs in all species were kept for the downstream species. To filter out low expressed genes, only gene with all sample read counts sum >10 were retained.

The filtered count matrix was normalized using median of ratios method^124^ implemented in DESeq2 package^125^. The matrix of effective lengths for each gene in each sample was delivered to the DESeq2 ‘DESeqDataSet’ object to avoid biased comparative quantifications resulting from species-specific transcript length variation. Differential expression analysis was performed using DESeq2 and log transformed fold changes were used for gene set enrichment analysis to assess the differential expression of DNA repair pathways in bowhead whale, cow, and mouse compared to human. Genes of DNA repair pathways were compiled from 3 resources: MsigDB database, GO ontology, and a curated gene list (www.mdanderson.org/documents/Labs/Wood-Laboratory/human-dna-repair-genes.html)^126,127^.

### Nanopore sequencing

72h after transfection, cells were harvested and genomic DNA was isolated with the Wizard Genomic DNA Purification Kit (Promega). DNA concentration was measured on a Nanodrop spectrophotometer and 100 ng of DNA per sample was PCR-amplified with Q5 High-Fidelity 2x Master Mix (NEB). PCR products were prepared for multiplexed Nanopore sequencing using the Native Barcoding Kit 96 V14 SQK-NBD114.96 (Oxford Nanopore Technologies). Following end prep, barcoding, and adapter ligation, samples were cleaned up using AMPure XP Beads and loaded onto a R10.4.1 flow cell on a MinION Mk1C (Oxford Nanopore Technologies) for sequencing. Raw data was basecalled in Super-High accuracy mode with barcode and adapter trimming enabled, demultiplexed, and aligned to the NHEJ reporter construct reference sequence FASTA in Dorado. A custom Python script was used to parse CIGAR strings from the resulting BAM files and quantify indels.

### In vitro Cas9 pulldown

The conserved *PTEN* target amplicon used for assessing DSB repair fidelity was PCR-amplified from untreated control human fibroblasts as described in “Next-generation sequencing of CRISPR repair products.” The *PTEN* target site was cloned into a plasmid using the TOPO TA Cloning Kit (Thermo Fisher Scientific). CRISPR RNP complexes with guide RNAs specific to the target were prepared as follows: 7.2 μL of 200 μM Alt-R™ tracRNA was combined with 7.2 μL of 200 μM Alt-R™ crRNA specific to the *PTEN* genomic target. The tracRNA/crRNA mixture was heated to 95°C and then cooled to room temperature. To the guide duplex were added 25.2 μL of PBS and 20.4 μL of either Alt-R™ S.p. Cas9 Nuclease V3 (Integrated DNA technologies) or Alt-R™ S.p. dCas9 Nuclease V3 (Integrated DNA technologies). RNPs were incubated at room temperature for 20 minutes. Next, 10 μL of Cas9 and dCas9 RNP were combined with 75 μg of *PTEN* TOPO plasmid and incubated at 37°C for 1 hour in EMEM (ATCC). Following incubation, 100 ng of soluble nuclear protein extract from either human or bowhead whale primary fibroblasts, which had been extracted using the NE-PER Nuclear and Cytoplasmic Extraction Kit (Thermo Fisher Scientific), was added to the RNP-plasmid complexes to produce cleavage reactions with Cas9 and control reactions with dCas9 for each species. Nuclear protein extracts were incubated with RNP-plasmid complexes at 37°C for 45 min. To pull down nuclear proteins associated with Cas9/dCas9-plasmid complexes (by the His-tag on Cas9/dCas9), 100 μL Ni-NTA Agarose (Qiagen) was added to each reaction and incubated at 4°C with rotation for 30 min. Ni-NTA Agarose was washed 3x with EMEM + pMaxGFP plasmid to remove proteins with non-specific plasmid binding, and washed 2x with EMEM + 40 mM imidazole + pMaxGFP. To aid elution of bound proteins, the *PTEN* plasmid bound by Cas9/dCas9 was digested by adding 1 μL Benzonase for 5 min. Elution was completed by adding 1x SDS lysis buffer for S-trap with 250 mM imidazole and eluted proteins were analyzed by LC-MS as described in “LC-MS proteomic analysis of fibroblasts.” To distinguish between non-quantifiable and non-detected proteins for figure displays, proteins detected but below the limit of quantification were imputed to an abundance of 10^4^, and proteins not detected were imputed to an abundance of 0. For the figure display, the absolute abundance for each protein was normalized to a value of 1 for the maximum abundance detected for that protein in any of the Cas9 pulldowns.

### Genomic DNA extraction and whole genome sequencing of tumor xenografts

Matching primary cell lines, transformed cell lines, and tumor xenograft samples were prepared as described above. Samples included 1 mouse cell line, 2 human cell lines, and 2 bowhead whale cell lines. 1 fresh cell pellet was prepared for each primary and transformed cell line. For frozen tumor samples, 1 tumor for mouse, 1 tumor for each human cell line (2 tumors total), 4 tumors for whale cell line 14B11SF, and 5 tumors for whale cell line 18B2SF were included in the analysis. Genomic DNA extraction and whole genome sequencing were performed as previously described with minor modifications^49,128^. Briefly, DNA was extracted from samples using the QIamp DNA Mini Kit, per manufacturer’s recommendations. Isolated genomic DNA was quantified with Qubit 2.0 DNA HS Assay (ThermoFisher, Massachusetts, USA) and quality assessed by agarose gel. Library preparation was performed using KAPA Hyper Prep kit (Roche, Basel, Switzerland) per manufacturer’s recommendations. gDNA was sheared to approximately 400bp using Covaris LE220-plus, adapters were ligated, and DNA fragments were amplified with minimal PCR cycles. Library quantity and quality were assessed with Qubit 2.0 DNA HS Assay (ThermoFisher, Massachusetts, USA), Tapestation High Sensitivity D1000 Assay (Agilent Technologies, California, USA), and QuantStudio ® 5 System (Applied Biosystems, California, USA). Illumina® 8-nt dual-indices were used. Equimolar pooling of libraries was performed based on QC values and sequenced on an Illumina® NovaSeq X Plus (Illumina, California, USA) with a read length configuration of 150 PE for 60M PE reads (30M in each direction) per sample.

### Bioinformatic analysis of tumor xenograft whole genome sequencing

The bioinformatic processing pipeline of raw whole-genome (WGS) high throughput sequencing data was adapted for human, murine and bowhead whale data ^49^.

Sequencing FASTA files were applied to FastQC^129^ for quality control, adapters were trimmed by Trimmomatic^130^, and the genomic fragments were aligned to the human, mouse, and whale genome reference (hg19, mm10 and the published bowhead whale genome assembly^17^) using Burrows-Wheeler Aligner (BWA)^131^, then sorted and indexed by Samtools^132^. Somatic mutations were detected from tumor samples using MuTect v2^133^ to call somatic single-nucleotide variants (SNVs) and small indels (<10bp). Tumor samples from WGS were compared to their respective matched healthy tissue. All mutations were also filtered for depth (tumor sample coverage > 30x, normal sample coverage > 30x) and variant allele frequency (VAF ≥ 0.1). Structural variations were called by Manta applying default settings and structural variant length > 6000bp were used for downstream analysis ^134^.

### Alkaline Comet Assay

For the alkaline comet assay, we adapted the Alkaline CometAssay protocol provided by TREVIGEN based on a published in-gel comet assay method^135^ to increase the number of cell lines and time points assessed and minimize assay variation introduced during sample harvesting and processing. Slides were pre-coated with a base layer 50µl of 1% SeaKem LE Agarose (Lonza) to enhance adhesion. We cultured cells to near 100% confluency and then resuspended them in CometAssay LMAgarose (R&D Systems). We applied 500 cells suspended in 100 μL LMAgarose onto each slide. The slides were then placed in the dark at 4°C for 10 minutes to allow the agarose to solidify. After that, slides with live cells were incubated in tissue culture incubators in fibroblast culture medium containing 700 µM freshly diluted H_2_O_2_ for 30 minutes, followed by washing with PBS and incubation for various recovery periods (ranging from 0 minutes to 12 hours) in culture medium. Slides were collected at each time point, washed with PBS, and immersed in CometAssay Lysis Solution (R&D Systems). Before electrophoresis, slides were placed in alkaline unwinding solution prepared according to TREVIGEN’s protocol for 10 minutes. After electrophoresis at 22V for 30 minutes, the slides were placed in a DNA precipitation buffer following the TREVIGEN protocol for 10 minutes and subsequently washed three times with distilled water. The slides were then immersed in 70% ethanol for 10 minutes and allowed to air dry in the dark. Before imaging, each sample was stained with 50 µl of 1x SYBR Gold (Thermo Fisher Scientific) for 5 minutes before being washed three times with distilled water. Comet images were acquired through fluorescent microscopy. For scoring, we used profile analysis in OpenComet^136^ within ImageJ. Outliers automatically flagged by OpenComet were excluded from analysis and remaining incorrectly demarcated comets were further systematically filtered out according to two criteria: a comet area greater than 5000 or head area greater than 500.

### Tissue processing

Tissues obtained from wild-caught animals were assumed to be of younger/middle age since predation normally precedes aging in the wild. Postmortem interval was minimized, and, in all cases, samples were kept on ice and frozen in less than 24h. At the earliest opportunity after dissection, tissues from representative animals from each species were flash frozen in liquid nitrogen and stored at −80°C. Tissues were pulverized to a fine powder within a Biosafety cabinet under liquid nitrogen using a stainless-steel pulverizer Cell Crusher (Fisher Scientific) chilled in liquid nitrogen and delivered to storage tubes with a scoop that had also been pre-chilled in liquid nitrogen and kept on dry ice. Similarly, when sampled for various “omics” processing, pulverized tissues were removed with a stainless-steel spatula that was pre-chilled in liquid nitrogen. Samples were never thawed after initial freezing until extractions were performed.

### Cross-species tissue proteomics

We employed a “shotgun” style untargeted data-dependent acquisition (DDA) label-free quantitative (LFQ) approach. Approximately 5 mg of tissue was mixed with 250µl of 50mM TEAB pH7.6; 5% SDS, mixed by pipetting, and briefly vortexed. Samples were sonicated in a chilled cup-horn Q800R3 Sonicator System (Qsonica; Newtown, CT) for a total of 15min at 30% output and duration of 30 x 30 sec pulses (with 30 sec in between pulses) at 6°C using a chilled circulating water bath. When nuclear proteomes were analyzed, nuclei were first isolated using a hypotonic lysis approach as in the preparation of histones^137^. Isolated nuclei were lysed and processed as indicated above with SDS and sonication and then handled similarly for the rest of the prep. Samples were heated to 90°C for 2min and allowed to cool to room temperature (RT). Next, samples were centrifuged at 14,000xg for 10min to pellet insoluble debris and the supernatants were transferred to clean tubes. Total protein was quantified by the BCA assay and 100µg was reduced with 5 mM dithiothreitol (DTT) for 30min at 60°C. Samples were cooled to RT and then alkylated with 10mM iodoacetamide (from a freshly prepared stock) for 30 min at RT in the dark. Samples were processed using the standard S-trap mini column method (Protifi; Farmingdale, NY). Samples were digested with 4μg trypsin overnight at 37°C. Elution fractions were pooled and dried using a Speedvac (Labconco). Peptides were resuspended in 100μl MS-grade water (resistance ≥18MΩ) and quantified using the Pierce Quantitative Fluorometric Peptide Assay (Thermo). Common internal Retention Time standards (CiRT) peptide mix was added (50 fmol mix/2 µg tryptic peptides) and 2 µg (in 4 µL) of tryptic peptides were injected/analyzed by mass spectrometry (MS) on a Orbitrap Tribrid Fusion Lumos instrument (Thermo) equipped with an EASY-Spray HPLC Column (500mm x 75um 2um 100A P/N ES803A, Nano-Trap Pep Map C18 100A; Thermo). Buffer A was 0.1% formic acid and buffer B was 100% acetonitrile (ACN) with 0.1% FA. Flow rate was 300nl/min and runs were 150 min: 0-120 min, 5% B to 35% B; then from 120-120.5 min, 35-80% B; followed by a 9-minute 80% B wash until 130min. From 130-130.5min B was decreased to 5% and the column was re-equilibrated for the remaining 20-min at 5% B. the instrument was run in data dependent analysis (DDA) mode. MS2 fragmentation was with HCD (30% energy fixed) and dynamic exclusion was operative after a single time and lasted for 30sec. Additional instrument parameters may be found in the Thermo RAW files.

### Computational proteomics analysis

Raw files were analyzed directly with the MSFragger/Philosopher pipeline^138,139^ and included Peptide and Protein Prophet modules^140^ for additional quality control.

Quantitation at the level of MS1 was performed with the “LFQ-MBR; label-free quant match-between-runs” workflow using default parameters. This allows for alignment of chromatographic peaks between separate runs. Methionine oxidation and N-terminal acetylation were set as variable modifications. MaxLFQ with a minimum of two ions was implemented and normalization of intensity across runs was selected^141^.

### LC-MS proteomic analysis of fibroblasts

2 15-cm dishes of growing primary fibroblasts from 2 cell lines for each species were harvested for protein. Cells were washed with PBS and pellets were snap frozen and stored in liquid nitrogen until processing. Cells were solubilized with 5% SDS; 50 mM TEAB pH 7 and sonicated at 8°C with 10x 45s pulses using 30% power with 15 s rest between each pulse with a cup horn Q800R3 Sonicator System (Qsonica; Newtown CT). Soluble proteins were reduced with 10 mM DTT for 30 min at 55°C, followed by alkylation with 15 mM iodoacetamide at 25°C in the dark for 30 min. S-trap micro columns (Protifi; Farmingdale, NY) were employed after this step for overnight tryptic digestion and peptide isolation according to manufacturer instructions. All solvents were MS-grade. Resulting tryptic peptides were resuspended in MS-grade water and were quantified using a Pierce™ Quantitative Fluorometric Peptide Assay (Thermo Fisher cat #23290). Prior to MS, peptides were mixed with a common internal retention time standards115 (CiRT) peptide mix (50 fmol CiRT/2µg total tryptic peptides) and acetonitrile (ACN) and formic acid were added to concentrations of 5% and 0.2% respectively. The final concentration of the peptide mix was 0.5µg/µl. 2 µg (4µl) of each were resolved by nano-electrospray ionization on an Orbitrap Fusion Lumos MS instrument (Thermo) in positive ion mode. A 30 cm home-made column packed with 1.8 μm C-18 beads was employed to resolve the peptides. Solvent A was 0.1% formic acid and solvent B was 80% acetonitrile with 0.1% formic acid and flow rate was 300 nl/min. The length of the run was 3 h with a 155 min gradient from 10-38% B. HCD (30% collision energy) was used for MS2 fragmentation and dynamic exclusion was operative after a single time and lasted for 30s. Other details of the run parameters may be found in the embedded run report of the RAW data file uploaded to the ProteomeXchange database. Peptide assignments and quantitation were done using the label-free quant match between runs (LFQ-MBR) workflow of MSFragger^138–140^. MaxLFQ with a minimum of two ions was implemented and normalization was selected. Additional details are available in MSFragger log files. Searches were performed within the Philosopher/Fragpipe pipeline that incorporates PeptideProphet and ProteinProphet filtering steps to increase the likelihood of correct assignments^140^. The databases used for searches were predicted proteins from the published bowhead genome^17^ as well as our custom proteome derived from our de novo sequenced and Trinity^89,142^–assembled pool of transcriptomes from whale tissues. Human (UP000005640), mouse (UP000000589), and bovine (UP000009136) databases were from the latest build available from Uniprot^143^. For the searches, databases also included a reverse complement form of all peptides as well as common contaminants to serve as decoys for false discovery rate (FDR) calculation by the target/decoy approach (decoy present at 50%). Final FDR was below 1%. To distinguish between non-quantifiable and non-detected proteins in figure displays, proteins detected but below the limit of quantification were imputed to an abundance of 10^4^, and proteins not detected were imputed to an abundance of 0.

### LC-MS identification of proteins upregulated after DNA damage

3 primary fibroblast lines each for human and bowhead whale were prepared under 3 conditions: untreated control, H_2_O_2_-treated, and UV treated. For H_2_O_2_ treatment, culture medium was replaced with medium containing 400 μM H_2_O_2_ that had been diluted into the medium immediately prior to use. For UV treatment, culture medium was aspirated and replaced with a thin layer of PBS. Cells were exposed to 6 J/m^2^ UVC in a UV Crosslinker (Fisher Scientific) with the culture dish lid removed. 4h after treatments, cells were harvested by washing with PBS and lysed directly in-dish by addition of 2x SDS lysis buffer for S-trap. Cells were subsequently processed for LC-MS as described in “LC-MS proteomic analysis of fibroblasts.” Data were acquired on an Orbitrap Astral Mass Spectrometer (Thermo) equipped with an EASY-Spray HPLC Column (500mm x 75μm 2μm 100A P/N ES803A, Nano-Trap Pep Map C18 100A; Thermo). Samples were run in DIA mode. Computational-Raw data files were converted to mzML files using ProteoWizard with peak picking set to 1-n, and demultiplexing selected^144,145^. The mzML files were searched using DIANN^146^ with along with FASTA file databases described above for tissue extracts using library-free search/library generation with deep learning-based spectra retention time (RT) and IMs prediction criteria selected. Mass accuracy was set to 20 and MS1 accuracy set to 5.0, and oxidized methionine (Ox(M)) was also selected. Nearest human homologs for each species’ protein (determined by BLAST as previously described^89^) were added to the protein group matrix (DIANN output). As before, the human annotation was used to facilitate cross-species comparisons. Missing values for protein groups were imputed using deterministic minimum imputation^147,148^.

### Stimulated emission depletion (STED) microscopy

Immortalized primary fibroblast cultures were plated on 10mm diameter glass coverslips embedded in 35mm plates (Mattek, P35G-1.5-10-C) at a concentration of 10,000 cells per plate. 48 hours after plating, plates were washed once with 1x PBS and fixed in 3.7% formaldehyde for 15min on a shaking platform. Plates were washed 3x 10min in PBS and permeabilized in 0.5% Triton X-100 for 15 minutes on a shaking platform.

Plates were washed again 3x 10min in PBS before blocking for 1h at RT with 5% BSA in PBS. After blocking, primary antibody was added at a concentration of 1:50 diluted in 100µL blocking buffer and incubated overnight at 4°C. Plates were washed in wash buffer ( 0.1% Triton X-100 in PBS) 4x 10min before adding Alexa Fluor 594-labelled secondary antibody (Abcam, ab150080) at a concentration of 1:50 diluted in 100µL blocking buffer and incubating for 1h at RT. Plates were washed 4x 10min with wash buffer and incubated with 100 µL of 1 µg/mL DAPI in PBS for 5min. Plates were washed once with PBS for 5min before adding mountant (Invitrogen, P10144) and placing a cover slip. Gated Stimulated Emission Depletion microscopy was performed in the URMC Center for Advanced Light Microscopy and Nanoscopy (RRID:SCR_023177) on an Abberior) Göttingen, Germany) 3D STED instrument (S10 OD023440) equipped with an Olympus UPlanSApo 100x/1.4NA objective. Images were acquired with a pixel size of 20nm and 775nm depletion laser at 30% power with 15% directed toward 3D resolution. Single-channel images were pseudocolored red (Alexa-Fluor 594) and blue (DAPI), merged, and scale bars added using ImageJ software.

### Doxycycline-inducible I-SceI NHEJ reporter

The plasmid was assembled from several parts. The backbone was amplified from a pN1 plasmid without f1 bacteriophage origin of replication and modified by the addition of short insulator sequences^149^ (E2, A2 and A4) purchased from Integrated DNA Technologies (IDT). The GFP reporter gene with I-SceI endonuclease sites was amplified from the reporter described above and fused via the P2A self-cleaving peptide with TetOn transactivator, amplified from Lenti-X™ Tet-One™ Inducible Expression System Puro (Takara, 631847). A bi-directional promoter sequence featuring hPGK and TRE3GS was amplified from the same plasmid and cloned upstream of the GFP reporter, in the orientation for TetOn-P2A-reporter to be driven by the constitutive hPGK promoter. Downstream of the Tre3GS promoter was closed codon-optimized sequence for intron-encoded endonuclease I (I-SceI) with SV40 nuclear localization sequence (NLS) at the N-terminus and nucleoplasmin NLS at the C-terminus fused to the enhanced blue fluorescent protein (eBFP2) via P2A. The fusion was purchased from IDT. EBFP2 sequence was derived from eBFP2-N2 plasmid (Addgene #54595).

Cloning was done with In-fusion Snap assembly kit (Takara 638947), NEBuilder HiFi DNA Assembly (NEB, E5520) and T4 DNA ligase (NEB, M0202). The efficiency of non-homologous end-joining (NHEJ) double-strand break (DSB) repair was analyzed in immortalized normal human dermal fibroblasts (NHDF2T). The expression cassette containing a GFP reporter gene under hPGK promoter and an I-SceI endonuclease under doxycycline-inducible Tre3GS promoter was inserted into the genome by random integration method. The positive clones were selected by G418 for 10 days and the clones were pooled together. The GFP reporter had a short adeno-exon flanked by two I-SceI recognition sites (in inverted orientation) surrounded by the rat Pem1 intron.

Upon stimulation with doxycycline (100 ng/ml), I-SceI produced two non-ligatable double strand breaks, resulting in excision of the adeno-exon and reconstitution of the functional GFP.

### EMSA

Recombinant human CIRP protein (Abcam AB106903) was incubated in the indicated amounts with the indicated nucleic acid substrates in 20 μL EMEM (ATCC) at 37°C for 1 hour. Subsequently, reactions were mixed with 4 μL sucrose loading dye (2M sucrose + 0.2% Orange G) and loaded into agarose gels immersed in 0.5x TAE buffer followed by electrophoresis at 30V. Following electrophoresis, gels were stained in 1x SYBR Gold (Thermo Fisher Scientific) and imaged. Extraction of genomic DNA from human primary fibroblasts was with the Monarch HMW DNA Extraction Kit for Cells & Blood (NEB #T3050L). To produce the damaged DNA samples and induce PAR formation, cells were treated with H_2_O_2_ and UV prior to genomic DNA extraction. For H_2_O_2_ treatment, culture medium was replaced with medium containing 400 μM H_2_O_2_ that had been diluted into the medium immediately prior to use. For UV treatment, culture medium was aspirated and replaced with a thin layer of PBS. Cells were exposed to 6 J/m^2^ UVC in a UV Crosslinker (Fisher Scientific) with the culture dish lid removed. During genomic DNA extraction from damaged chromatin, Proteinase K was added per manufacturer instructions, but RNase A was omitted and Protector Rnase inhibitor (Sigma-Aldrich) was added to the extraction buffers and eluate. Nucleic acids used in reactions were sonicated to uniform size in a QSONICA Sonicator.

### Statistical analyses

Statistical comparisons were performed as indicated in the figure legends. Unless otherwise specified in the text or legend, *n* refers to separate biological replicate cell lines, isolated from different individuals for a given species. Exceptions include specific genetically modified cell lines or clones, e.g. tumor suppressor knockout lines and Ku-deficient MEFs. In such cases, *n* refers to technical replicates and indicates the number of times the experiment was repeated with the specified cell line. Details for comparisons done by ANOVA are included in Supplementary Information.

## Data Availability

Proteomics data are accessible through ProteomeXchange [URL to be added]. DNA and RNA sequencing data are accessible through NCBI Sequence Read Archive (SRA) [URL to be added].

---

**Extended Data Figure 1.**
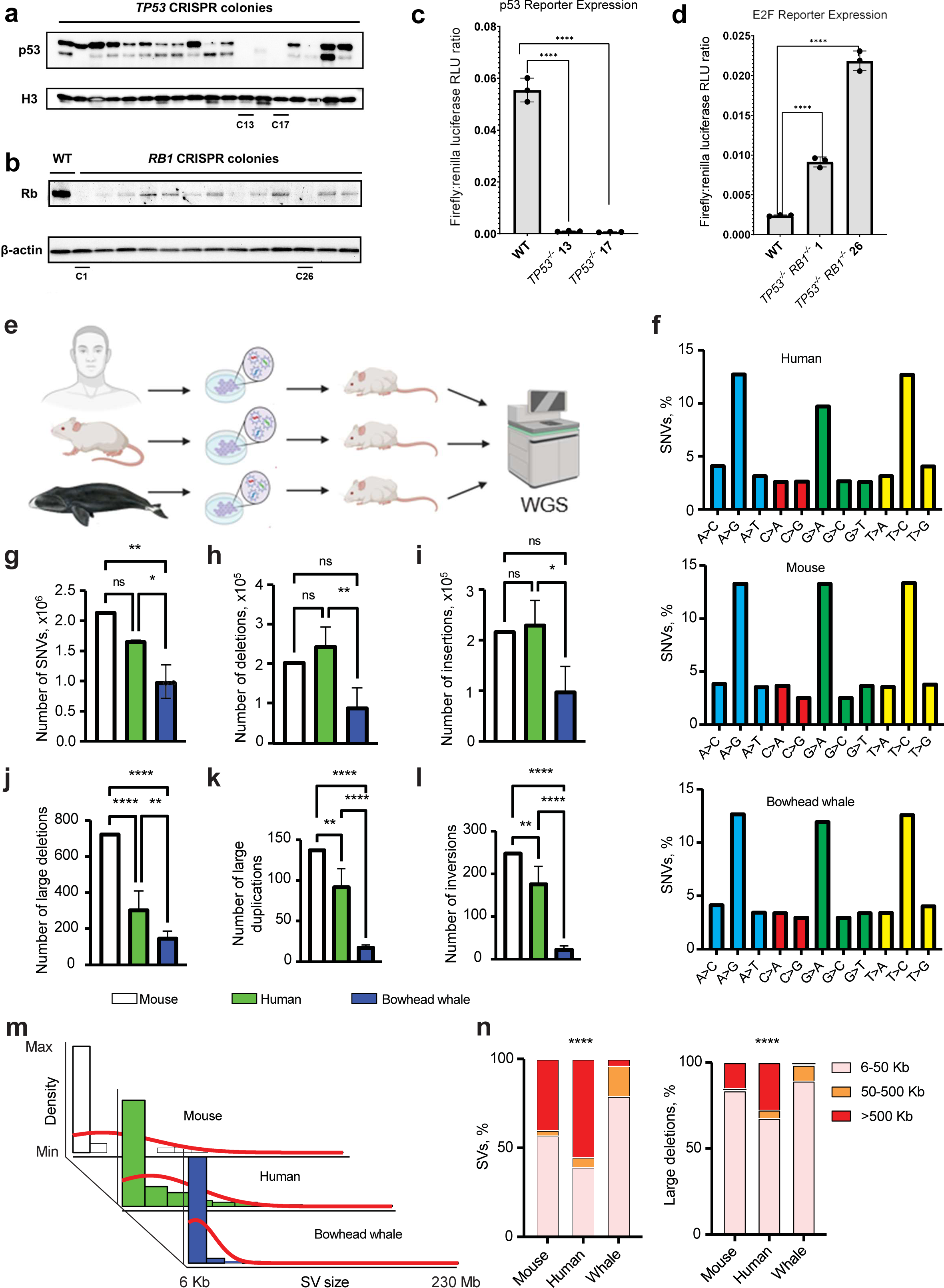
Mutation rates in bowhead whale cells during tumor progression. **a,** Western blot for p53 protein in clonally isolated fibroblast colonies following CRISPR targeting of *TP53*. Underlined lanes indicate colonies selected for further validation and experiments. **b,** Western blot for Rb protein in clonally isolated fibroblast colonies following CRISPR targeting of *RB1* on an existing p53 knockout background. **c,** Ratio of firefly:renilla luciferase luminescence in fibroblasts transfected with firefly luciferase reporter of p53 transcriptional activity and renilla luciferase control. Cells were treated with etoposide to induce p53 activity. **d,** Ratio of firefly:renilla luciferase luminescence in fibroblasts transfected with firefly luciferase reporter of E2F transcriptional activity and renilla luciferase control. Transfected cells were serum starved for 24h and returned to complete medium for 24h before luminescence measurement. Higher E2F activity results from reduced Rb activity. Error bars represent SD. p<0.001 (two-tailed t test), n=3. **e,** Schematic showing experimental design and samples processed for WGS (whale N = 9 tumors; human N = 2 tumors; mouse N = 1 tumor). **f,** Bar plot displaying percentages of SNV types across species with similarities of mutational processes across species. **g-I,** Bar plot showing quantifications of numbers of SNVs and small indels (size 1-10bp) across species. **j-l**, Bar plot showing quantification of number of large SVs (size > 6000bp) across species. **m,** Histograms and trend curves showing distribution of SVs size across species. **n,** Bar plot showing distribution of small, medium and large (6-50Kb, 50-500Kb, >500Kb respectively) SVs and deletions across species. Error bars represent SD. P values are a result of ordinary One-Way Anova with Tukey’s multiple comparison test (**g-l**) and chi-square test (**n**). * p < 0.05; ** p < 0.01; *** p < 0.0001; ns = not significant.

**Extended Data Figure 2.**
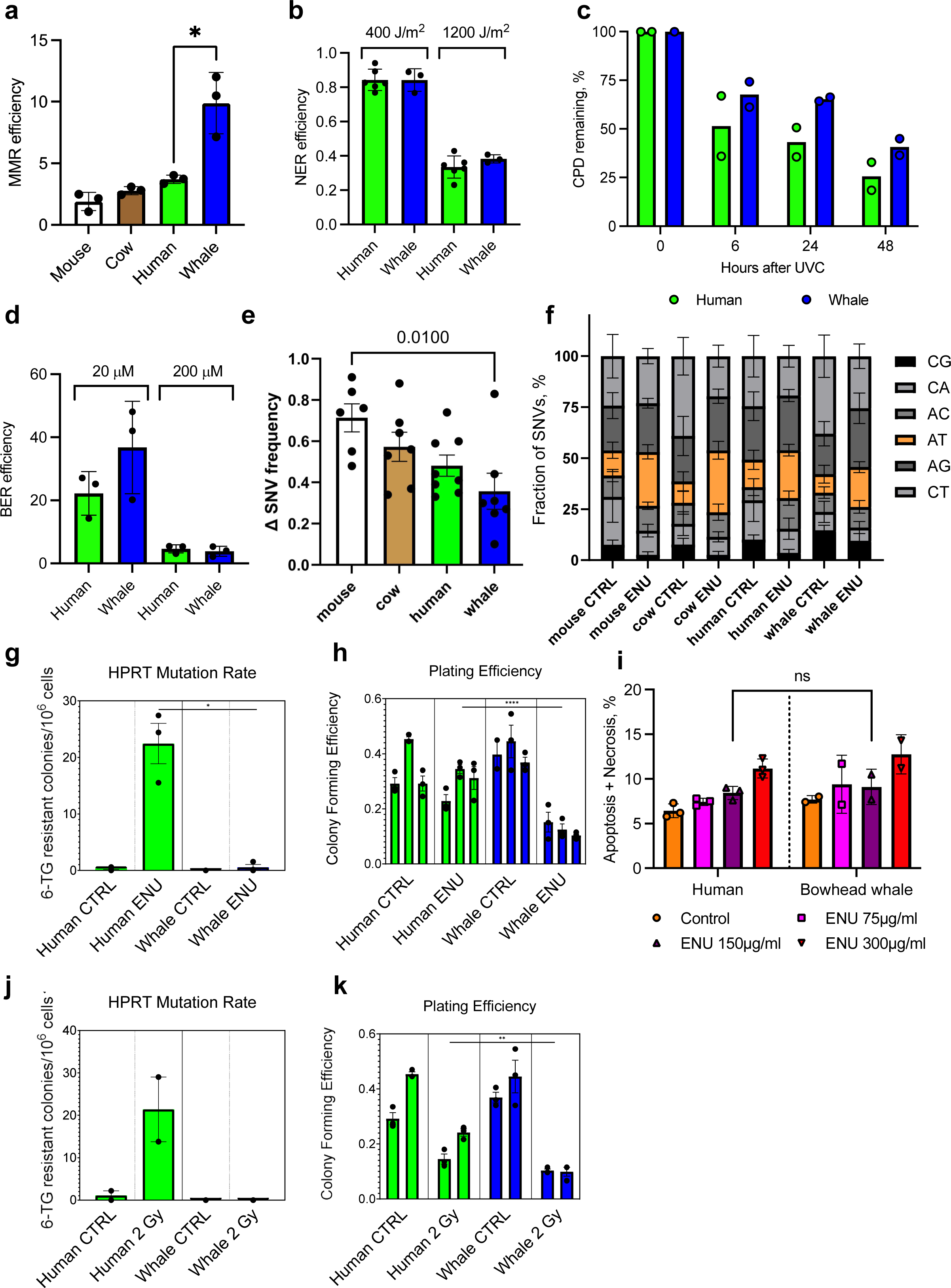
Mismatch repair, excision repair, and mutagenesis in bowhead whale cells. **a,** MMR reactivation of a heteroduplex eGFP plasmid containing a G/T mismatch. Growing fibroblasts were transfected with the heteroduplex plasmid and a DsRed plasmid as a transfection control. The repair efficiency was calculated as the ratio of GFP+/DsRed+ cells. Each dot represents cell line isolated from different individual (n=3). **b**, NER efficiency was measured by host cell reactivation assay where a plasmid containing luciferase reporter is UV-irradiated *in vitro* to induce DNA damage, transfected into cells, and reactivation of the reporter is measured (n=3 for each cell line). **c**, Kinetics of cyclobutane pyrimidine dimer repair after 30 J/m^2^ UVC. Confluent human and whale skin fibroblasts were subjected to UVC, harvested at different time-points, genomic DNA was isolated and analyzed for cyclobutene dimers as described in Methods (n=2 for each cell line). **d**, BER efficiency was measured by host cell reactivation where luciferase reporter plasmid is treated with methylene blue and light to induce oxidative DNA damage, transfected into cells, and luciferase activity measured as described in Methods. **e**, ENU-induced mutational load by SMM-seq in fibroblasts of the indicated species. Delta SNV frequency was calculated for each cell line (n=6-8 fibroblasts/species; Kruskal-Wallis test). **f**, Analysis of mutational spectra showing a pattern typical for ENU. An increase in A>T transversions (orange bars) can be found in ENU-treated mammalian cells. **g**, HPRT mutagenesis assay in ENU-treated cells, adjusted by plating efficiency measured for each cell line (n=3 cell lines/species) **h**, Colony forming efficiency for HPRT mutagenesis assay. Error bars represent mean ± SD. * p<0.05, ** p<0.01, *** p<0.001, **** p<0.0001 ns=not significant (heteroscedastic two-tailed t test). **i,** Apoptosis/necrosis of human and bowhead whale fibroblasts in response to ENU. Cells at growing stage were treated for 3h with ENU at indicated dosages. After treatment cells were washed in PBS and incubated for 3 days. For measuring Apoptosis/Necrosis cells were stained with AnnexinV/PI and analyzed by flow cytometry. **j**, HPRT mutagenesis assay in cells treated with 2 Gy *γ*-irradiation, adjusted by plating efficiency measured for each cell line (n=2 cell lines/species) **k**, Colony forming efficiency for HPRT mutagenesis assay.

**Extended Data Figure 3.**
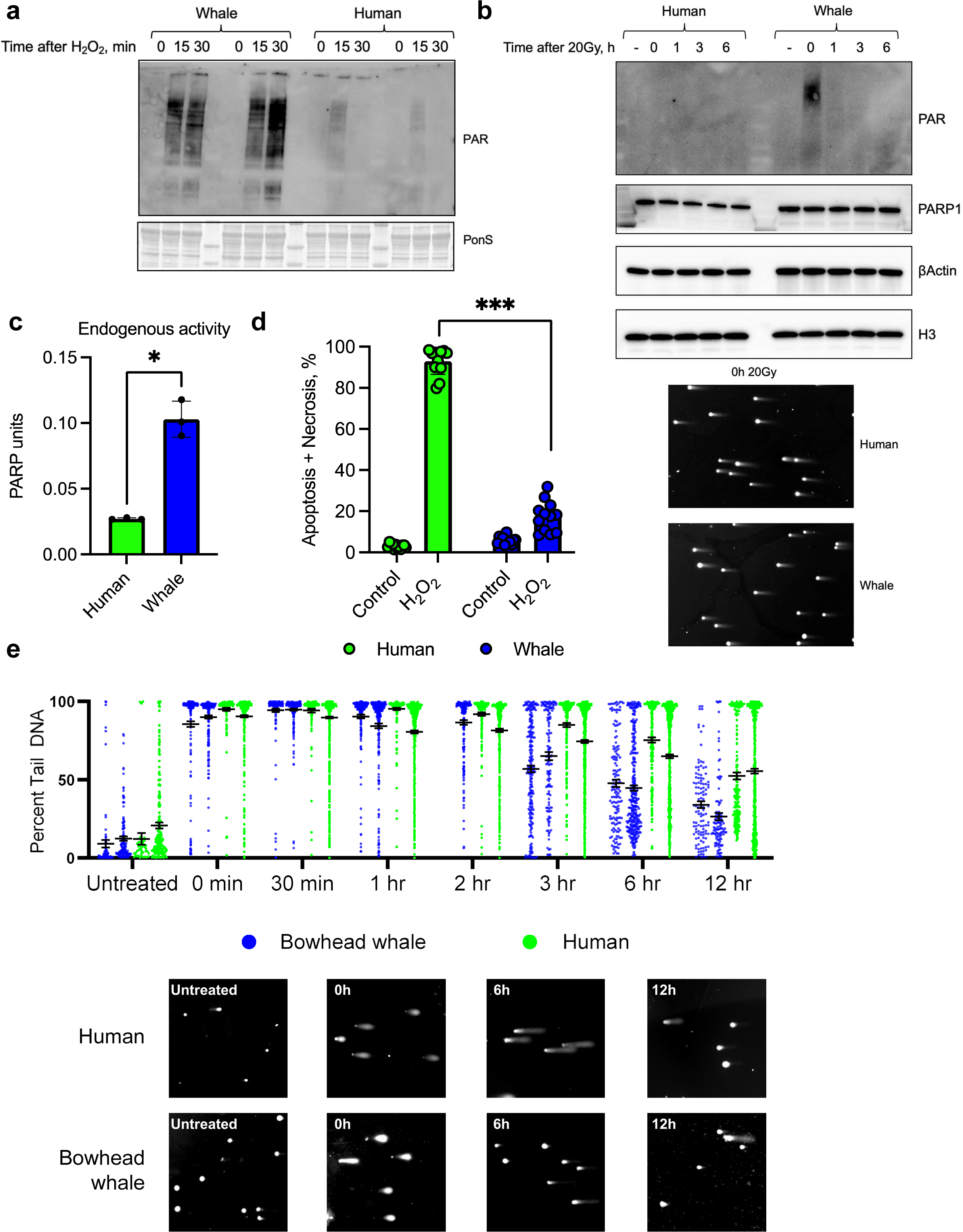
Poly-ADP-ribosylation and DNA repair of oxidative damage in bowhead whale cells. **a**, Bowhead whale cells show greater poly-ADP-ribosylation in response to hydrogen peroxide treatment. **b,** Bowhead whale cells show greater poly-ADP-ribosylation after γ-irradiation. Cells were harvested immediately or at indicated time-points after radiation for Western blot analysis (top panel). Representative images of comet tails under neutral conditions (bottom panel). Cells were processed immediately after radiation. **c**, Nuclear extracts of bowhead whale fibroblasts exhibit higher endogenous PARP activity (n=3). Error bars represent mean ± SD. * p<0.05 (Welch’s t-test). Whale=bowhead whale. **d,** Apoptosis/Necrosis of human and bowhead whale fibroblasts in response to hydrogen peroxide at concentration 700µM. Two days after hydrogen peroxide, cells were harvested and subjected to an Annexin V apoptosis assay using flow cytometry. Error bars represent mean ± SD. *** p<0.001. Welch’s t-test was used to quantify the significance (n=12). **e,** Percent tail DNA by alkaline comet assay at various time points after 700 μM H_2_O_2_ treatment in 2 cell lines each of human and bowhead whale fibroblasts. Points represent individual cells. Representative comet images shown below. Bars indicate mean +-SEM.

**Extended Data Figure 4.**
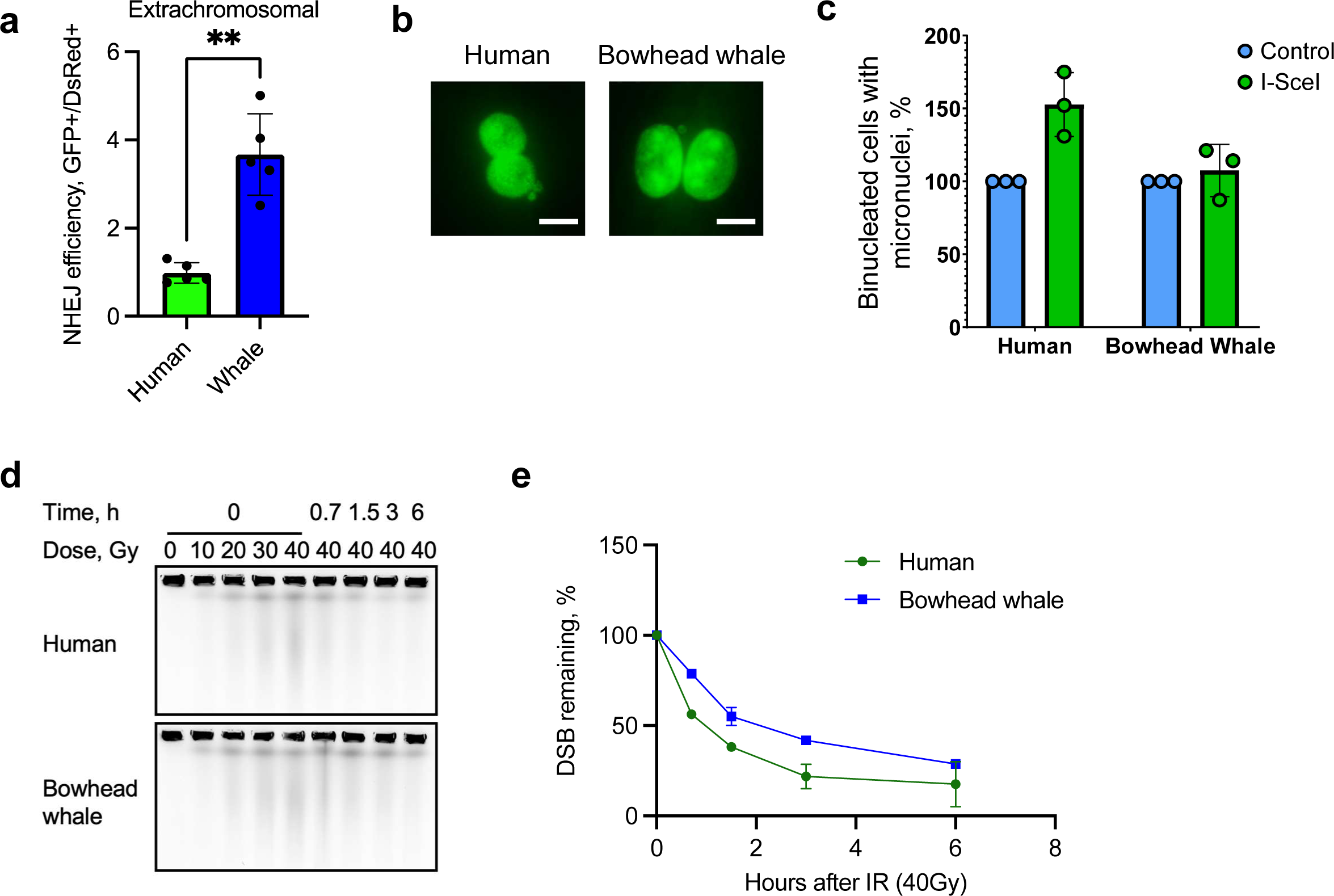
DSB repair efficiency in bowhead whale cells. **a,** NHEJ efficiency in extrachromosomal assay. NHEJ reporter construct was pre-digested with I-SceI, purified and co-transfected with DsRed into human and bowhead skin fibroblasts. Three days after transfection cells were harvested and subjected to flow cytometry to calculate NHEJ efficiency (n=3). Error bars represent mean ± SD. *** p<0.001 (two-tailed t-test) **b,** Representative images of human and bowhead whale binucleated cells containing micronuclei after 2 Gy of *γ*-irradiation. Scale bar indicates 20 µm. **c,** Frequency of micronuclei after DSB induction with I-SceI in primary fibroblasts carrying a chromosomally integrated NHEJ reporter cassette. Each cell line was transiently transfected with a BFP-expressing control plasmid or an I-SceI expression plasmid and micronuclei were quantified after 5d in media containing cytochalasin B to prevent cytokinesis. Micronucleus frequencies for each cell line are shown normalized to BFP control (paired t-test, n=3 cell lines/species). **d,** Pulse-field gel stained with ethidium bromide, showing chromosomal DNA fragmentation in human and bowhead confluent skin fibroblasts immediately after different doses of *γ*-irradiation 0.7, 1.5, 3 and 6h after 40 Gy of *γ*-irradiation. **e,** Kinetics of DSB repair measured by PFGE in confluent human and bowhead fibroblasts after 40 Gy of *γ*-irradiation. n=2 for each species.

**Extended Data Figure 5.**
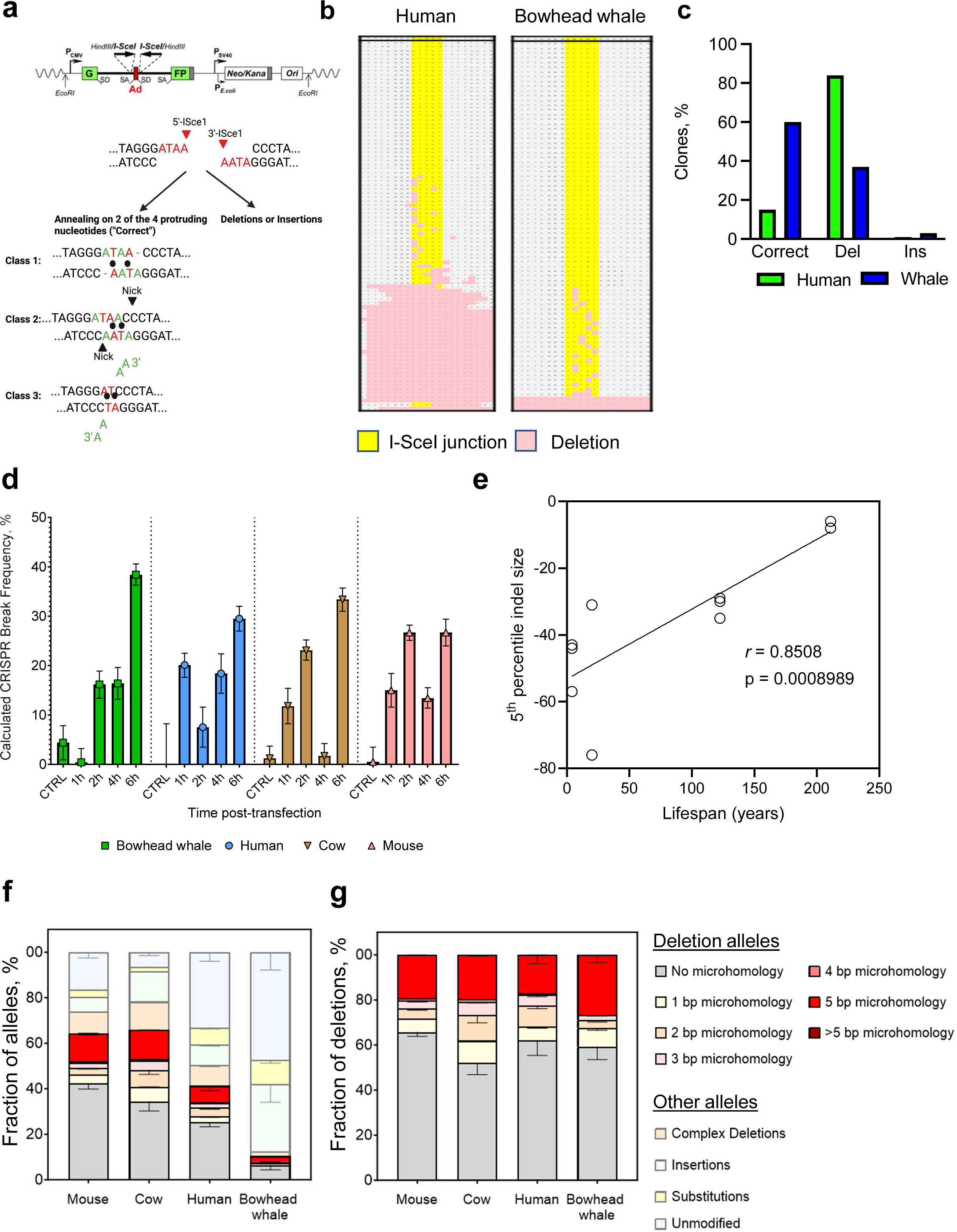
Sequencing of DNA DSB repair products in bowhead whale cells. **a,** Possible repair outcomes after induction of DSBs with incompatible ends by I-SceI in NHEJ reporter construct. **b,** Allele plot of Sanger sequencing products resulting from repair of integrated NHEJ reporter cassette after I-SceI cleavage. **c,** NHEJ fidelity in extrachromosomal assay. NHEJ reporter construct was pre-digested with I-SceI, purified and co-transfected with DsRed into human and bowhead skin fibroblasts. Three days after transfection genomic DNA was isolated, subjected to PCR, cloned and analyzed by Sanger sequencing. At least 100 clones were analyzed for each species. Correct – annealing on 2 of the 4 protruding nucleotides **d,** Time course of CRISPR cleavage measured by digital droplet PCR (ddPCR). PTEN copy number at varying time points after CRISPR RNP transfection was measured with ddPCR using primers flanking the predicted cleavage site and normalized within each sample to a single-copy genomic ultraconserved element as described in Methods. Error bars show confidence intervals of Poisson distribution calculated in QuantaSoft. **e,** Pearson correlation between 5^th^ percentile indel size and species lifespan (r=0.8508, 95% CI = 0.5125 to 0.9605, p=0.0009, n=11). **f,** Absolute frequencies of alleles by base pairs of microhomology across species in CRISPR-targeted PTEN repair products. **g,** Relative proportions of deletion alleles by base pairs of microhomology across species in CRISPR-targeted PTEN repair products.

**Extended Data Figure 6.**
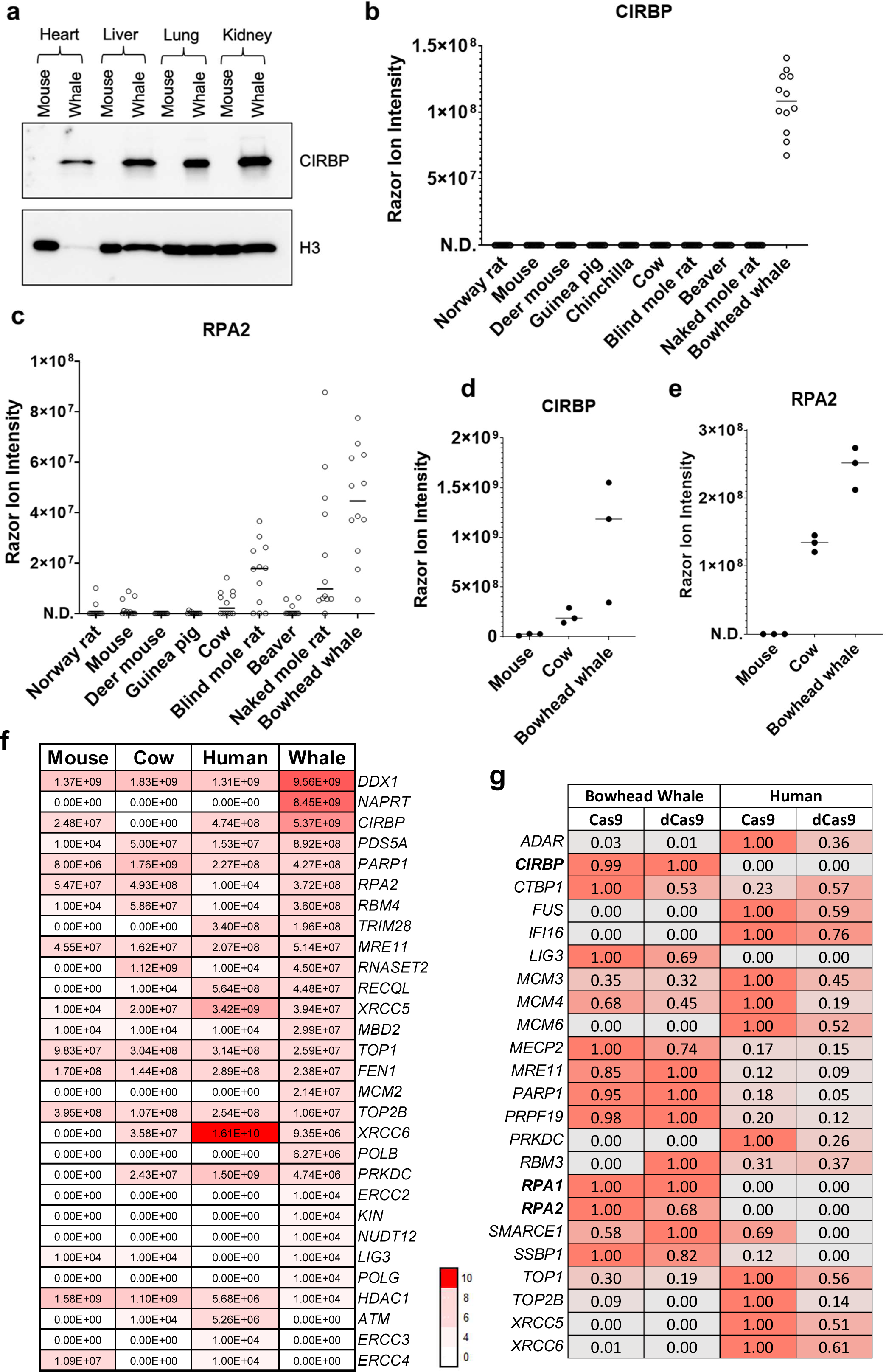
Proteomic quantification of DNA repair proteins. **a,** Western blot for CIRBP on bowhead whale and mouse organs **b,** Abundance of CIRBP protein by LC-MS in liver tissue of mammal species (n=12 per species; 3 biological x 4 technical replicates; N.D.=not detected) **c,** Abundance of RPA2 protein by LC-MS in liver tissue of mammal species (n=12 per species; 3 biological x 4 technical replicates; N.D.=not detected) **d,** Abundance of CIRBP protein by LC-MS in nuclear extracts of liver tissue of mammal species (n=3 biological replicates per species) **e,** Abundance of RPA2 protein by LC-MS in nuclear extracts of liver tissue of mammal species (n=3 biological replicates per species; N.D.=not detected) **f,** Heatmap of LC-MS protein abundance for primary fibroblasts of the indicated species and proteins. Color intensity scale corresponds to log_10_ ion intensity. **g,** Per-protein normalized abundance by LC-MS of proteins identified in pulldowns of His-tagged Cas9/dCas9 bound to a plasmid containing the genomic *PTEN* target sequence after incubation in extracts of soluble nuclear proteins from human and bowhead whale.

**Extended Data Figure 7.**
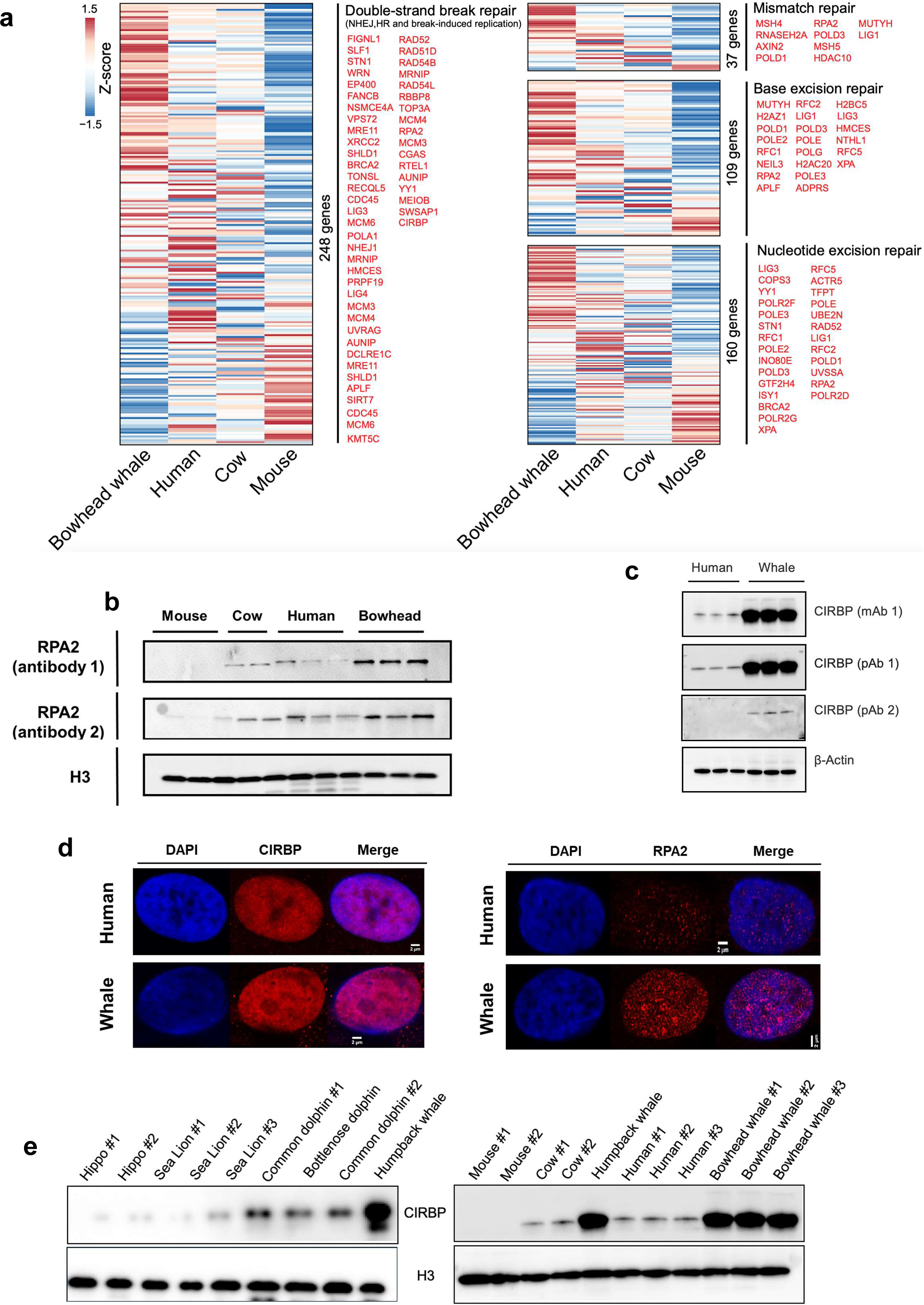
Transcriptome, Western blot, and STED quantification of DNA repair proteins. **a,** Relative expression level of genes in 6 DNA repair pathways among species. Z-scores are scaled by row. Genes in each pathway are ordered decreasingly based on the expression level in bowhead whale. Genes with higher expression in bowhead whale compared to all 3 other species are highlighted in red text to the right of the heatmap. Genes of each gene set were compiled from 3 resources: MsigDB database, GO ontology, and a curated gene list (www.mdanderson.org/documents/Labs/Wood-Laboratory/human-dna-repair-genes.html) **b,** Western blot abundance of RPA2 in cultured skin fibroblasts, using 2 different monoclonal primary antibodies targeting conserved epitopes and normalized to histone H3. A third polyclonal antibody produced the same results but had higher background reactivity and is not shown. Each lane is a primary fibroblast line from a different adult individual. Fluorescent secondary antibodies were used to increase linear dynamic range for higher quantitative accuracy. **c,** Western blot for CIRBP with 3 different antibodies in 3 fibroblast lines per species. mAb=monoclonal antibody, pAb=polyclonal antibody. **d,** Stimulated emission depletion (STED) images of RPA2 and CIRBP localization in human and bowhead whale fibroblasts. Target protein in red, nuclear DAPI stain in blue. **e,** Western blot for CIRBP in fibroblasts isolated from various mammalian species.

**Extended Data Figure 8.**
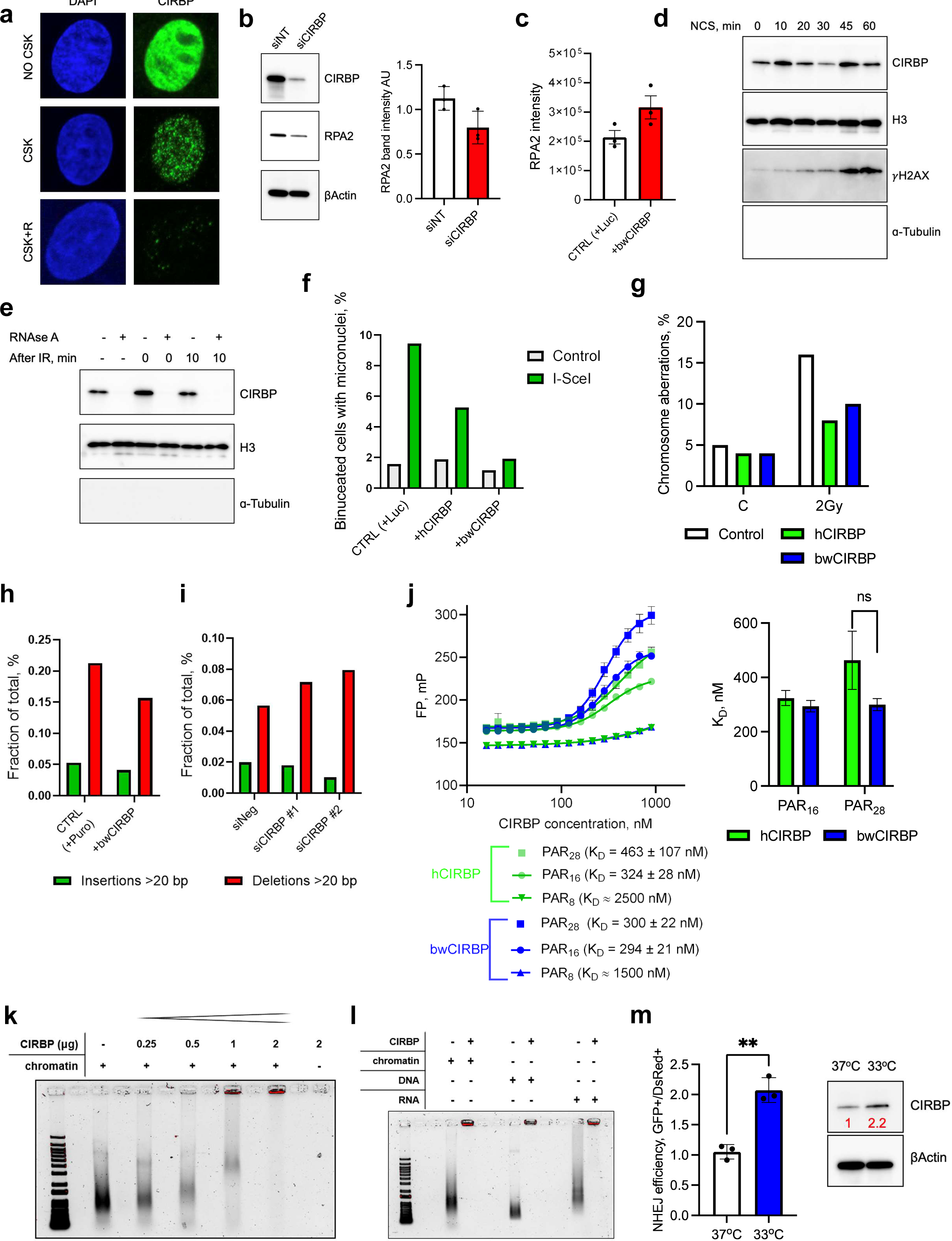
Analysis of CIRBP’s role in DNA DSB repair. **a,** CIRBP localization in whale cells. Before formaldehyde fixation, cells were pre-extracted with CSK buffer +/- RNAseA for 3min. After standard immunocytochemistry procedure images were collected using confocal microscope. **b,** Western blot of bowhead whale fibroblasts with knockdown of CIRBP (left panel) and band intensity quantification from 3 independent experiments (right panel) suggesting partial dependence of RPA2 protein abundance on CIRBP expression. **c,** Ion intensity by LC-MS of RPA2 in human fibroblasts with and without lentiviral overexpression of bwCIRBP (n=3 human cell lines). Error bars show mean +-SEM. **d,** DSBs induce CIRBP enrichment in chromatin. Exponentially growing cells were treated with neocarzinostatin (NCS) for the indicated period of time and lysed in CSK buffer to enrich chromatin-bound fraction. ɑ-Tubulin staining was used to verify the absence of cytoplasmic contamination in chromatin-bound fraction. **e,** DSBs induced by γ-irradiation lead to CIRBP enrichment in chromatin. This enrichment is promoted by RNA. Exponentially growing cells were treated with γ-irradiation and at the indicated period of time were lysed in CSK buffer with/without RNAse A to enrich proteins in chromatin-bound fraction. **f,** Overexpression of CIRBP decreases the percentage of binucleated cells containing micronuclei in human cells after I-Sce1-induced DSBs. Each bar indicates an experimental replicate. At least 150 binucleated cells were scored per condition. **g,** Frequency of chromosomal aberrations in human fibroblasts with and without CIRBP overexpression after 2Gy γ-irradiation. 100 metaphases were analyzed per sample. C=control untreated cells. **h,** Frequency of insertions and deletions >20 bp in NHEJ reporter constructs PCR-amplified from human fibroblasts with and without bwCIRBP overexpression after I-SceI expression. Insertion/deletion frequencies were determined from Nanopore sequencing data of PCR products and normalized within each sample to total frequency of all insertions or deletions. **i,** Frequency of insertions and deletions as shown in (**h**) but for bowhead whale fibroblasts with negative control or CIRBP-targeting siRNAs. **j,** Calculated dissociation constants (K_D_) and fluorescence polarization (FP) measurements for CIRBP proteins titrated into solutions containing a fixed concentration (3 nM) of fluorescently labeled PAR of various polymer lengths. **k,** EMSA of increasing amounts of recombinant human CIRBP incubated in vitro with 300 ng sheared chromatin from fibroblasts exposed to UVC and oxidative DNA damage as described in Methods. Chromatin was treated with Proteinase K but not RNAse. Nucleic acids are stained with SYBR Gold. Red overlay indicates saturated pixels. **l,** EMSA of 300 ng sheared purified genomic DNA, purified cellular RNA, or chromatin as described in (**k**) incubated in vitro with 5 μg rhCIRBP. **m,** Hypothermia promotes NHEJ efficiency in primary human fibroblasts (left panel). Cells were pre-incubated at 33°C for 2 days, co-transfected with I-SceI-digested NHEJ reporter and DsRed, and returned to the 33°C incubator. NHEJ efficiency was measured by flow cytometry 3 days following transfection (n=3). Western blot showing CIRBP upregulation in human cells exposed to 33°C for 2 days (right panel). Western blot images were analyzed in ImageLab software (Bio-Rad). Error bars represent mean ± SD. ** p<0.01 (Welch’s t-test).

**Extended Data Figure 9.**
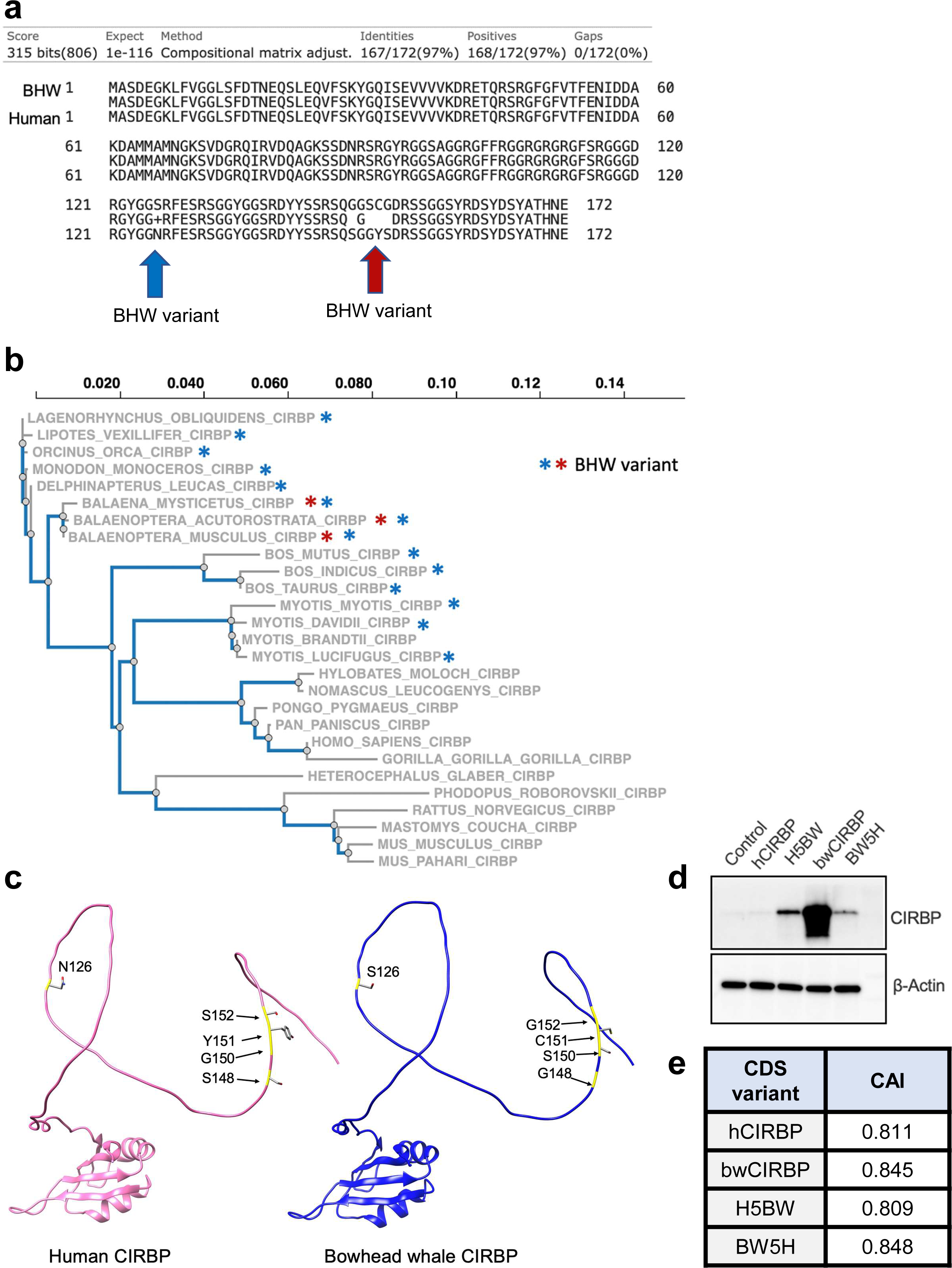
Analysis of bwCIRBP coding sequence mutations and protein expression levels. **a,** Comparison of amino acid sequences between human and bowhead whale CIRBP through BLAST analysis. **b,** Phylogenetic tree illustrating the relationships among CIRBP coding sequences from representative species with genome sequence information available. The asterisk indicates the presence of BHW-specific variants in the species. The colors indicate the position of variants shown in (**a**). **c,** SwissModel/AlphaFold models of human (left, pink) and bowhead whale (right, blue). Side chains of whale residues that diverge from human are shown, and their ribbon is colored yellow in the model. The key takeaway is that all the residues that differ between whale and human are in the C-terminal disordered region, whereas the N-terminal RNA recognition motif (RRM) is structured and conserved. **d,** Western blot abundance of bwCIRBP, hCIRBP, and reciprocal amino acid mutants overexpressed in human cells. **e,** Calculated codon adaptation index (CAI) for CIRBP coding sequence variants.

**Extended Data Figure 10.**
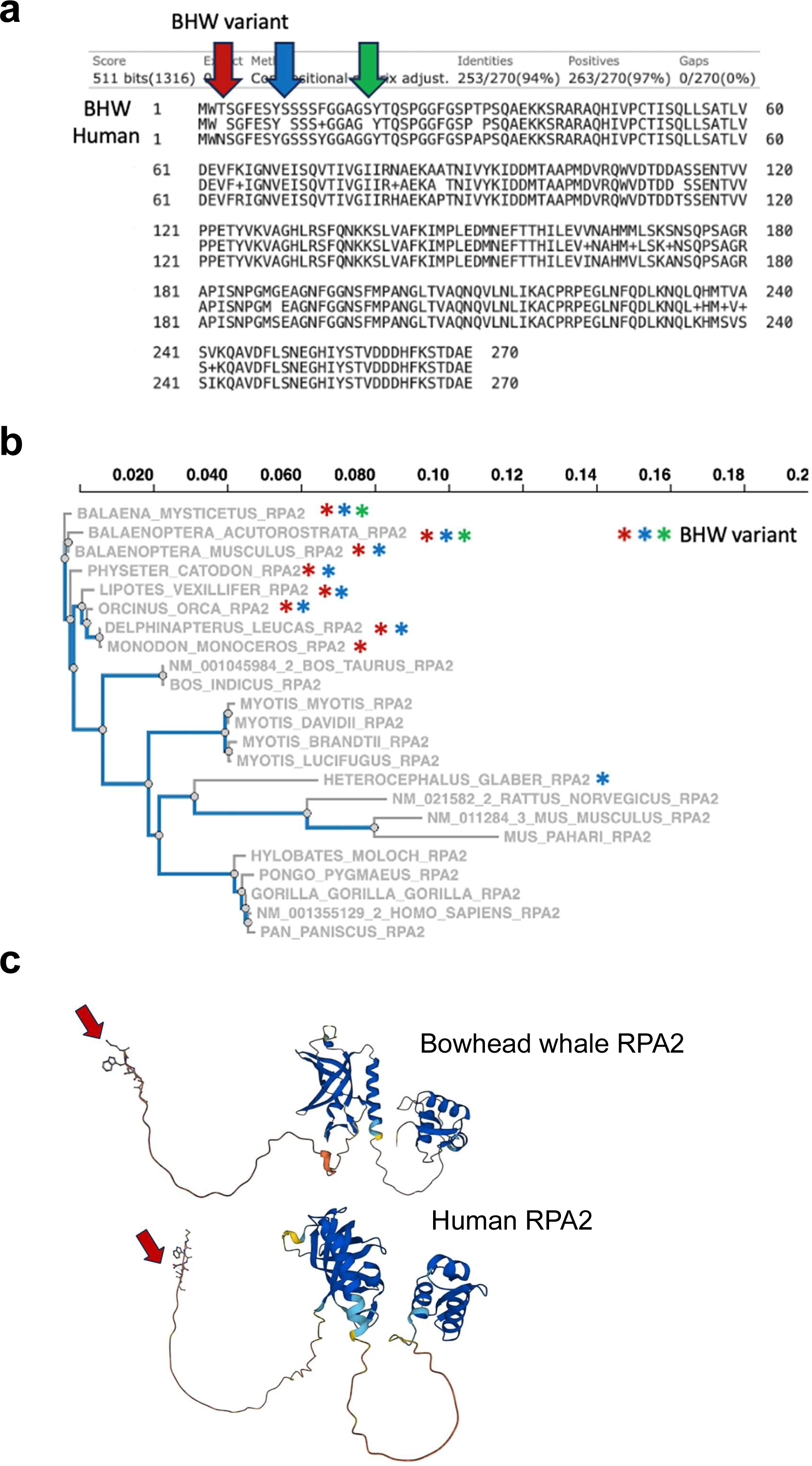
Analysis of bowhead whale RPA2 sequence. **a,** Comparison of amino acid sequences between human and bowhead whale RPA2 through BLAST analysis. **b,** Phylogenetic tree illustrating the relationships among RPA2 coding sequence from different representative species. The asterisk indicates the presence of BHW-specific variants in the species. The colors indicate the position of variants shown in (**a**). **c,** AlphaFold protein structures of human and bowhead whale RPA2 showing the position of the variants.

**Extended Data Figure 11.**
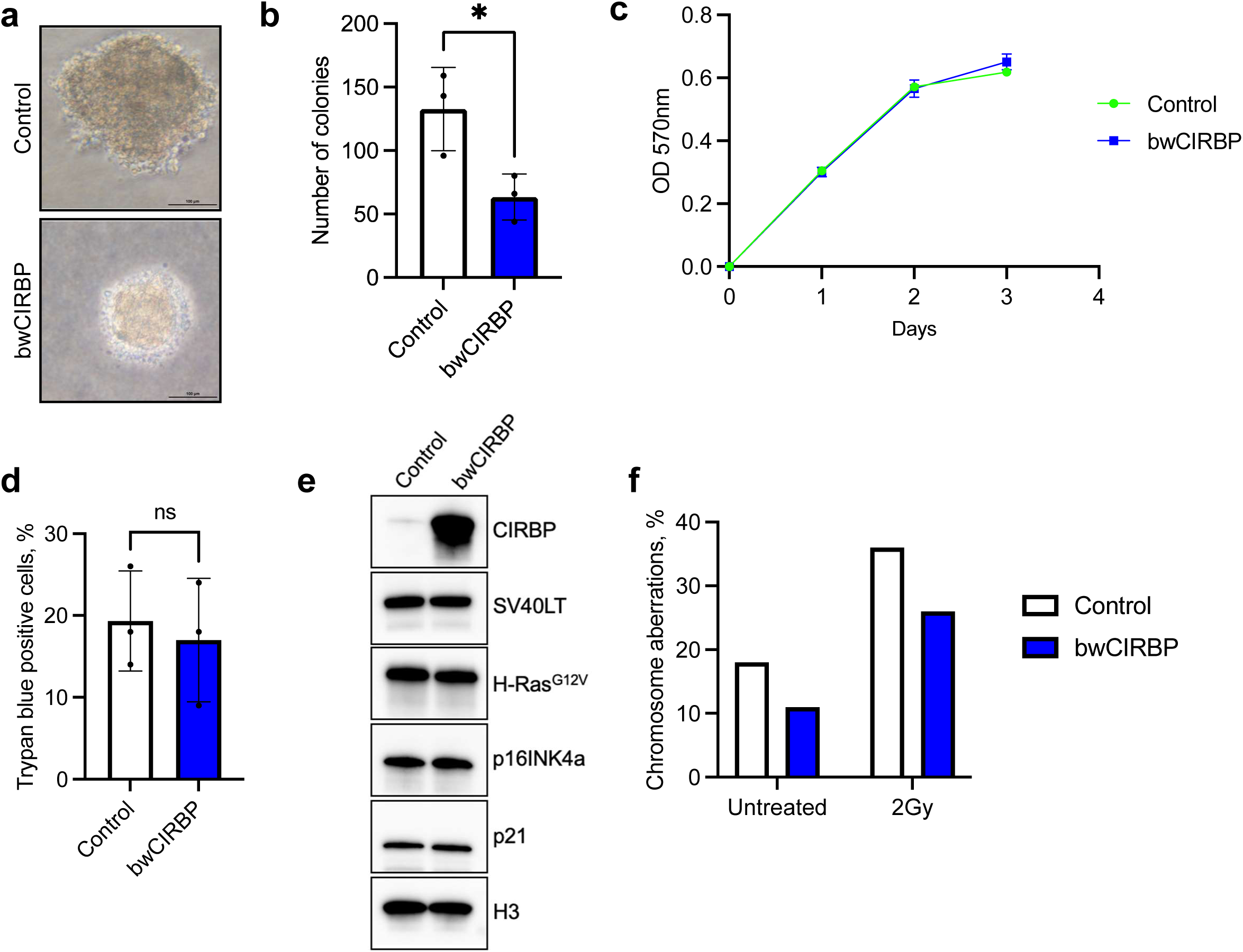
Bowhead whale CIRBP reduces anchorage-independent cell growth. **a,** Images of representative human transformed fibroblast colonies with and without bwCIRBP overexpression after 23 days of growth in soft agar. 20x magnification. Bar 100µm. **b,** Quantification of colonies after staining with nitro blue tetrazolium chloride. Colonies were counted using ImageJ software as described in Methods. Error bars represent SD. *p<0.05 (Welch’s t-test). **c,** Cell proliferation MTT assay. **d,** Trypan Blue exclusion test of cell viability. **e,** Western blot showing expression of LT, Ras, p16 and p21 after overexpression of bwCIRBP. **f,** Frequency of chromosomal aberrations in human transformed cells after bwCIRBP overexpression. 100 metaphases were analyzed per sample.

