## Supplementary Information & Figures for "DNA repair and anti-cancer mechanisms in the long-lived bowhead whale"

This file contains Supplementary Methods, Supplementary Notes, Supplementary Tables 5-7, and Supplementary Figures 1-5.

Supplementary Methods contain descriptions and catalogue numbers of Antibodies, kits, chemicals, and sequences for siRNAs and primers.

Supplementary Notes contains discussion of therapeutic hypothermia.

Supplementary Tables 5-7 contain statistical values for Figure 5.

Supplementary Figure 1 contains sequences of $p53^{-/-}$ clones.
Supplementary Figure 2 contains sequences of $Rb^{-/-}$ clones
Supplementary Figure 3 shows alignments of PTEN target amplicons and the CRISPR guide RNA sequence used for cross-species DSB repair fidelity assays.
Supplementary Figure 4 shows transfection efficiency of CRISPR RNP.
Supplementary Figure 5 shows gating strategy for the NHEJ assay.

#### Supplementary Methods

| Reagent | Source | Identifier or Sequence |
|---|---|---|
| ***Antibodies & Enzymes*** | | |
| Rabbit polyclonal anti-DNA PKcs | Abcam | ab70250 |
| Rabbit polyclonal anti-Ku80/XRCC5 | Novus Biologicals | NB100-503 |
| Ku70 (D10A7) Rabbit mAb | Cell Signaling Technology | 4588S |
| Rabbit polyclonal anti-Mre11 | Novus Biologicals | NB100-142 |
| Rabbit polyclonal anti-Rad50 | Novus Biologicals | NBP2-20054 |
| Rabbit polyclonal anti-Nbs1 | Novus Biologicals | NB100-143 |
| Rabbit polyclonal anti-PARP1 | Novus Biologicals | NBP2-13732 |
| SirT6 (D8D12) Rabbit mAb | Cell Signaling Technology | 12486S |
| RPA34 (RPA2) Mouse Monoclonal Antibody [Clone ID: OTI1H10] | OriGene | TA500765 |
| RPA34 (RPA2) Mouse Monoclonal Antibody [Clone ID: OTI7C12] | OriGene | TA500786 |
| Rabbit monoclonal [EPR2877Y] to RPA32/RPA2 | Abcam | ab76420 |
| Rabbit polyclonal anti-CIRBP | Proteintech | 10209-2-AP |
| Rabbit monoclonal [EPR18783] anti-CIRP | Abcam | ab191885 |
| Goat polyclonal anti-CIRP | Abcam | ab106230 |
| Mouse monoclonal [PAb 240] anti-p53 | Abcam | ab26 |
| Rabbit polyclonal anti-p53 | Abcam | ab131442 |
| Rabbit polyclonal anti-Rb | Abcam | ab226979 |

| | | |
|---|---|---|
| PTEN (D4.3) XP Rabbit mAb | Cell Signaling Technology | 9188S |
| Ras (G12V Mutant Specific) (D2H12) Rabbit mAb | Cell Signaling Technology | 14412S |
| Sv40 Large T Antigen (D1E9E) Rabbit mAb | Cell Signaling Technology | 15729S |
| Rabbit polyclonal anti-Histone H3 | Abcam | ab1791 |
| Rabbit polyclonal anti-beta actin | Abcam | ab8227 |
| Poly/Mono-ADP Ribose (E6F6A) Rabbit mAb | Cell Signaling Technology | 83732 |
| CtIP (D76F7) Rabbit mAb | Cell Signaling Technology | 9201S |
| Rabbit polyclonal anti-53BP1 | Abcam | ab172580 |
| Anti-phospho-Histone H2A.X (Ser139) Antibody, clone JBW301 | Sigma-Aldrich | 05-636 |
| Goat anti-mouse IgG H&L (HRP) | Abcam | ab6789 |
| Goat anti-rabbit IgG H&L (HRP) | Abcam | ab6721 |
| Rabbit Anti-Goat IgG H&L (HRP) | Abcam | ab6741 |
| Mouse Anti-Cyclobutane Pyrimidine Dimers (CPDs) mAb (Clone TDM-2) | Cosmo Bio | CAC-NM-DND-001 |
| Goat anti-Mouse IgG (H+L) Cross-Adsorbed Secondary Antibody, Biotin | ThermoFisher Scientific | 62-654-0 |
| Goat anti-Mouse IgG (H+L) Highly Cross-Adsorbed Secondary Antibody, Alexa Fluor™ 568 | ThermoFisher Scientific | A11031 |

| | | |
|---|---|---|
| Goat Anti-Rabbit IgG H&L (Alexa Fluor® 488) | Abcam | ab150077 |
| Starbright™ Blue 520 Goat Anti-Mouse IgG | Bio-Rad | 12005866 |
| Starbright™ Blue 520 Goat Anti-Rabbit IgG | Bio-Rad | 12005869 |

| | | |
|---|---|---|
| Streptavidin - HRP | ThermoFisher Scientific | 434323 |
| MS-grade trypsin | ThermoFisher Scientific | 90058 |
| rhRPA | Enzymax | 61 |
| rhCIRBP | Millipore | SRP6553 |
| Alt-R™ S.p. Cas9 Nuclease V3 | IDT | 1081058 |
| ***Kits*** | | |
| EpiQuik Nuclear Extraction Kit | Epigentek | OP-0002-1 |
| PARP Universal Colorimetric Assay Kit | Trevigen | 4677-096-K |
| Annexin V FLUOS Staining Kit | Roche | 1988549 |
| Annexin V Apoptosis Kit [FITC] | Novus Biologicals | NBP2-29373-100Tests |
| QIAamp DNA Blood Mini Kit | Qiagen | 51104 |
| Q5 Site-Directed Mutagenesis Kit | New England Biolabs | E0554 |
| One Shot TOP10 Chemically Competent Cells | ThermoFisher Scientific | C404006 |
| Zero Blunt TOPO PCR Cloning Kit for Sequencing | ThermoFisher Scientific | 45-003-1 |
| TOPO TA Cloning Kit for Subcloning | ThermoFisher Scientific | 450641 |

| | | |
|---|---|---|
| Phusion Hot Start Flex DNA Polymerase | New England Biolabs | M0535 |
| QIAprep Spin Miniprep Kit | Qiagen | 27106 |
| EndoFree Plasmid Maxi Kit | Qiagen | 12362 |
| NHDF Nucleofector® Kit | Lonza | VPD-1001 |
| Quick-RNA™ MiniPrep | Zymo Research | R1054 |
| Dual-Luciferase Reporter 1000 Assay System | Promega | E1980 |
| Protamine Sulfate Coated ELISA plate | Cosmo Bio | CSR-NM-MA-P002 |
| KAPA 2G Robust HotStart ReadyMix | Roche | 07961375001 |
| S-Trap Micro Column | Protifi | C02-micro-80 |
| Wizard Genomic DNA Purification Kit | Promega | A1120 |
| QIAquick Gel Extraction Kit | Qiagen | 28706 |
| NEBNext Q5 Hot Start HiFi PCR Master Mix | New England Biolabs | M0543L |
| ***Chemicals*** | | |
| Cytochalasin B from Drechslera dematioidea | Sigma-Aldrich | C2743 |
| Methylene Blue | Sigma-Aldrich | M9140 |
| Agarose, low gelling temperature | Sigma-Aldrich | A9414 |
| 50% Hydrogen Peroxide | Fisher Scientific | H341 |
| Antifade mounting medium with DAPI | Vector laboratories | H-1200 |
| Acridine Orange | ThermoFisher Scientific | A1301 |

| | | |
|---|---|---|
| Geneticin™ Selective Antibiotic (G418 Sulfate) | ThermoFisher Scientific | 10131035 |
| Puromycin | ThermoFisher Scientific | A1113803 |
| Etoposide | Sigma-Aldrich | E1383 |
| N-Nitroso-N-ethylurea,ISOPAC® | Sigma-Aldrich | N3385 |
| 6-thioguanine | Chem-Impex | 01491 |
| Nitro blue tetrazolium chloride | Thermo Fisher | N6495 |
| TDRL-505 | Sigma-Aldrich | 5305350001 |

| | | |
|---|---|---|
| Alt-R® Cas9 Electroporation Enhancer | IDT | 1075915 |
| SYBR Gold | ThermoFisher Scientific | S11494 |
| Lipofectamine™ RNAiMAX Transfection Reagent | ThermoFisher Scientific | 13778150 |
| RNase A, DNase and protease-free (10 mg/mL) | ThermoFisher Scientific | EN0531 |
| Neocarzinostatin from Streptomyces carzinostaticus | Sigma-Aldrich | N9162-100UG |
| Bleomycin Sulfate, Streptomyces verticillus | EMD Millipore | 203401-10MG |
| VECTASHIELD PLUS Antifade Mounting Medium with DAPI | Vector Laboratories | H-2000-10 |
| ***siRNA*** | | |
| Non-target | Dharmacon | UGGUUUACAUGUCGACUAA |
| anti-bwCIRBP | Dharmacon | GACAGATCTCAGAAGTGGTGGTCGT |
| Silencer™ Select Negative Control No. 2 siRNA | ThermoFisher Scientific | 4390846 |
| Silencer™ Select Custom anti-bwCIRBP #1 | ThermoFisher Scientific | CUACGACAGUUACGCUACA |

| | | |
|---|---|---|
| Silencer™ Select Custom anti-bwCIRBP #2 | ThermoFisher Scientific | AGCAGGUCUUCUCAAAAUA |
| ***Primers & Probes*** | | |
| Pem-F | IDT | GCTAAGTGCTTAGTAAAGCAATAGACTGCAT |
| Pem-R | IDT | GGACTAGTAATTGTTTAACATGTGGGAAGTT |
| PTEN ddPCR probe | IDT | /56-FAM/ACCGCCAAG/ZEN/TCCAGAGCCAT/3IABkFQ/ |
| UCE.359 ddPCR probe | IDT | /5HEX/ATGAGACGG/ZEN/GAGACAAGTGATGCT/3IABkFQ/ |
| UCE.359-F | IDT | ATCTGAGACTTGTGACAT |
| UCE.359-R | IDT | GTGTTAATTGGTAATGACTATT |
| *RB1* KO up-F | IDT | TCGGGAAGAGACAGGTTCAG |
| *RB1* KO up-R | IDT | TCGCTCCCGAACCCAAA |
| *RB1* KO down 1-F | IDT | ACTAAAAGGTCTGAGTGAGTGC |
| *RB1* KO down 1-R | IDT | CTCATATGAGAAGCCCAGAAGG |
| *RB1* KO down 2-F | IDT | ACTAAAAGGTCTGAGTGAGTGC |
| *RB1* KO down 2-R | IDT | CTTTCGAAGCAGAACTCCAAAC |
| *TP53* KO up-F | IDT | GCAGAAATAGGTAGACACTGAGGAA |
| *TP53* KO up-R | IDT | ACGGGAGAAACGAGAGTCAGAA |
| *TP53* KO down 1-F | IDT | CTTCCTGTGTTGGAAAGCCATG |
| *TP53* KO down 1-R | IDT | CACCAAGCAGAGGTTTAAGCAC |
| *TP53* KO down 2-F | IDT | CTTCCTGTGTTGGAAAGCCATG |
| *TP53* KO down 2-R | IDT | CTGCCCCTTCTTAGACTTCAGG |
| *PTEN* KO up-F | IDT | CAGTCGCTGCAACCATCCA |
| *PTEN* KO up-R | IDT | ATCCGACCAGGGTTAAATGTCATAG |
| *PTEN* KO down 1-F | IDT | AAGAGGGTCATTTAAAAGGCCC |
| *PTEN* KO down 1-R | IDT | TCTGACACAATGTCCTATTGCC |
| *PTEN* KO down 2-F | IDT | TAGGGCGAATGGAATCCTAAAC |
| *PTEN* KO down 2-R | IDT | TCTGACACAATGTCCTATTGCC |
| *PTEN* fidelity-F | IDT | TGGCTGCTGAGGAGAAGC |
| *PTEN* fidelity-R | IDT | GCATCCGTCTACTCCCACG |
| *PTEN* fidelity-R (cow SNP) | IDT | GCATCCGTCTATTCCCACG |
| *PTEN* ddPCR-F | IDT | TTCTGCCATCTCTCTCC |
| *PTEN* ddPCR-R | IDT | CAGGTCAAGTCTAAGTCG |

| | | |
|---|---|---|
| Lego-iC2-hCIRBP-F | IDT | CCGCGAATTCGCCACCATGGCATCAGATGAAGGCAAAC |
| Lego-iC2-hCIRBP-R | IDT | CCGCGCGGCCGCTTACTCGTTGTGTGTAGCGTAACTGTCATAACTGTCTC |
| Lego-iC2-hCIRBPopt-F | IDT | CCGCGAATTCGCCACCATGGCCAGCGACGAAG |
| Lego-iC2-hCIRBPopt-R | IDT | CCGCGCGGCCGCTCACTCGTTGTGTGTGGCATAAGAGTCATAGG |
| Lego-iC2-bwCIRBP-F | IDT | CCGCGAATTCGCCACCATGGCATCAGATGAGGGC |
| Lego-iC2-bwCIRBP-R | IDT | CCGCGCGGCCGCTTACTCGTTGTGTGTAGCGTAACTGTCGTAGC |
| ***CRISPR Guide RNAs*** | | |
| Alt-R® CRISPR-Cas9 tracrRNA | IDT | 1072533 |

| | | |
|---|---|---|
| Alt-R® CRISPR-Cas9 tracrRNA, ATTO™ 550 | IDT | 1075928 |
| *RB1* KO crRNA 1 | IDT | GCTGCTCTACGGGGGGTTTT |
| *RB1* KO crRNA 2 | IDT | GGCTGCTCTACGGGGGGTTT |
| *RB1* KO crRNA 3 | IDT | AGACACCAGCTGATACTACG |
| *TP53* KO crRNA 1 | IDT | GTCACAGGCAGAACTCGGCG |
| *TP53* KO crRNA 2 | IDT | CGTGGAGCCCCCTCTGAGTC |
| *TP53* KO crRNA 3 | IDT | AAATAGGGGAGTGATAACCG |
| *PTEN* KO crRNA 1 | IDT | GCTAACGATCTCTTTGATGA |
| *PTEN* KO crRNA 2 | IDT | AGATCGTTAGCAGAAACAAA |
| *PTEN* KO crRNA 3 | IDT | ATTCTCTGGATCAGAGTCAG |
| *PTEN* fidelity crRNA | IDT | GCTAACGATCTCTTTGATGA |
| ***Plasmids*** | | |
| pPB-SV40 large T (LT) | [1] | |
| pPB-$LT^{K1}$ | [1] | |
| pPB-$LT^{\Delta 434-444}$ | [1] | |
| pPB- SV40 small T (ST) + LT | [1] | |
| pPB-H-Ras V12 | [1] | |
| pPB-*hTERT* | [1] | |
| Super PiggyBac Transposase Expression Vector | System Biosciences | PB210PA-1 |
| pRP[Exp]-Neo-CMV-hCIRBP | VectorBuilder | VB210330-1269gmt |

| | | |
|---|---|---|
| pRP[Exp]-Neo-CMV-hCIRBPopt | VectorBuilder | VB230901-1324ctc |
| pRP[Exp]-Neo-CMV-bwCIRBP | VectorBuilder | VB210330-1268bpc |
| pRP[Exp]-Neo-CMV-bwCIRBP-RGG 9R/A | VectorBuilder | VB220320-1032yqx |
| pRP[Exp]-Neo-CMV-Puro | VectorBuilder | VB220318-1212uym |
| pCMV-Luc | [2] | |
| pCMV-RL | Promega | E2261 |
| NHEJ reporter construct | [3] | |
| HR reporter construct | [3] | |
| pCMV-ISce1 | [3] | |
| pDsRed | [3] | |

| | | |
|---|---|---|
| LeGO-iC2-Luciferase | This manuscript | |
| LeGO-iC2-hCIRBP | This manuscript | |
| LeGO-iC2-hCIRBPopt | This manuscript | |
| LeGO-iC2-bwCIRBP | This manuscript | |
| pp53-TA-Luc | Clontech (Takara) | 631914 |
| pE2F-TA-Luc | Clontech (Takara) | 631914 |
| pRb-TA-Luc | Clontech (Takara) | 631914 |
| Lego-iC2 | Addgene | 27345 |
| pVSV-G | Addgene | 138479 |
| psPAX2 | Addgene | 12260 |
| pmaxGFP | Lonza | VPD-1001 |
| ***Cell Lines*** | | |
| 14B8SF | [4] | Primary dermal fibroblasts isolated by the authors from adult female bowhead whale |
| 14B10SF | [4] | Primary dermal fibroblasts isolated by the authors from adult male bowhead whale |
| 14B11SF | [4] | Primary dermal fibroblasts isolated by the authors from adult male bowhead whale |
| 18B2SF | [5] | Primary dermal fibroblasts isolated by the authors from adult female bowhead whale |

| | | |
|---|---|---|
| 18B9SF | [5] | Primary dermal fibroblasts isolated by the authors from adult male bowhead whale |
| 18B12SF | [5] | Primary dermal fibroblasts isolated by the authors from adult female bowhead whale |
| NHDF | ATCC | PCS-201-012, Lot 64540954, normal adult male human primary dermal fibroblasts |
| HDF637 | ATCC | PCS-201-012, Lot 63792061, normal adult male human primary dermal fibroblasts |
| YAHSF | ATCC | CRL-2691, normal adult male human primary dermal fibroblasts |
| MJ | | Normal neonatal male foreskin primary fibroblasts provided by collaborator |
| WI-38 | ATCC | CCL-75, primary female fetal lung fibroblasts |
| BT1SF | | Primary dermal fibroblasts isolated by the authors from adult *Bos taurus* |
| BT2SF | | Primary dermal fibroblasts isolated by the authors from adult *Bos taurus* |
| BT3SF | | Primary dermal fibroblasts isolated by the authors from adult *Bos taurus* |
| WTMSF8 | | Primary dermal fibroblasts isolated by the authors from adult wild-caught *Mus musculus* |
| WTMSF9 | | Primary dermal fibroblasts isolated by the authors from adult wild-caught *Mus musculus* |
| WTMSF10 | | Primary dermal fibroblasts isolated by the authors from adult wild-caught *Mus musculus* |
| BL6AMSF2 | | Primary dermal fibroblasts isolated by the authors from adult male, normal C57BL/6 *Mus musculus* |
| Hippopotamus primary fibroblasts | San Diego Zoo Wildlife Alliance | |
| Common dolphin fibroblasts | San Diego Zoo Wildlife Alliance | |
| Humpback whale primary fibroblasts | San Diego Zoo Wildlife Alliance | |
| Bottlenose dolphin primary fibroblasts | | Primary fibroblasts isolated by the authors from bottlenose dolphin tissues collected by Georgia Aquarium through Tara Harrison (Exotic Species Cancer Research Alliance) |

| | | |
|---|---|---|
| California sea lion primary fibroblasts |  | Primary fibroblasts by the authors from California sea lion tissues collected by the Marine Mammal Care Center Los Angeles under Institutional Animal Care and Use Committee oversight and National Marine Fisheries Service permit number 21636. |
| Lenti-X 293T | Takara | 632180 |

#### Supplementary Note

#### Therapeutic hypothermia

Beneficial effects of cold as a therapeutic agent have been noted for a long time with brief cold-water immersion believed to promote health and hardening in Nordic cultures. Currently, whole body cryotherapy is widely used in sports medicine to reduce inflammation and facilitate recovery after exercise or injury[1]. While molecular mechanisms responsible for the beneficial effects of cryotherapy are largely unknown, we speculate that increased CIRBP expression may contribute to health benefits by facilitating DNA repair. Indeed, when we tested whether a hypothermia-mediated increase in human CIRBP could affect DSB repair efficiency, we observed an increase in NHEJ efficiency (Extended Data Figure 8m).

Cold showers provide a mild and transient reduction in body temperature, whereas artificial reduction of body temperature, or therapeutic hypothermia, is used clinically during surgery[2]. Interestingly, recovery from therapeutic hypothermia is compromised in CIRBP-KO rats and improved in rats with transgenic CIRBP overexpression[3]. Similar protective effects of CIRBP during hypothermia have been reported in brain[4], kidney[5] intestine[6], and cold-preserved hearts[7]. Given the dramatically enhanced protein production driven by bwCIRBP and hCIRBP mutants, local delivery of such CIRBP variants (e. g. through mRNA/DNA vector injection) could hold promise as strategies to improve tissue recovery from surgery or organ transplant.

### Supplementary Tables

#### Supplementary Table 5. ANOVA for Figure 5a

| Uncorrected Fisher's LSD | Mean Diff. | 95.00% CI of diff. | Below threshold? | Summary | Individual P Value |
|---|---|---|---|---|---|
| Unmodified | | | | | |
| CRISPR vs. CRISPR +TDRL-505 | 0.05767 | -0.03855 to 0.1539 | No | ns | 0.1232 |
| CRISPR vs. CRISPR + RPA | -0.1992 | -0.3692 to -0.02927 | Yes | * | 0.0371 |
| Substitutions | | | | | |
| CRISPR vs. CRISPR +TDRL-505 | 0.03891 | 0.03454 to 0.04328 | Yes | *** | 0.0007 |
| CRISPR vs. CRISPR + RPA | 0.01569 | -0.001850 to 0.03323 | No | ns | 0.0614 |
| Insertions >1 bp | | | | | |
| CRISPR vs. CRISPR +TDRL-505 | 0.01742 | -0.01002 to 0.04486 | No | ns | 0.1119 |
| CRISPR vs. CRISPR + RPA | 0.001321 | -0.02206 to 0.02470 | No | ns | 0.8306 |
| Insertions ≤1 bp | | | | | |
| CRISPR vs. CRISPR +TDRL-505 | 0.005787 | -0.01362 to 0.02519 | No | ns | 0.328 |
| CRISPR vs. CRISPR + RPA | 0.01687 | 0.01075 to 0.02299 | Yes | ** | 0.007 |
| Complex Deletions | | | | | |
| CRISPR vs. CRISPR +TDRL-505 | 0.0279 | 0.02401 to 0.03180 | Yes | ** | 0.0011 |
| CRISPR vs. CRISPR + RPA | 0.04349 | 0.009846 to 0.07713 | Yes | * | 0.0308 |
| Deletions ≤20bp | | | | | |
| CRISPR vs. CRISPR +TDRL-505 | -0.1519 | -0.2873 to -0.01646 | Yes | * | 0.0404 |
| CRISPR vs. CRISPR + RPA | 0.1241 | 0.03022 to 0.2180 | Yes | * | 0.0296 |
| Deletions >20 bp | | | | | |
| CRISPR vs. CRISPR +TDRL-505 | 0.004173 | -0.001390 to 0.009736 | No | ns | 0.0841 |
| CRISPR vs. CRISPR + RPA | -0.002245 | -0.03359 to 0.02910 | No | ns | 0.7871 |

| Test details | Mean 1 | Mean 2 | Mean Diff. | SE of diff. | N1 | N2 | t | DF |
|---|---|---|---|---|---|---|---|---|
| Unmodified | | | | | | | | |
| CRISPR vs. CRISPR +TDRL-505 | 0.3298 | 0.2722 | 0.05767 | 0.0224 | 3 | 3 | 2.579 | 2 |
| CRISPR vs. CRISPR + RPA | 0.3298 | 0.5291 | -0.1992 | 0.0395 | 3 | 3 | 5.044 | 2 |
| Substitutions | | | | | | | | |
| CRISPR vs. CRISPR +TDRL-505 | 0.07402 | 0.0351 | 0.03891 | 0.001 | 3 | 3 | 38.32 | 2 |
| CRISPR vs. CRISPR + RPA | 0.07402 | 0.05833 | 0.01569 | 0.0041 | 3 | 3 | 3.849 | 2 |
| Insertions >1 bp | | | | | | | | |
| CRISPR vs. CRISPR +TDRL-505 | 0.03177 | 0.01434 | 0.01742 | 0.0064 | 3 | 3 | 2.732 | 2 |
| CRISPR vs. CRISPR + RPA | 0.03177 | 0.03044 | 0.001321 | 0.0054 | 3 | 3 | 0.2431 | 2 |
| Insertions ≤1 bp | | | | | | | | |
| CRISPR vs. CRISPR +TDRL-505 | 0.05978 | 0.05399 | 0.005787 | 0.0045 | 3 | 3 | 1.283 | 2 |
| CRISPR vs. CRISPR + RPA | 0.05978 | 0.04291 | 0.01687 | 0.0014 | 3 | 3 | 11.86 | 2 |
| Complex Deletions | | | | | | | | |
| CRISPR vs. CRISPR +TDRL-505 | 0.0917 | 0.0638 | 0.0279 | 0.0009 | 3 | 3 | 30.83 | 2 |
| CRISPR vs. CRISPR + RPA | 0.0917 | 0.04821 | 0.04349 | 0.0078 | 3 | 3 | 5.562 | 2 |
| Deletions ≤20bp | | | | | | | | |
| CRISPR vs. CRISPR +TDRL-505 | 0.3699 | 0.5218 | -0.1519 | 0.0315 | 3 | 3 | 4.826 | 2 |
| CRISPR vs. CRISPR + RPA | 0.3699 | 0.2458 | 0.1241 | 0.0218 | 3 | 3 | 5.687 | 2 |
| Deletions >20 bp | | | | | | | | |
| CRISPR vs. CRISPR +TDRL-505 | 0.04299 | 0.03881 | 0.004173 | 0.0013 | 3 | 3 | 3.228 | 2 |
| CRISPR vs. CRISPR + RPA | 0.04299 | 0.04523 | -0.002245 | 0.0073 | 3 | 3 | 0.3082 | 2 |

#### Supplementary Table 6. ANOVA for Figure 5b

| Uncorrected Fisher's LSD | Mean Diff. | 95.00% CI of diff. | Below threshold? | Summary | Individual P Value | | | |
|---|---|---|---|---|---|---|---|---|
| CRISPR - CRISPR +TDRL-505 | | | | | | | | |
| Deletions >3 bp | -0.03388 | -0.1015 to 0.03374 | No | ns | 0.3007 | | | |
| Insertions >1 bp | 0.02337 | -0.04425 to 0.09100 | No | ns | 0.4708 | | | |
| Unmodified | 0.2158 | 0.1482 to 0.2834 | Yes | **** | <0.0001 | | | |
| Deletions ≤3 bp | -0.06979 | -0.1374 to -0.002167 | Yes | * | 0.044 | | | |
| Insertions ≤1 bp | -0.2277 | -0.2954 to -0.1601 | Yes | **** | <0.0001 | | | |
| Substitutions | 0.08478 | 0.01715 to 0.1524 | Yes | * | 0.0176 | | | |
| Complex Deletions | 0.007456 | -0.06017 to 0.07508 | No | ns | 0.8165 | | | |
| Test details | Mean 1 | Mean 2 | Mean Diff. | SE of diff. | N1 | N2 | t | DF |
| CRISPR - CRISPR +TDRL-505 | | | | | | | | |
| Deletions >3 bp | 0.021 | 0.05489 | -0.03388 | 0.03153 | 3 | 3 | 1.075 | 14 |
| Insertions >1 bp | 0.05066 | 0.02728 | 0.02337 | 0.03153 | 3 | 3 | 0.7413 | 14 |
| Unmodified | 0.4719 | 0.2561 | 0.2158 | 0.03153 | 3 | 3 | 6.844 | 14 |
| Deletions ≤3 bp | 0.08296 | 0.1528 | -0.06979 | 0.03153 | 3 | 3 | 2.214 | 14 |
| Insertions ≤1 bp | 0.2444 | 0.4722 | -0.2277 | 0.03153 | 3 | 3 | 7.223 | 14 |
| Substitutions | 0.1069 | 0.02209 | 0.08478 | 0.03153 | 3 | 3 | 2.689 | 14 |
| Complex Deletions | 0.02218 | 0.01473 | 0.007456 | 0.03153 | 3 | 3 | 0.2365 | 14 |

#### Supplementary Table 7. ANOVA for Figure 5c

| Uncorrected Fisher's LSD | Mean Diff. | 95.00% CI of diff. | Below threshold? | Summary | Individual P Value | | | |
|---|---|---|---|---|---|---|---|---|
| CRISPR - CRISPR + bwCIRBP | | | | | | | | |
| Deletions >20 bp | -0.001176 | -0.04744 to 0.04509 | No | ns | 0.9573 | | | |
| Insertions >1 bp | 0.0102 | -0.03607 to 0.05647 | No | ns | 0.6437 | | | |
| Deletions ≤20bp | 0.007581 | -0.03869 to 0.05385 | No | ns | 0.7305 | | | |
| Insertions 1 bp | 0.02677 | -0.01950 to 0.07303 | No | ns | 0.2351 | | | |
| Substitutions | -0.003783 | -0.05005 to 0.04249 | No | ns | 0.8633 | | | |
| Complex Deletions | 0.007353 | -0.03892 to 0.05362 | No | ns | 0.7383 | | | |
| Unmodified | -0.04694 | -0.09321 to -0.0006692 | Yes | * | 0.0472 | | | |
| Test details | Mean 1 | Mean 2 | Mean Diff. | SE of diff. | N1 | N2 | t | DF |
| CRISPR - CRISPR + bwCIRBP | | | | | | | | |
| Deletions >20 bp | 0.0616 | 0.06277 | -0.001176 | 0.02157 | 3 | 3 | 0.0545 | 14 |
| Insertions >1 bp | 0.01988 | 0.009684 | 0.0102 | 0.02157 | 3 | 3 | 0.4726 | 14 |
| Deletions ≤20bp | 0.4482 | 0.4406 | 0.007581 | 0.02157 | 3 | 3 | 0.3514 | 14 |
| Insertions 1 bp | 0.05208 | 0.02531 | 0.02677 | 0.02157 | 3 | 3 | 1.241 | 14 |
| Substitutions | 0.04352 | 0.0473 | -0.003783 | 0.02157 | 3 | 3 | 0.1753 | 14 |
| Complex Deletions | 0.06763 | 0.06028 | 0.007353 | 0.02157 | 3 | 3 | 0.3408 | 14 |
| Unmodified | 0.3071 | 0.354 | -0.04694 | 0.02157 | 3 | 3 | 2.176 | 14 |

**a**

#### *TP53* upstream

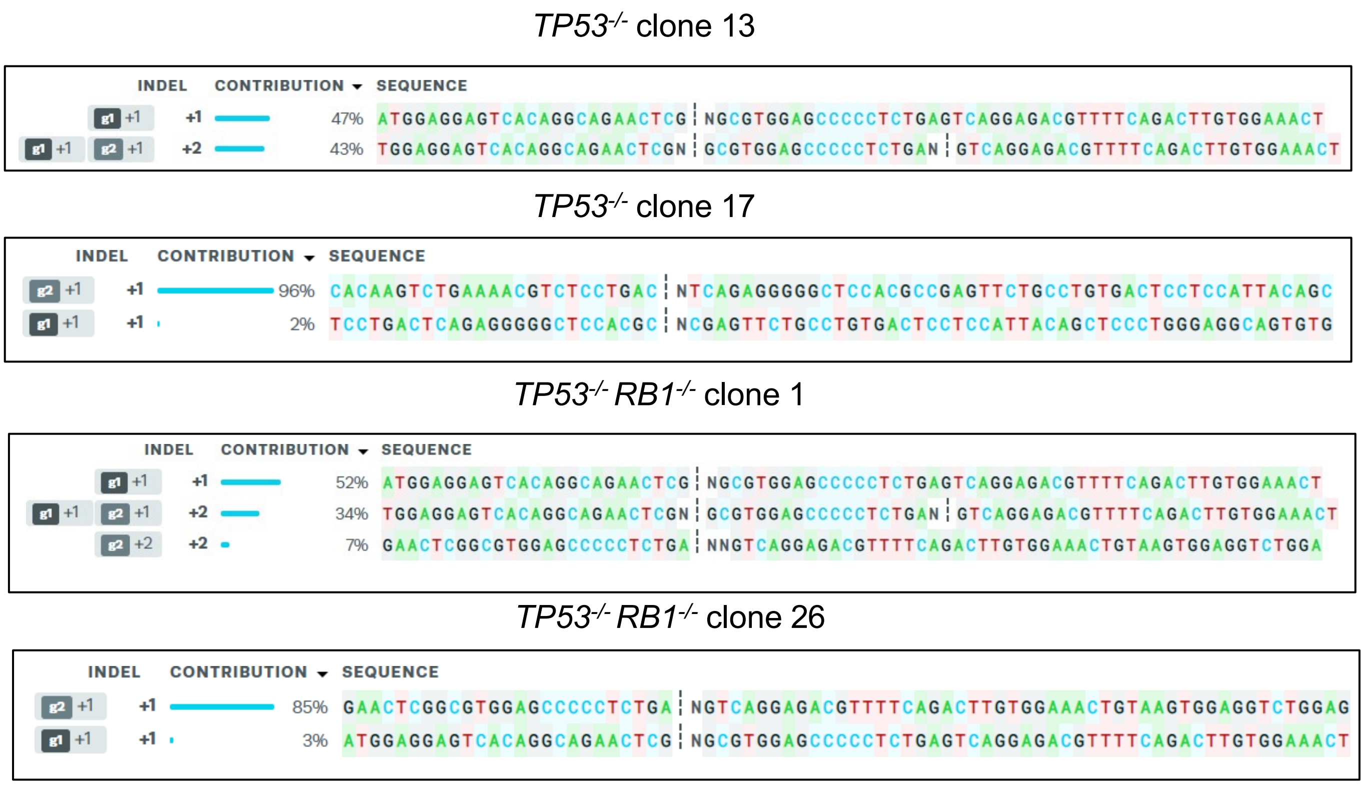

**b**

#### *TP53* downstream

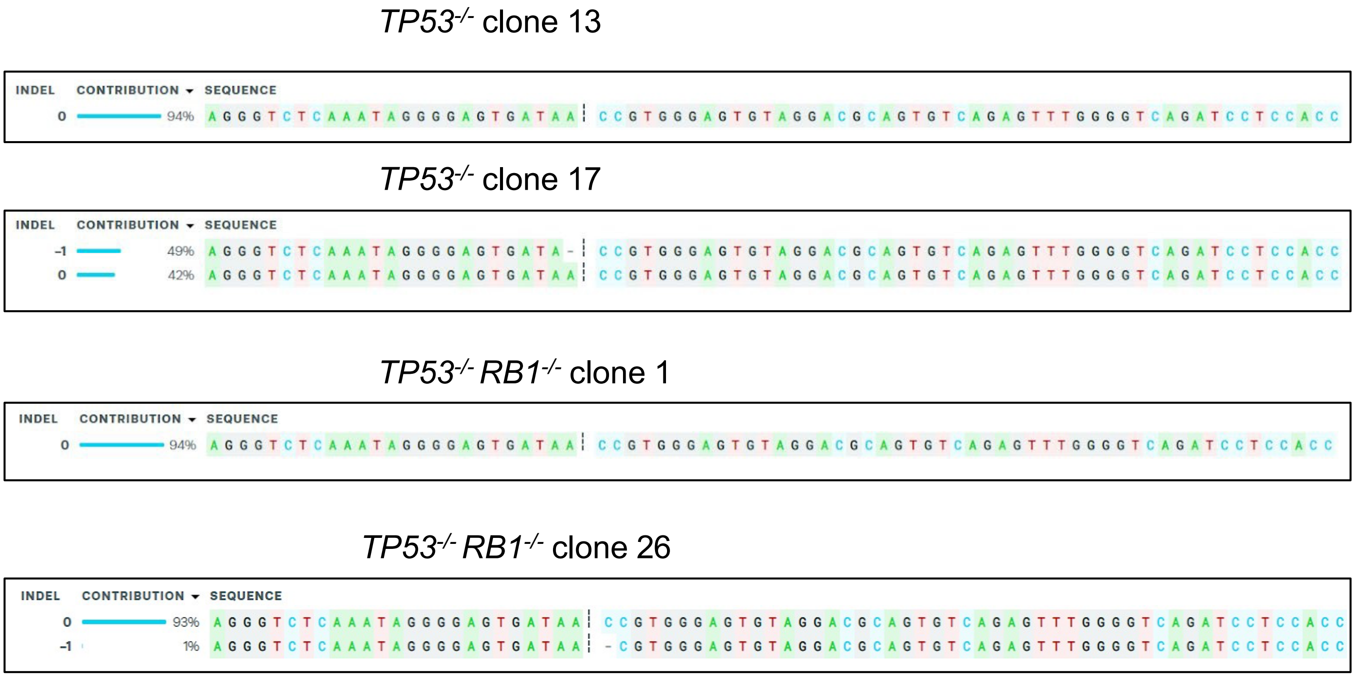

**Supplementary Figure 1.** a) Results of Synthego ICE deconvolution of Sanger sequencing traces from PCR products amplified from the upstream *TP53* target site for the indicated cell lines. Estimated allele frequencies close to 100% are considered to indicate homozygous alleles, while the presence of 2 alleles with estimated frequencies close to 50% indicate heterozygosity at this site. Predicted alleles with single-digit frequencies are considered to result from background noise in Sanger traces rather than genuine alleles. b) Results of Synthego ICE deconvolution of Sanger sequencing traces from PCR products amplified from the downstream *TP53* target site for the indicated cell lines.

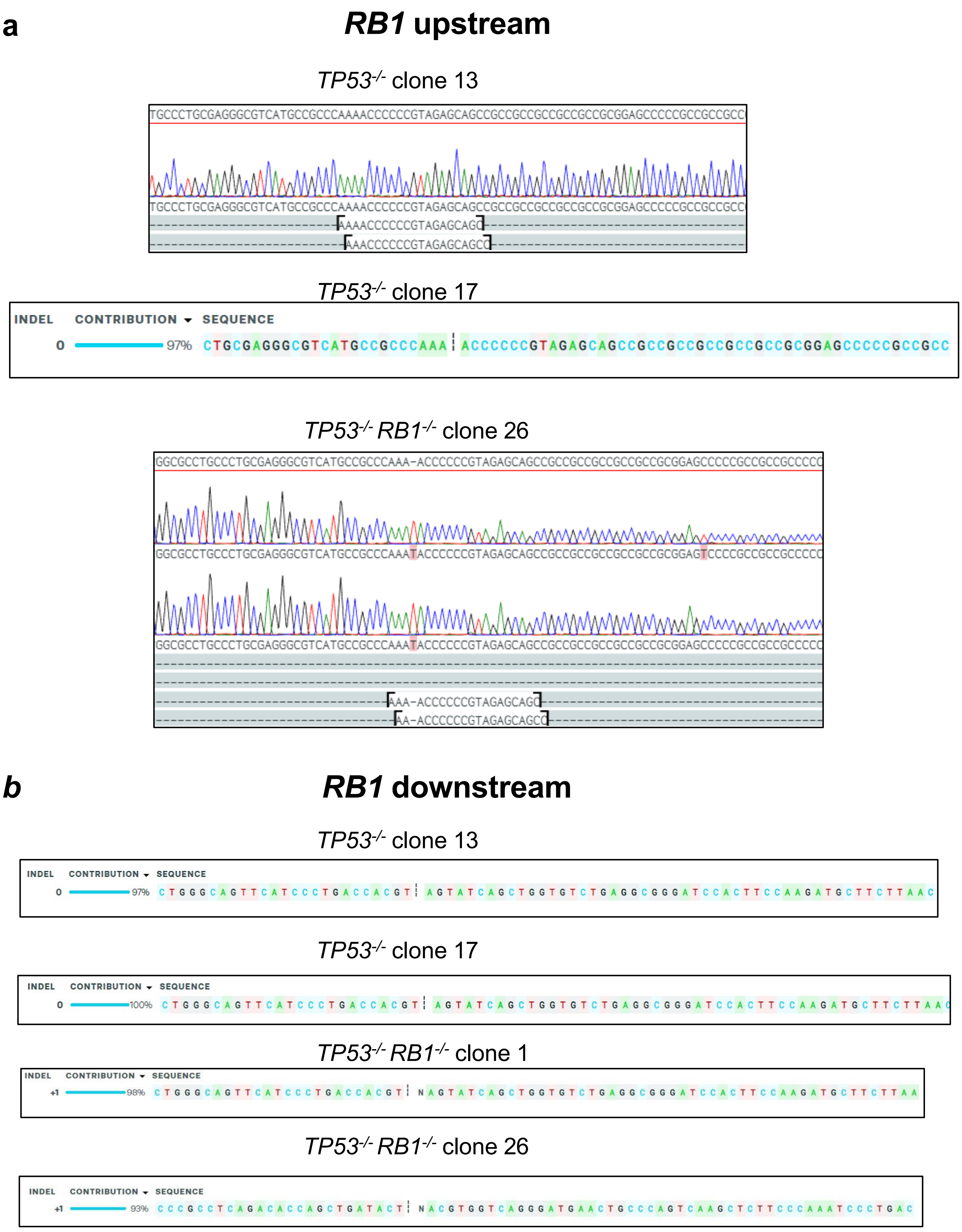

a
RB1 upstream
TP53-/- clone 13
TP53-/- clone 17
INDEL
CONTRIBUTION
SEQUENCE
0
97%
TP53-/- RB1-/- clone 26
b
RB1 downstream
TP53-/- clone 13
INDEL
CONTRIBUTION
SEQUENCE
0
97%
TP53-/- clone 17
INDEL
CONTRIBUTION
SEQUENCE
0
100%
TP53-/- RB1-/- clone 1
INDEL
CONTRIBUTION
SEQUENCE
+1
98%
TP53-/- RB1-/- clone 26
INDEL
CONTRIBUTION
SEQUENCE
+1
93%

**Supplementary Figure 2.** a) Results of Synthego ICE deconvolution of Sanger sequencing traces from PCR products amplified from the upstream *RB1* target site for the indicated cell lines. Estimated allele frequencies close to 100% are considered to indicate homozygous alleles, while the presence of 2 alleles with estimated frequencies close to 50% indicate heterozygosity at this site. Predicted alleles with single-digit frequencies are considered to result from background noise in Sanger traces rather than genuine alleles. For PCR products that failed Synthego ICE analysis, individual alleles were sequenced through TOPO cloning + Sanger sequencing of bacterial colonies, and sequenced alleles are shown as alignments to the reference sequence, with target site guide RNA sequences aligned below (Benchling). b) Results of Synthego ICE deconvolution of Sanger sequencing traces from PCR products amplified from the downstream *RB1* target site for the indicated cell lines.

```
              1                                                                                82
Consensus     TGGCTGCTGAGGAGAAGCAGGCCCAGTCGCTGCAACCATCCAGCAGCCGCCGCAGCAGCCATTACCCGGCTGCGGTCCAGAG
Whale         TGGCTGCTGAGGAGAAGCAGGCCCAGTCGCTGCAACCATCCAGCAGCCGCCGCAGCAGCCATTACCCGGCTGCTGTCCAGAA
Cow           TGGCTGCTGAGGAGAAGCAGGCCCAGTCGCTGCAACCATCCAGCAGCCGCCGCAGCAGCCATTACCCGGCTGCGGTCCAGAG
Human         TGGCTGCTGAGGAGAAGCAGGCCCAGTCGCTGCAACCATCCAGCAGCCGCCGCAGCAGCCATTACCCGGCTGCGGTCCAGAG
Mouse         TGGCTGCTGAGGAGAAGCAGGCCCAGTCTCTGCAACCATCCAGCAGCCGCCGCAGCAGCCATTACCCGGCTGCGGTCCAGGG
PTEN Exon 1   ----------------------------------------------------------------------------------
Guide Site    ----------------------------------------------------------------------------------

              83                                                                              164
Consensus     CCAAGCGGCAGCAGAGCGAGGGGCATCAGCCACCGCCAAGTCCAGAGCCATTTCCATCCTGCAGAAGAAGCCCCGCCACCAG
Whale         CCAAGCGGCAGCAGAGCGAGGGGCATCAGCCACCGCCAAGTCCAGAGCCATTTCCATCCTGCAGAAGAAGCCCCGCCACCAG
Cow           CCAAGCGGCAGCAGAGCGAGGGGCATCAGCCACCGCCAAGTCCAGAGCCATTTCCATCCTGCAGAAGAAGCCCCGCCACCAG
Human         CCAAGCGGCGGCAGAGCGAGGGGCATCAGCTACCGCCAAGTCCAGAGCCATTTCCATCCTGCAGAAGAAGCCCCGCCACCAG
Mouse         CCAAGCGGCAGCAGAGCGAGGGGCATCAGCGACCGCCAAGTCCAGAGCCATTTCCATCCTGCAGAAGAAGCCTCGCCACCAG
PTEN Exon 1   ----------------------------------------------------------------------------------
Guide Site    ----------------------------------------------------------------------------------

              165                                                                             246
Consensus     CAGCTTCTGCCATCTCTCTCCTCCTTTTTCTTCAGCCACAGGCTCCCAGACATGACAGCCATCATCAAAGAGATCGTTAGCA
Whale         CAGCTTCTGCCATCTCTCTCCTCCTTTTTCTTCAGCCACAGGCTCCCAGACATGACAGCCATCATCAAAGAGATCGTTAGCA
Cow           CAGCTTCTGCCATCTCTCTCCTCCTTTTTCTTCAGCCACAGGCTCCCAGACATGACAGCCATCATCAAAGAGATCGTTAGCA
Human         CAGCTTCTGCCATCTCTCTCCTCCTTTTTCTTCAGCCACAGGCTCCCAGACATGACAGCCATCATCAAAGAGATCGTTAGCA
Mouse         CAGCTTCTGCCATCTCTCTCCTCCTTTTTCTTCAGCCACAGGCTCCCAGACATGACAGCCATCATCAAAGAGATCGTTAGCA
PTEN Exon 1   ---------------------------------------------------ATGACAGCCATCATCAAAGAGATCGTTAGCA
Guide Site    -------------------------------------------------------------TCATCAAAGAGATCGTTAGC-

              247                                                                             328
Consensus     GAAACAAAAGGAGATATCAAGAGGATGGATTCGACTTAGACTTGACCTGTATCCATTTCTGCGGCTGCCGCTCCTCTTTGCC
Whale         GAAACAAAAGGAGATATCAAGAGGATGGATTCGACTTAGACTTGACCTGTATCCATTTCTGTGGCTGCCGCTCCTCTTTGCC
Cow           GAAACAAAAGGAGATATCAAGAGGATGGATTCGACTTAGACTTGACCTGTATCCATTTCTGTGGCTGCTGTTCCTCTTTGCC
Human         GAAACAAAAGGAGATATCAAGAGGATGGATTCGACTTAGACTTGACCTGTATCCATTTCTGCGGCTGC---TCCTCTTTACC
Mouse         GAAACAAAAGGAGATATCAAGAGGATGGATTCGACTTAGACTTGACCTGTATCCATTTCTGCGGCTGT---TCCTCTTTGCT
PTEN Exon 1   GAAACAAAAGGAGATATCAAGAGGATGGATTCGACTTAGACTTGACCT----------------------------------
Guide Site    ----------------------------------------------------------------------------------

              329                                     369
Consensus     TTTCTGTCACTCTCTCTTATAACGTGGGAGTAGACGGATGC
Whale         TTTCTGTCACTCTCTCTTATAACGTGGGAGTAGACGGATGC
Cow           TTTCTGTCACTCTCTCTGATAACGTGGGAATAGACGGATGC
Human         TTTCTGTCACT--CTCTTAGAACGTGGGAGTAGACGGATGC
Mouse         TTTCTGTCA----CTCTGATAACGTGGGAGTAGACGGATGC
PTEN Exon 1   -----------------------------------------
Guide Site    -----------------------------------------
```

**Supplementary Figure 3. Alignment of CRISPR repair fidelity target sequences.** Alignment of NGS amplicon sequences for each species, along with *PTEN* Exon 1, and guide RNA target site, used in comparative DSB repair fidelity experiments.

Amplicon/primer pair was designed based on size appropriate for paired-end sequencing, conservation of internal sequence and primer binding sites, and optimal positioning to detect large deletions (these were observed to occur upstream of the guide site in preliminary Sanger sequencing data using a longer amplicon). Mismatches to consensus sequence are highlighted in red. Alignments are shown in Benchling with Clustal Omega.

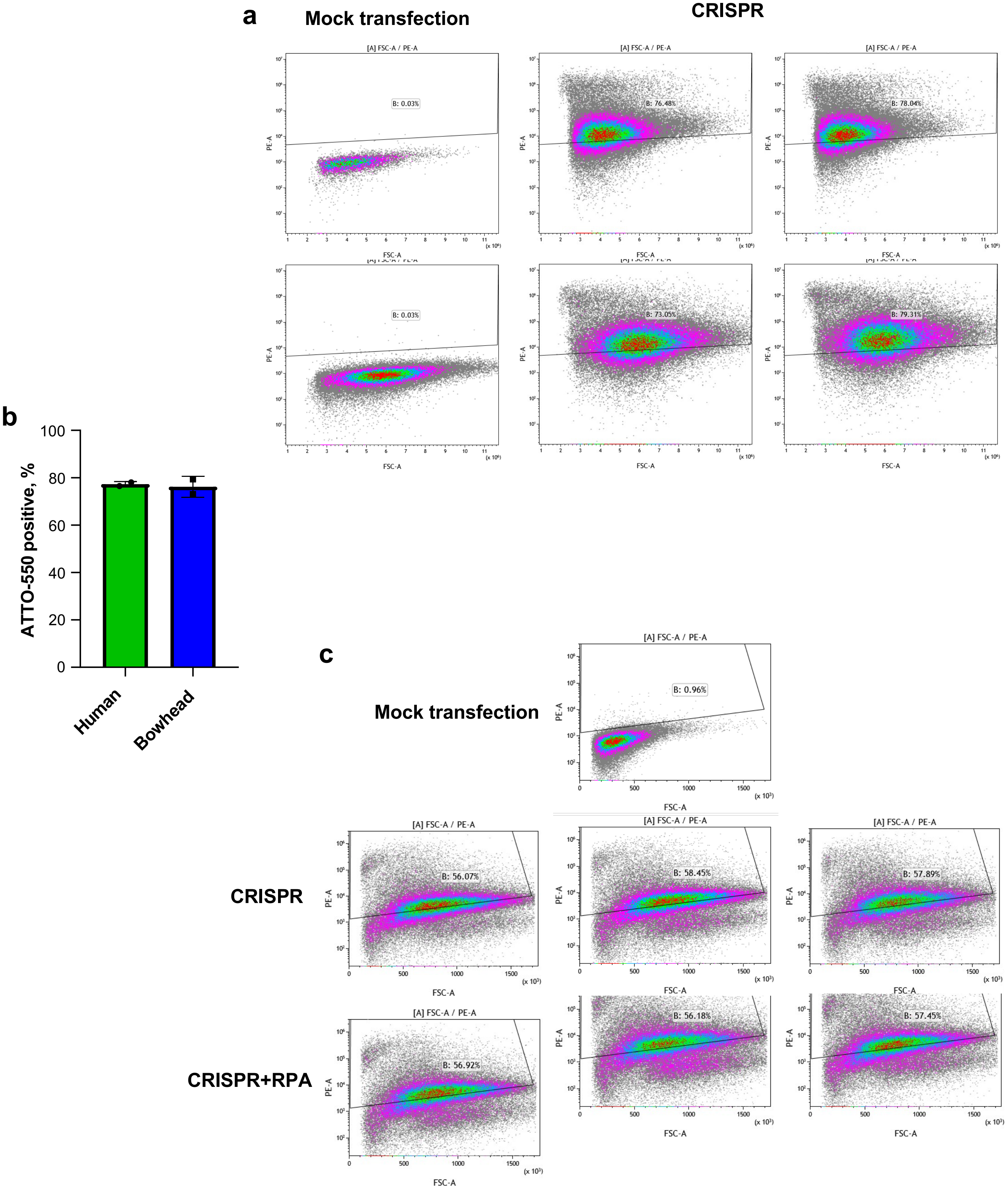
a
Mock transfection
CRISPR
[A] FSC-A / PE-A
B: 0.03%
B: 76.48%
B: 78.04%
B: 0.03%
B: 73.05%
B: 79.31%
PE-A
FSC-A
b
ATTO-550 positive, %
100
80
60
40
20
0
Human
Bowhead
c
Mock transfection
B: 0.96%
CRISPR
B: 56.07%
B: 58.45%
B: 57.89%
CRISPR+RPA
B: 56.92%
B: 56.18%
B: 57.45%

**Supplementary Figure 4. Transfection efficiency of CRISPR RNP with ATTO 550- labeled tracRNA.** a) Flow cytometry measurement of percentage ATTO-550 positive cells immediately (0h) post transfection in human and bowhead primary fibroblasts.
ATTO-550 positive gate was set based on mock-transfected control. Plots in each row show results by species for mock transfection and replicate ATTO-550 labeled RNP transfections. b) Quantification of average percentage ATTO-550 positive cells by species. c) Flow cytometry measurement of percentage ATTO-550 positive cells immediately (0h) post transfection in human primary fibroblasts with or without co- transfection of rhRPA. ATTO-550 positive gate was set based on mock-transfected control. Plots in each row show results of triplicate transfections.

**Mock transfection**

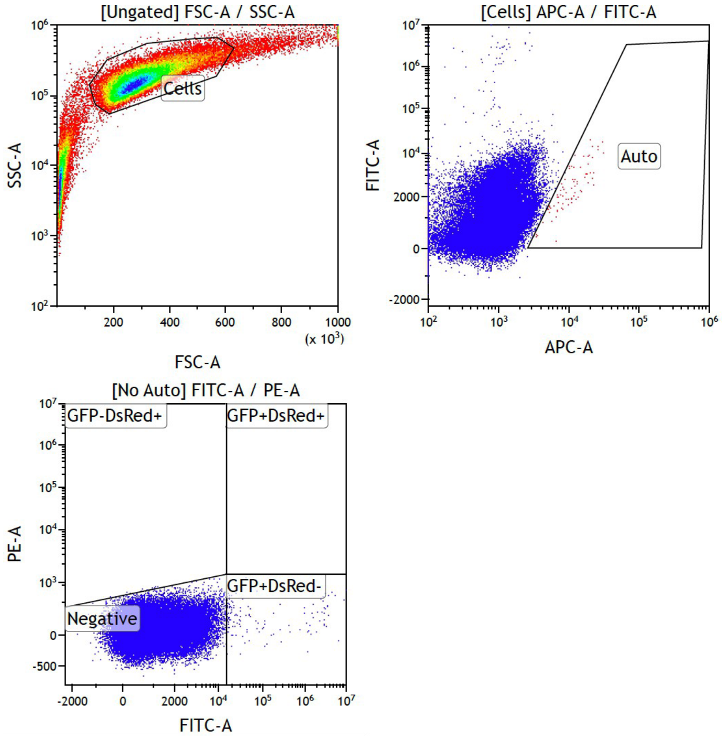

| Gate | Number | %Gated |
|---|---|---|
| All | 51,052 | 100.0 |
| GFP-DsRed+ | 14 | 0.0 |
| GFP+DsRed- | 140 | 0.3 |
| GFP+DsRed+ | 0 | 0.0 |
| Negative | 50,898 | 99.7 |

**I-Sce1/DsRed transfection**

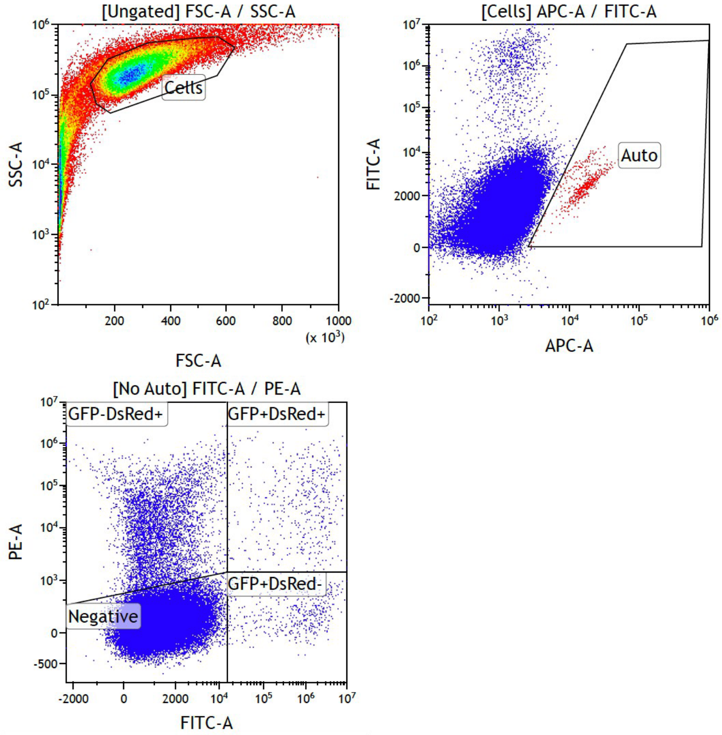

| Gate | Number | %Gated |
|---|---|---|
| All | 52,429 | 100.0 |
| GFP-DsRed+ | 4,840 | 9.2 |
| GFP+DsRed- | 443 | 0.8 |
| GFP+DsRed+ | 485 | 0.9 |
| Negative | 46,661 | 89.0 |

**Supplementary Figure 5. Gating strategy for NHEJ assay.** Mock transfected cells were used to set positive/negative gates that were applied to all samples within a given experiment. The live, singlet cell population was gated based on FSS-SSC and cellular debris was excluded. Autofluorescent cells were gated out based on fluorescence along the APC channel. For calculating NHEJ efficiency, the total numbers of GFP+ and DsRed+ cells were calculated from the FITC-PE plot. Fluorescence compensation was performed in Kaluza.
